# Spreading of a mycobacterial cell surface lipid into host epithelial membranes promotes infectivity

**DOI:** 10.1101/845081

**Authors:** C.J. Cambier, Steven M. Banik, Joseph A. Buonomo, Carolyn R. Bertozzi

## Abstract

Several virulence lipids populate the outer cell wall of pathogenic mycobacteria (Jackson, 2014). Phthiocerol dimycocerosate (PDIM), one of the most abundant outer membrane lipids (Anderson, 1929), plays important roles in both defending against host antimicrobial programs (Camacho et al., 2001; Cox et al., 1999; Murry et al., 2009) and in evading these programs altogether (Cambier et al., 2014a; Rousseau et al., 2004). Immediately following infection, mycobacteria rely on PDIM to evade toll-like receptor (TLR)-dependent recruitment of bactericidal monocytes which can clear infection (Cambier et al., 2014b). To circumvent the limitations in using genetics to understand virulence lipid function, we developed a chemical approach to introduce a clickable, semi-synthetic PDIM onto the cell wall of *Mycobacterium marinum*. Upon infection of zebrafish, we found that PDIM rapidly spreads into host epithelial membranes, and that this spreading inhibits TLR activation. PDIM’s ability to spread into epithelial membranes correlated with its enhanced fluidity afforded by its methyl-branched mycocerosic acids. Additionally, PDIM’s affinity for cholesterol promoted its occupation of epithelial membranes; treatment of zebrafish with statins, cholesterol synthesis inhibitors, decreased spreading and provided protection from infection. This work establishes that interactions between host and pathogen lipids influence mycobacterial infectivity and suggests the use of statins as tuberculosis preventive therapy by inhibiting PDIM spread.

## INTRODUCTION

*Mycobacterium tuberculosis*, the causative pathogen of the pulmonary disease tuberculosis (TB), is estimated to have evolved within the confines of the human lung for millennia (Comas et al., 2013). A result of this co-evolution is a choreographed response of innate and adaptive immune cells culminating in the formation of granulomas, specialized structures that permit bacterial replication and ultimately promote transmission (Ramakrishnan, 2012). A key strategy used by mycobacteria throughout infection is to avoid and manipulate host immune pathways so as to afford the pathogen safe harbor in otherwise bactericidal myeloid cells (Cambier et al., 2014a; Urdahl, 2014).

To better understand these host-pathogen interactions, we have taken advantage of the optically transparent zebrafish larva, a natural host of the pathogen *M. marinum* (Ramakrishnan, 2020; Takaki et al., 2013). Infection of the hindbrain ventricle (HBV), an epithelium-lined cavity, allows for the visualization and characterization of the cellular immune response (Davis and Ramakrishnan, 2009), a response that is comparable to that seen in the mouse lung following infection with *M. tuberculosis* (Srivastava et al., 2014). In both models, mycobacteria are initially phagocytosed by tissue-resident macrophages and are eventually transferred to monocytes which go on to form granulomas (Cambier et al., 2017; Cohen et al., 2018).

In order to reach growth-permissive cells, mycobacteria must first evade prototypical anti-bacterial monocytes. In response to mucosal commensal pathogens, bactericidal monocytes are recruited downstream of TLR signaling (Medzhitov, 2007). Screening of *M. marinum* genetic mutants found that the cell surface lipid phthiocerol dimycocerosate (PDIM) is required to avoid this TLR-dependent immune response (Cambier et al., 2014b). In addition to evading immune detection, PDIM is reported to promote pathogenesis in other ways, such as being required for the relative impermeability of the mycobacterial cell wall (Camacho et al., 2001) and promoting escape from phagolysosomes (Augenstreich et al., 2017; Barczak et al., 2017; Lerner et al., 2017; Quigley et al., 2017). However, the molecular details underlying PDIM’s myriad pathogenic functions remain unknown.

To accomplish mechanistic studies of virulence lipids, we and others developed metabolic labeling strategies where unnatural metabolic precursors are fed to growing bacteria (Siegrist et al., 2015). The unnatural metabolite contains a bioorthogonal functional group that facilitates visualization of macromolecules in living bacteria (Sletten and Bertozzi, 2009). An example is the labeling of trehalose containing lipids with azide-functionalized trehalose (Swarts et al., 2012). Trehalose labeling is so robust that it has since been elaborated on with various functional probes (Kamariza et al., 2018; Rodriguez-Rivera et al., 2017).

Here we have developed comparable chemical tools to monitor PDIM’s distribution during infection, allowing us to better understand PDIM’s mechanism of action. By visualizing PDIM during in vivo infection, we found that the first step in PDIM mediated pathogenesis is to spread into epithelial cells in order to inhibit TLR signaling. Structure function analysis revealed that PDIM’s inherent fluidity, afforded by its methyl-branched fatty acids, promoted occupation of this host lipid niche. PDIM spreading was also dependent on the lipid content of host membranes. Administration of the cholesterol lowering drug, atorvastatin (Lipitor), led to a decrease in PDIM spreading, and subsequent resistance to mycobacterial infection. Our findings provide a mechanistic explanation for the association of statin use with a decrease in TB incidence (Lai et al., 2016a) and support their use as a TB preventative therapy.

## RESULTS

### Controlling Mycomembrane Composition by Chemical Extraction and Reconstitution

Due to the conserved synthesis of lipid intermediates, PDIM lacks unique biosynthetic precursors to facilitate metabolic labeling (Onwueme et al., 2005). However, PDIM is removed following petroleum ether extraction (Moliva et al., 2019), a technique previously used to perform loss of function studies of outer mycomembrane lipids (Silva et al., 1985), as lipids can be removed and added back with petroleum ether (Indrigo, 2003). Using this approach, we hypothesized that we could chemically install a biorthogonal handle onto extracted PDIM and use this modified lipid to elucidate the fundamental mechanisms underlying PDIM’s contribution to virulence. Similar to reports on *M. tuberculosis* and *M. bovis*, we validated that petroleum ether extraction did not affect the growth rate of *M. marinum* in culture (Supplemental Figure 1A) and extracted lipids could be mixed with bacteria in petroleum ether followed by drying to reconstitute these lipids into the mycomembrane (Figure 1A and Supplemental Figure 1B). Similar to reports in macrophages (Indrigo, 2003), following infection of zebrafish (Figure 1B) delipidated bacteria were attenuated for growth and this phenotype was rescued upon reconstitution (Figure 1C and D).

**Figure 1.**
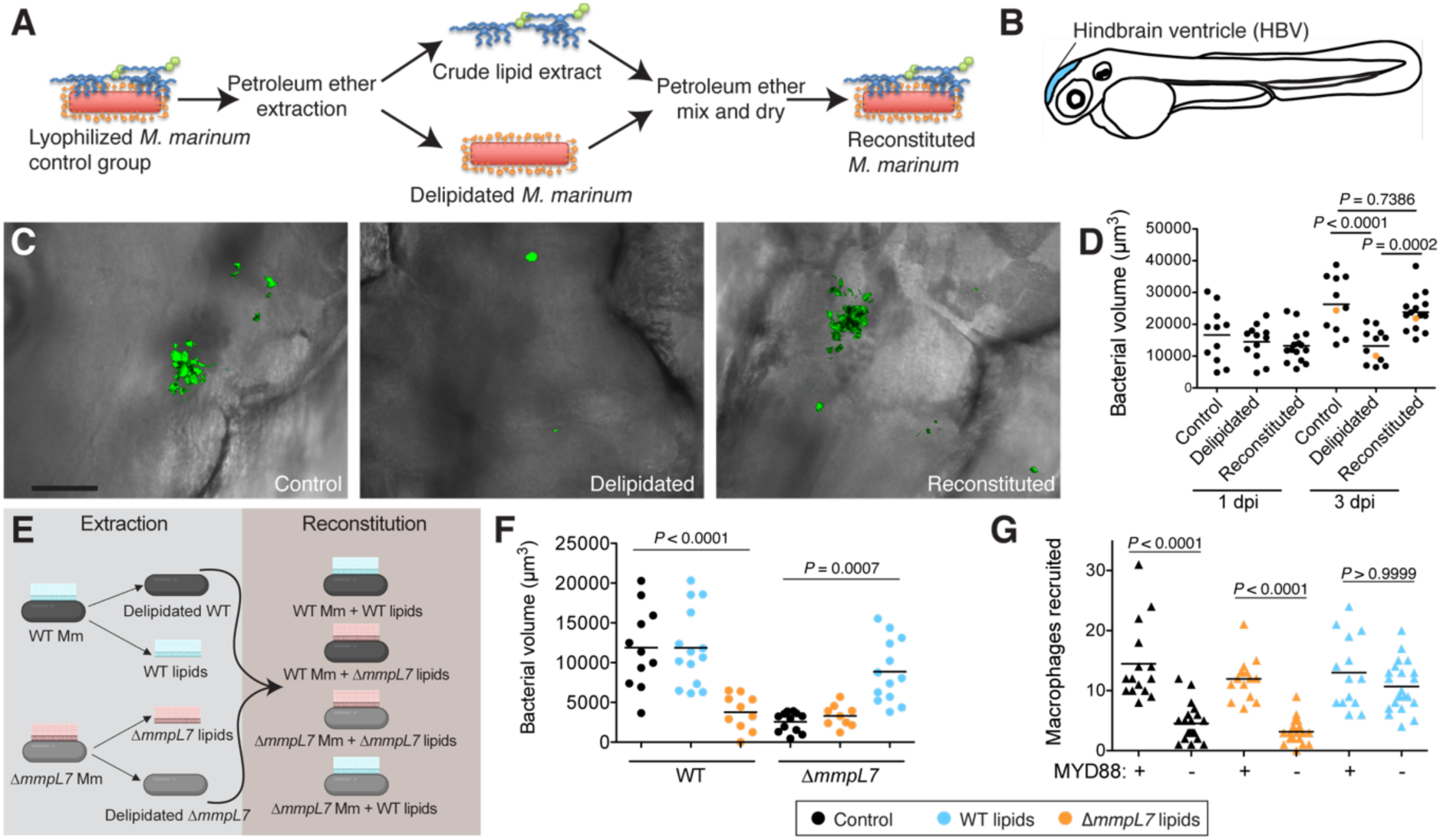
Chemical extraction and reconstitution reveals that pre-infection PDIM reservoirs are required for virulence. **(A)** Model of chemical extraction and reconstitution. **(B)** Model of zebrafish larva showing the hindbrain ventricle (HBV) injection site. **(C)** Representative images of the experiment in **D** (orange dots), scale bar = 50*μ*m. **(D)** Mean bacterial volume after HBV infection of wildtype fish with ∼100 control, delipidated, or reconstituted *M. marinum*. **(E)** Model of lipid swap experiment. **(F)** Mean bacterial volume after HBV infection of wildtype fish with ∼100 WT or Δ*mmpL7 M. marinum* treated as follows: non-extracted control (black), extracted and reconstituted with WT lipids (blue), or extracted and reconstituted with Δ*mmpL7* lipids (orange). **(G)** Mean macrophage recruitment at 3 hpi of the HBV with ∼100 Δ*mmpL7 M. marinum* as treated in **F. (D), (F)**, and **(G)** representative of at least three separate experiments. Ordinary one-way ANOVA with **(D)** Sidak’s multiple comparisons test for the comparison’s shown and **(F)** Tukey’s multiple comparisons test with selected adjusted Pvalues shown. **(G)** Kruskal-Wallis ANOVA for unequal variances with Dunn’s multiple comparisons test with selected adjusted *P* values shown.

### Pre-infection PDIM Reservoirs are Required for Virulence

*M. marinum* mutants in PDIM synthesis (Δ*mas*) and localization to the mycomembrane (Δ*mmpL7*) trigger TLR-dependent immune recognition following infection (Cambier et al., 2014b). TLR signaling leads to the recruitment of activated monocytes that can clear bacteria in an inducible nitric oxide synthase-dependent fashion. PDIM-sufficient wildtype bacteria are not detected by TLRs and instead recruit a comparable number of permissive monocytes downstream of the chemokine CCL2 (Cambier et al., 2014b). However, since Δ*mas* and Δ*mmpL7* lack proteins required for PDIM’s synthesis or export, the associated phenotypes could be attributed to the missing proteins rather than to a lack of PDIM. To directly test if the presence of PDIM on the bacterial surface is responsible for evasion of TLR-mediated immunity, we performed a lipid-swap experiment. Wildtype and Δ*mmpL7 M. marinum* were either untreated (control) or extracted and reconstituted with their native lipids or the lipids from the other strain (Figure 1E). Petroleum ether extraction of wildtype bacteria completely removed both dimycocerosic acid (DIM) containing lipids, PDIM and its metabolic precursor phthiodiolone dimycocerosate (PNDIM, Supplemental Figure 1C) both of which were absent in Δ*mmpL7* extracts (Supplemental Figure 1D). Following infection, wildtype control and wildtype bacteria reconstituted with wildtype lipids grew normally whereas wildtype bacteria reconstituted with Δ*mmpL7* lipids were attenuated for growth (Figure 1F). Conversely, Δ*mmpL7* bacteria were attenuated for growth, as expected, unless they were reconstituted with wildtype lipids, in which case they grew at wildtype bacterial rates (Figure 1F). Using an antisense morpholino to knockdown the TLR-adaptor MYD88 (Bates et al., 2007), we also found that the dependence on TLRs to recruit monocytes to Δ*mmpL7* bacteria was abolished with wildtype lipids (Figure 1G). Taken together these experiments highlight the strengths of this chemical approach. Not only does it recapitulate known phenotypes of PDIM genetic mutants, but it directly links the mutant phenotypes to the absence of PDIM on the mycomembrane. Furthermore, our data suggest that the PDIM present on the surface of the bacterium from the onset of infection is required and sufficient to avoid TLRs, as mutants unable to replenish PDIM on their surfaces become virulent when they are coated with wildtype lipids prior to infection.

### Synthesis of a Clickable, Biologically Active PDIM

Given the pathogenic importance of the pre-infection mycomembrane lipid content, we hypothesized that labeling this pool of PDIM would allow us to observe characteristics underlying its virulence mechanisms. Both PDIM and its biosynthetic precursor PNDIM are present in the outer mycomembrane. The only difference between these lipids is their diol backbones; PDIM has a methyl ether, while PNDIM has a ketone (Siméone et al., 2007). Either lipid alone can promote infection in mice (Siméone et al., 2007), suggesting chemical flexibility at this site with regards to virulence. Therefore, we converted the methyl ether of PDIM to an alkyl halide with trimethylsilyl iodide (Jung and Lyster, 1977). Subsequent addition of sodium azide provided azido-DIM (Figure 2A). Reconstitution of delipidated bacteria with lipids containing azido-DIM, followed by a copper-free click reaction with the cyclooctyne fluorophore DIBO-488 (Figure 2B) resulted in a ∼100-fold increase in fluorescence (Figure 2C). Confocal microscopy revealed the fluorescent signal to be membrane associated (Figure 2D, E), suggesting azido-DIM incorporated into the mycomembrane. Importantly, we found that adding back native DIMs or azido-DIM to DIM-depleted lipids prior to reconstitution (Supplemental Figure 1E) followed by DIBO-488 labeling rescued DIM-depleted bacteria’s growth attenuation (Figure 2F). Thus, with this approach we can generate bacteria with chemically functionalized PDIM that retain their pathogenicity.

**Figure 2.**
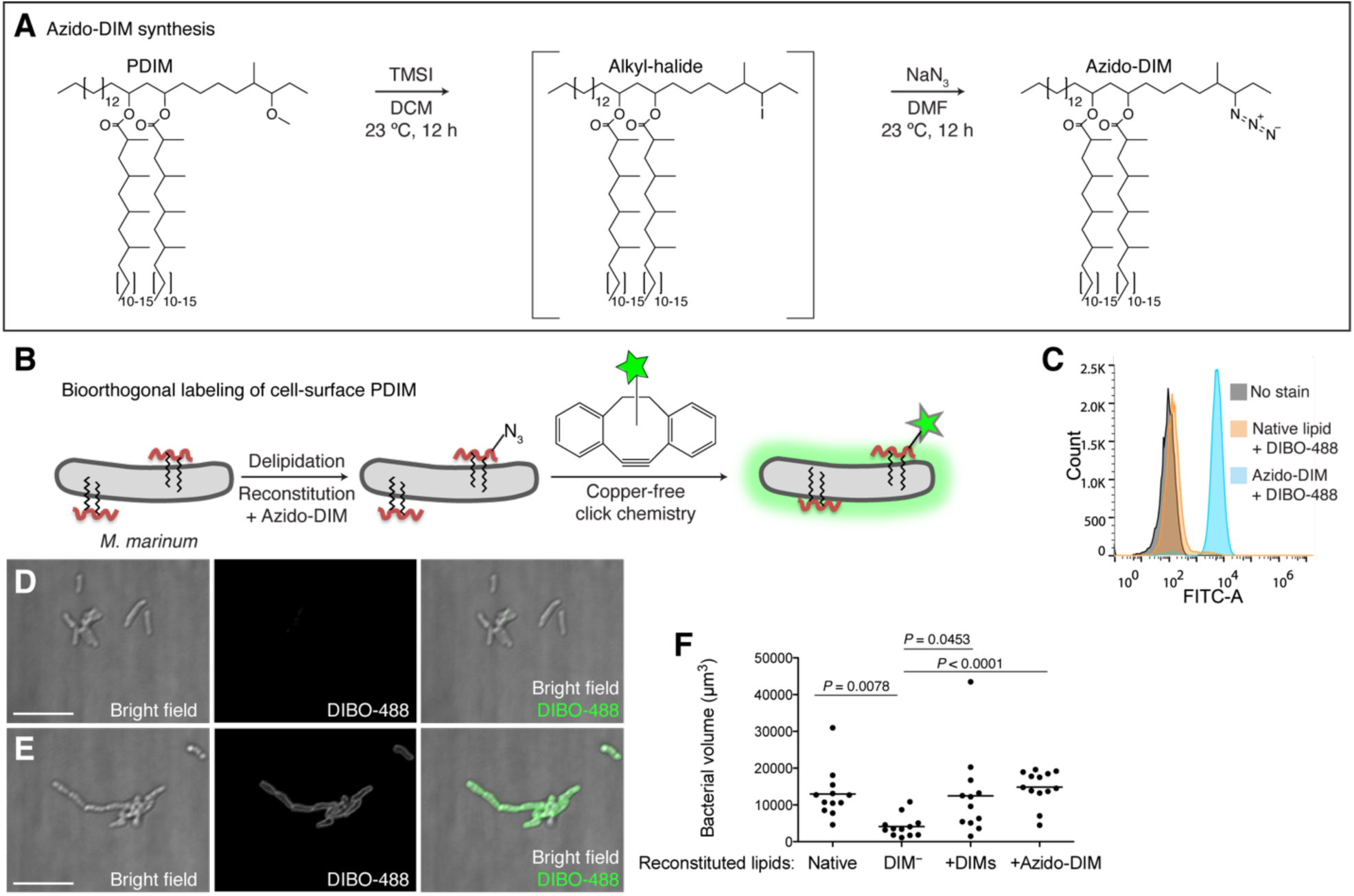
Synthesis and application of a chemically tractable, biologically active PDIM variant, azido-DIM. **(A)** Synthesis of azido-DIM. **(B)** Model of delipidation and reconstitution of bacteria with or without azido-DIM followed by treatment with an azide-reactive cyclooctyne, DIBO-488. **(C)** Flow cytometry analysis of control orazido-DIM reconstituted bacteria treated or untreated with DIBO-488. Image of **(D)** control or **(E)** azido-DIM reconstituted bacteria treated with DIBO-488, scale bar = 8*μ*m. **(F)** Mean bacterial volume 3 days following HBV infection of wildtype fish with ∼100 delipidated *M. marinum* reconstituted with Native, DIM depleted (DIM^-^), DIM^-^ plus native DIMs (+DIMs), or DIM^-^ plus azido-DIM (+Azido-DIM) lipids. Kruskal-Wallis ANOVA for unequal variances with Dunn’s multiple comparisons test with selected adjusted Pvalues shown. **(C)-(F)** representative of three separate experiments.

### PDIM Spreads into Host Cell Membranes In Vivo

To visualize PDIM’s distribution, we infected zebrafish with blue-fluorescent *M. marinum* that were reconstituted with azido-DIM followed by labeling with DIBO-488 (DIM-488). Immediately following phagocytosis, DIM-488 appeared to spread away from bacteria into host membranes (Supplemental Figure 2A). Real-time imaging revealed that the spreading was dynamic in nature, with DIM-488 deposits moving relative to host cells (Supplemental Movie 1). To better visualize spreading of PDIM, we used the transgenic zebrafish line *Tg(mfap4:tdTomato)* whose macrophages express the fluorescent protein tdTomato (Walton et al., 2015). As early as 3 hours post-infection (hpi), DIM-488 had spread into infected macrophage membranes, at both phagosome (Figure 3A, arrows) and more distal membrane sites (Figure 3A, arrow heads). Spreading increased across macrophage membranes by 3 days post-infection (dpi) (Figure 3B). Similar spreading was seen when azido-DIM was conjugated to DIBO-647 (Supplemental Figure 2B, C), suggesting that the lipid, and not the fluorescent probe, was responsible for this phenotype. To quantify the extent of PDIM spreading, we imaged the entire HBV infection site and calculated the proportion of fluorophore labelled azido-DIM that no longer localized with bacteria (Supplemental Figure 2D). Using this number as a proxy for lipid spread, we saw an increase in spreading as infection progressed (Figure 3C). Spreading also occurred following infection of THP-1 macrophages in culture (Supplemental Figure 2E).

**Figure 3.**
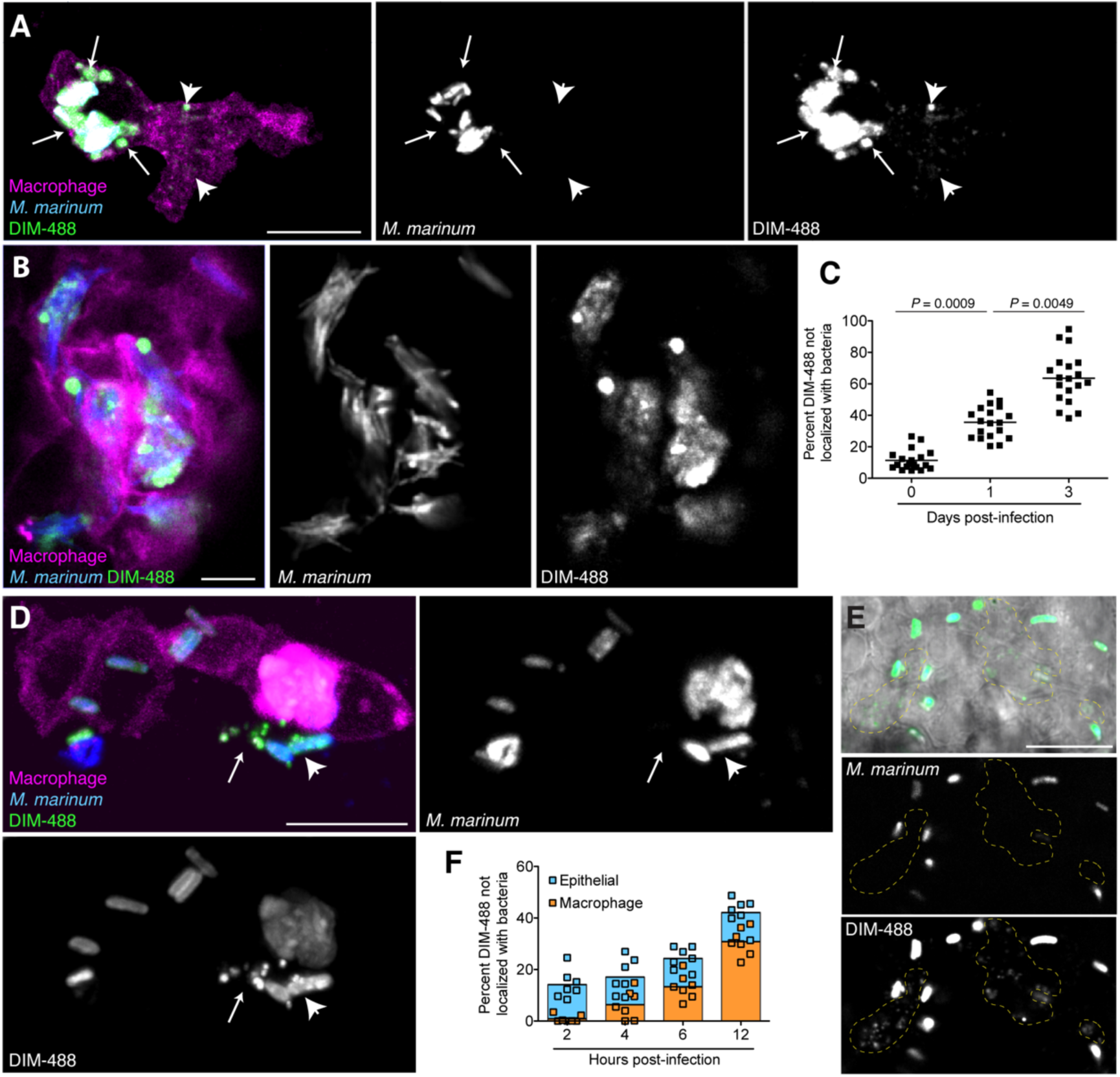
PDIM spreads into host cell membranes. Images of *M. marinum* expressing a cytosolic blue fluorescent protein reconstituted with DIBO-488 labeled azido-DIM (DIM-488) at **(A)** 3 hpi and **(B)** 3 dpi of ∼100 *M. marinum* in the HBV of transgenic fish whose macrophages express a fluorescent protein. Scale bar = 10 *μ*m. Arrows, DIM-488 spread in vicinity of phagosome, arrowheads, DIM-488 spread throughout macrophage. **(C)** Mean percent DIM-488 not localized with bacteria following HBV infection of wildtype fish with ∼100 *M. marinum*. Kruskal-Wallis ANOVA for unequal variances with Dunn’s multiple comparisons test with selected adjusted P values shown. Representative of at least three separate experiments. **(D)** Image highlighting DIM-488 spread from bacteria (arrowhead) to epithelial cells (arrows) at 3 hpi of ∼100 *M. marinum* in the HBV, colors correspond to features defined in A, scale bar = 10*μ*m. **(E)** Image highlighting DIM-488 spread onto epithelial surfaces (yellow-dashed outline) at 2 hpi of ∼100 *M. marinum* in the HBV, scale bar = 10*μ*m. **(F)** Mean percent DIM-488 in macrophage or epithelial cells not localized with bacteria following HBV infection with ∼100 *M. marinum*. Representative of two separate experiments.

These results were consistent with a recent report using cultured macrophages showing that *M. tuberculosis* PDIM occupies their cell membranes (Augenstreich et al., 2019). Nevertheless, we wanted to determine whether spreading was a true attribute of our labeled PDIM or if it was an artifactual property due to our reconstitution method. To address this, we used an established pan-glycolipid labeling method previously used to track the spread of mycobacterial glycolipids into macrophage membranes (Beatty et al., 2000). Control and reconstituted *M. marinum* were treated with periodate and then reacted with a fluorescent hydroxylamine prior to infection (Beatty et al., 2000). We found that spreading of the total pool of fluorophore-labeled glycolipids was equal between control and reconstituted bacteria (Supplemental Figure 3A and B). Thus, reconstitution does not appreciably influence the spreading dynamics of mycomembrane lipids. These data demonstrate that the introduction of a chemically functionalized PDIM into the mycomembrane provides physiologically relevant information regarding PDIM’s host distribution during in vivo infection.

We next wanted to understand how PDIM spreading might be promoting virulence. While it is appreciated that mycobacterial lipids spread throughout macrophage membranes (Beatty et al., 2000), the inability to track specific lipids in an in vivo setting has hampered studies on the pathophysiological relevance of these observations. PDIM has been suggested to interact with the protein substrates of the type VII secretion system ESX-1, including EsxA (Barczak et al., 2017). PDIM and EsxA are both required for cytosolic escape from phagolysosomes (Osman et al., 2020; Quigley et al., 2017; van der Wel et al., 2007), where PDIM is suggested to enhance the pore forming activity of EsxA through its ability to infiltrate macrophage membranes (Augenstreich et al., 2017). Thus, we hypothesized that PDIM localization would be dependent on EsxA secretion. Region of difference-1 *M. marinum* mutants (ΔRD1) which lack EsxA (Volkman et al., 2010), were reconstituted with DIM-488 prior to zebrafish infection. There was no difference in DIM-488 spreading kinetics between wildtype and ΔRD1 *M. marinum* (Supplemental Figure 3C), suggesting that EsxA does not influence PDIM spreading. Besides playing a role in cytosolic escape, PDIM spreading into macrophage membranes has also been shown to promote phagocytosis of extracellular bacteria (Astarie-Dequeker et al., 2009; Augenstreich et al., 2019). To test this in vivo, we measured the rate of phagocytosis of wildtype or Δ*mmpL7 M. marinum* following HBV infection by measuring the number of discrete bacterial objects over time. As bacteria are phagocytosed, individual bacteria can no longer be discerned by confocal microscopy and the number of discrete objects decreases. There was no measurable difference in the rate of phagocytosis of wildtype or Δ*mmpL7* bacteria (Supplemental Figure 3D).

Given the discrepancy regarding PDIM’s role in promoting phagocytosis between the cultured macrophage and zebrafish models, we wondered if the activation state of responding monocyte/macrophage populations in zebrafish larvae was influencing their phagocytic capacities. We know that PDIM-deficient bacteria recruit antimicrobial monocytes downstream of TLR signaling, whereas wildtype bacteria recruit permissive monocytes downstream of CCL2/CCR2 signaling (Cambier et al., 2014b). We also know that wildtype bacteria depend on resident macrophages to produce CCL2 in order to recruit CCR2-expressing monocytes, whereas PDIM-deficient bacteria recruit monocytes independent of resident macrophages (Cambier et al., 2017). Therefore, we wondered if PDIM plays a critical role in evading TLR detection in non-macrophage cell populations prior to any of its documented roles in directly modulating macrophage biology. Upon closer examination, we observed DIM-488 deposits on zebrafish epithelium at 24 hpi (Supplemental Figure 3E, arrows). Imaging at 3 hpi we captured extracellular bacteria having spread DIM-488 in the vicinity of an infected macrophage (Figure 3D). This ability to spread into epithelial membranes was confirmed in human A549 epithelial cells in vitro (Supplemental Figure 3F). To address the timing of PDIM spread into epithelial and macrophage membranes we imaged infected zebrafish at 2 hpi, prior to macrophage phagocytosis, and at 4, 6, and 12 hpi by which time the majority of bacteria are phagocytosed. Preceding any appreciable spreading onto macrophages, DIM-488 had already spread onto epithelial cells by 2 hpi (Figure 3E yellow outlines and 3F). These data suggest that mycobacteria spread PDIM onto epithelial cells prior to interactions with macrophages. To confirm this observation, we depleted macrophages from zebrafish larva using clodronate-loaded liposomes (Bernut et al., 2014). DIM-488 spreading still occurred in the absence of macrophages (Supplemental Figure 3G). These data demonstrate that PDIM spreads onto epithelial cells independent of and prior to macrophage phagocytosis.

### PDIM’s Fluidity Promotes Spread into Epithelial Cell Membranes

We sought to understand PDIM’s properties that facilitated its ability to spread into host membranes. It is well established that increased membrane fluidity promotes membrane mixing (Howell et al., 1972), leading us to ask if PDIM occupied membrane domains of high fluidity. Previous studies from our lab found that the mycomembrane of *Mycobacterium smegmatis* is relatively immobile by using fluorescence recovery after photobleaching (FRAP) of metabolically labeled trehalose monomycolate (TMM) (Rodriguez-Rivera et al., 2017). We confirmed this for *M. marinum* TMM. Using the metabolic label 6-azido-trehalose (Swarts et al., 2012) followed by reaction with DIBO-488 to track TMM (TMM-488), we found very little recovery following photobleaching of TMM-488, with only around 40% of the labeled lipids being mobile (Figure 4A–C, and Supplemental Figure 4A). When we assessed PDIM, we found DIM-488 recovery to be very efficient, with a half-life of 3 seconds and around 95% of the signal being mobile (Figure 4A–C, and Supplemental Figure 4B). Thus, PDIM-containing mycomembrane domains have a uniquely high fluidity.

**Figure 4.**
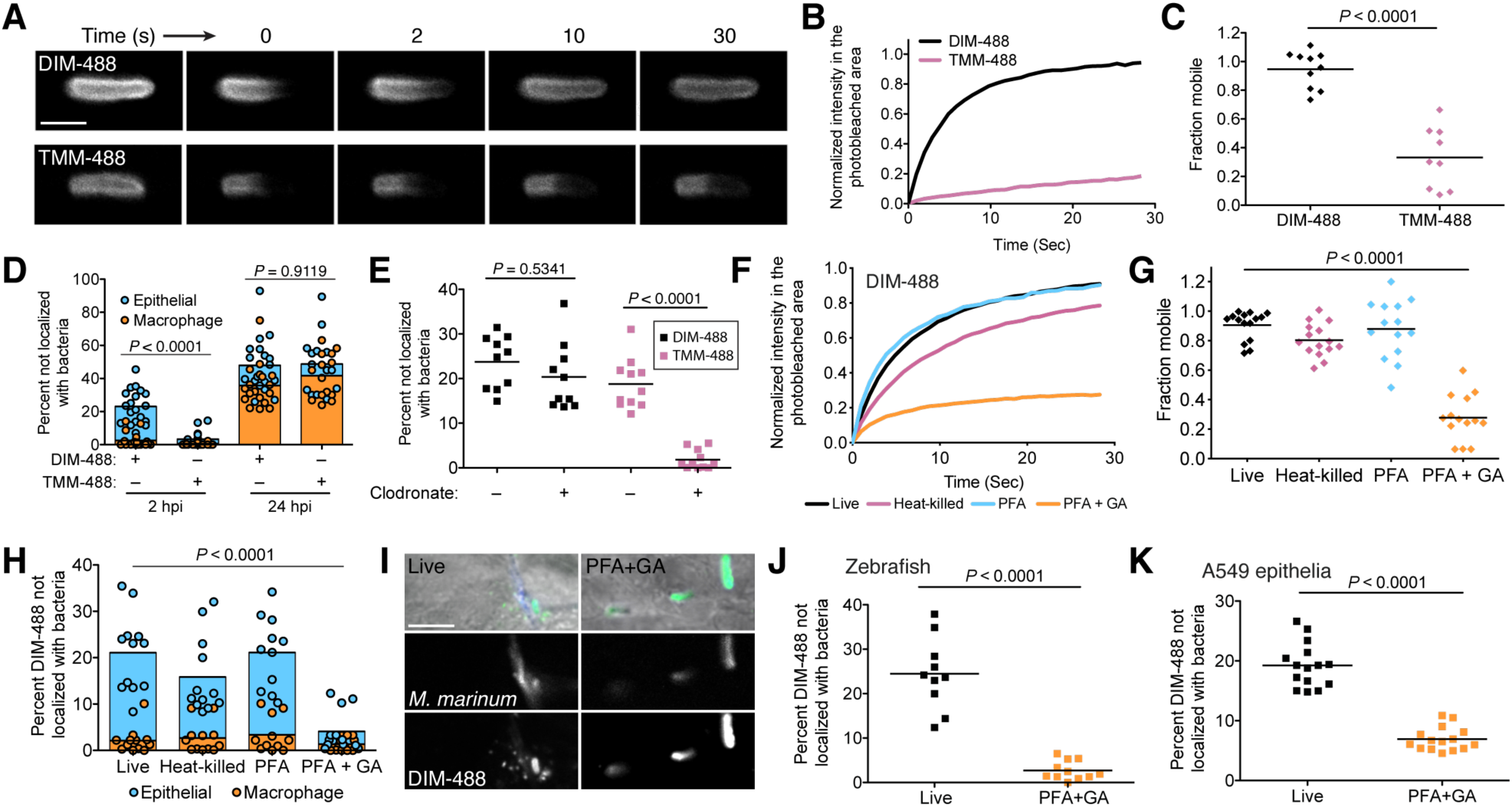
PDIM’s mobility promotes spread into epithelial cells. **(A)** Representative FRAP images of DIM-488 and TMM-488 labeled *M. marinum*, scale bar = 2*μ*m. **(B)** Fluorescent recovery curves after photobleaching of DIM-488 or TMM-488 labeled M. marinum, lines represent the average signal from n = 10 cells. **(C)** Mean fraction mobile following fitting of FRAP curves to data generated in **B. (D)** Mean percent DIM-488 or TMM-488 in macrophage or epithelial cells not localized with bacteria following HBV infection with ∼100 *M. marinum*, two-tailed Mann whitney test. **(E)** Mean percent DIM-488 or TMM-488 not localized with bacteria 24 h following HBV infection of lipo-PBS or lipo-clodronate treated fish with −100 *M. marinum*. **(F)** Fluorescent recovery curves after photobleaching of live, heat-killed, 4% paraformaldehyde (PFA) fixed, or 4% paraformaldehyde plus 1% glutaraldehyde (PFA+GA) fixed DIM-488 labeled *M. marinum*, lines represent the average signal from n = 14-15 cells. **(G)** Mean fraction mobile following fitting of FRAP curves to data generated in **F. (H)** Mean percent DIM-488 in macrophage or epithelial cells not localized with bacteria 2 h following HBV infection with −100 M. marinum treated as in **F. (I)** Images of live or PFA+GA treated DIM-488 labeled *M. marinum* at 2 hpi of the HBV with −100 bacteria, scale bar = 5*μ*m. Mean percent DIM-488 not localized with bacteria 24 h following **(J)** infection of lipo-clodronate treated fish or (K) A549 epithelial cells with live or PFA+GA fixed DIM-488 labeled M. marinum. **(C), (J)**, and **(K)** two-tailed, unpaired t test. **(E), (G)**, and **(H)** ordinary one-way ANOVA with Tukey’s multiple comparisons test with selected adjusted P values shown. **(B)-(H)** and **(J)-(K)** representative of three separate experiments.

If high membrane fluidity facilitates membrane mixing, then TMM should not be able to spread into host cells as efficiently as PDIM does. Indeed, we found that TMM-488 failed to spread onto epithelial cells at 2 hpi. Only when phagocytosed within macrophages at 24 hpi was TMM spreading detected (Figure 4D and Supplemental Figure 4C). Even in the absence of macrophages, where bacteria have a prolonged contact time with epithelial cells, TMM-488 spreading was negligible (Figure 4E). Thus, mycomembrane fluidity appears to correlate with the ability to spread into host epithelial membranes.

To determine if there is a causal relationship between fluidity and spreading, we used FRAP to identify conditions that decreased PDIM’s recovery after photobleaching. Neither heat-killing nor mild chemical fixation (4% paraformaldehyde (PFA)) significantly reduced DIM-488’s recovery (Figure 4F, G and Supplemental Figure 5). However, fixation with 4% PFA + 1% glutaraldehyde (GA), a more effective fixative for membrane-associated proteins (Huebinger et al., 2018), resulted in an almost 80% reduction in the mobile fraction of DIM-488 (Figure 4F, G and Supplemental Figure 5). PFA+GA treatment decreased DIM-488 spread onto epithelial cells at 2 hpi (Figure 4H and 4I). Likewise, fixed DIM-488 did not spread into epithelial membranes following a 24 h infection of macrophage-depleted zebrafish (Figure 4J). Nor did it spread onto A549 epithelial cells after 24 h (Figure 4K). Together these results suggest that PDIM’s fluidity promotes spreading into epithelial membranes.

**Figure 5.**
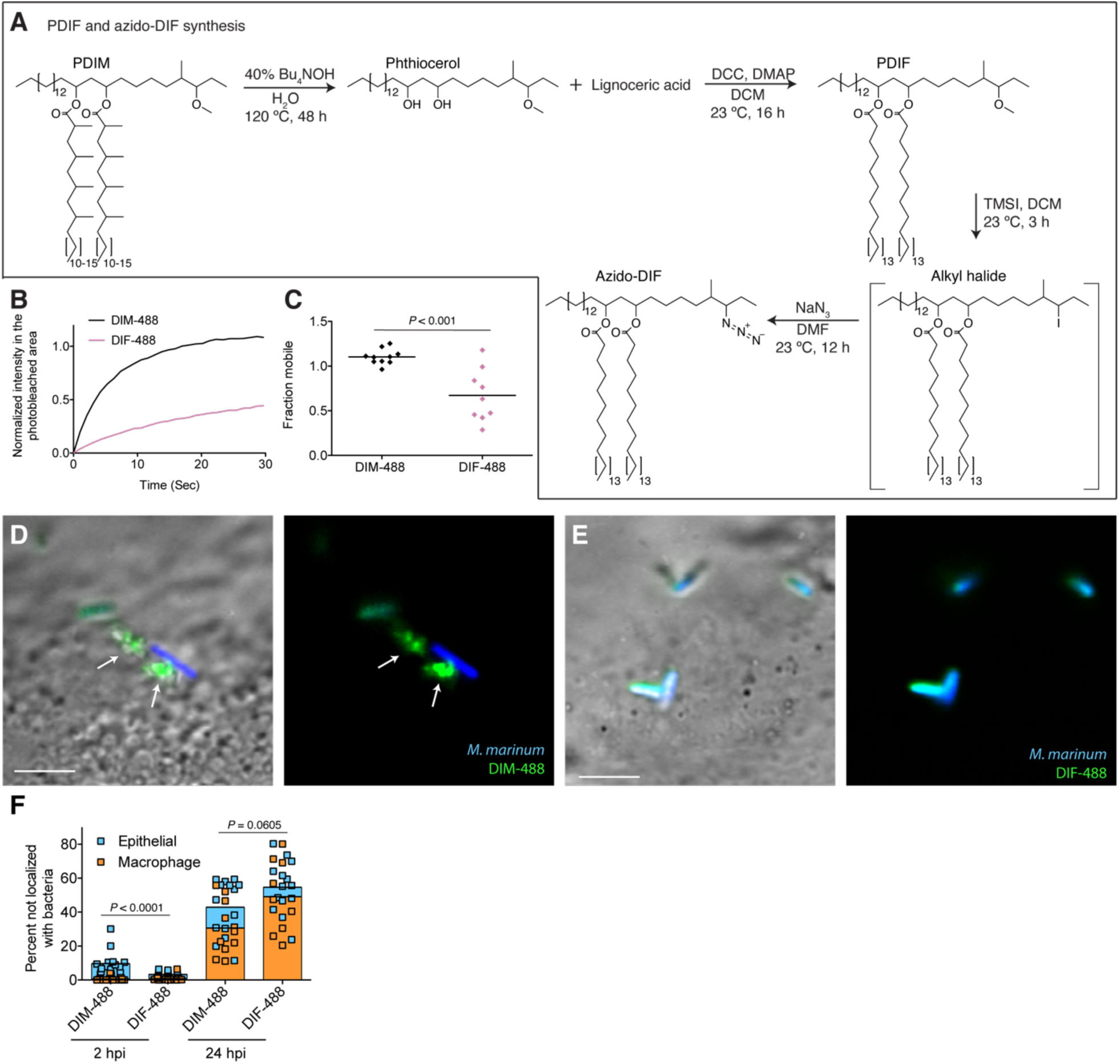
PDIM’s methyl-branched mycocerosic acids are required for virulence. **(A)** Phthiocerol di-fatty acid (PDIF) and azido-DIF synthesis. **(B)** Fluorescent recovery curves after photobleaching of DIM-488 or DIF-488 labeled *M. marinum*, lines represent the average signal from n = 9-10 cells. **(C)** Fraction mobile following fitting of FRAP curves to data generated in **B**. Two-tailed, unpaired t test. Images of *M. marinum* expressing a blue fluorescent protein reconstituted with **(D)** DIM-488 or **(E)** DIF-488 at 2 hpi into the HBV of wildtype fish, arrows indicated spread signal, scale bar = 5*μ*m. **(F)** Mean percent DIM-488 or DIF-488 in macrophage or epithelial cells no longer localized with bacteria following HBV infection with ∼100 *M. marinum*. Two-tailed Mann Whitney test for 2 hpi and two-tailed, unpaired t test for 24 hpi. **(B), (C)**, and **(F)** Representative of three separate experiments.

### PDIM’s Methyl-Branched Mycocerosic Acids Promote Fluidity and Spreading

To further test if PDIM’s high membrane fluidity is responsible for its spreading, we sought to take advantage of our ability to alter PDIM’s structure. Lipid structure is known to modulate membrane fluidity (Los and Murata, 2004). Saturated lipids form closely packed assemblies giving rise to more rigid bilayers, while unsaturated lipids do not pack as tightly and produce more fluid bilayers. Methyl branches on otherwise saturated lipids have also been shown to increase membrane fluidity (Budin et al., 2018; Poger et al., 2014). Therefore, we hypothesized that PDIM’s methyl-branched mycocerosic acids determine its fluidity. To test this, we hydrolyzed PDIM with tetrabutylammonium hydroxide to isolate phthiocerol. We then esterified phthiocerol with the saturated fatty acid lignoceric acid that resulted in straight chain lipids of comparable length to mycocerosic acid. This phthiocerol di-fatty acid (PDIF) was then treated similarly to PDIM (Figure 2A) to yield the azide-labeled PDIF, azido-DIF (Figure 5A). FRAP studies confirmed that DIBO-488 labeled azido-DIF (DIF-488) was not as mobile as DIM-488, with only ∼50% in the mobile fraction (Figure 5B, C and Supplemental Figure 6A–C). Upon zebrafish infection, DIF-488 failed to spread onto epithelial cells at 2 hpi (Figure 5D–F). However, similar to other mycobacterial lipids (Beatty et al., 2000) including TMM (Figure 4A), DIF-488 was capable of spreading into macrophage membranes at later timepoints (Figure 5F and Supplemental Figure 6D, E). Thus, PDIM’s high fluidity mediated by its methyl-branched lipid tails is the reason for its unique ability to spread into epithelial membranes.

**Figure 6.**
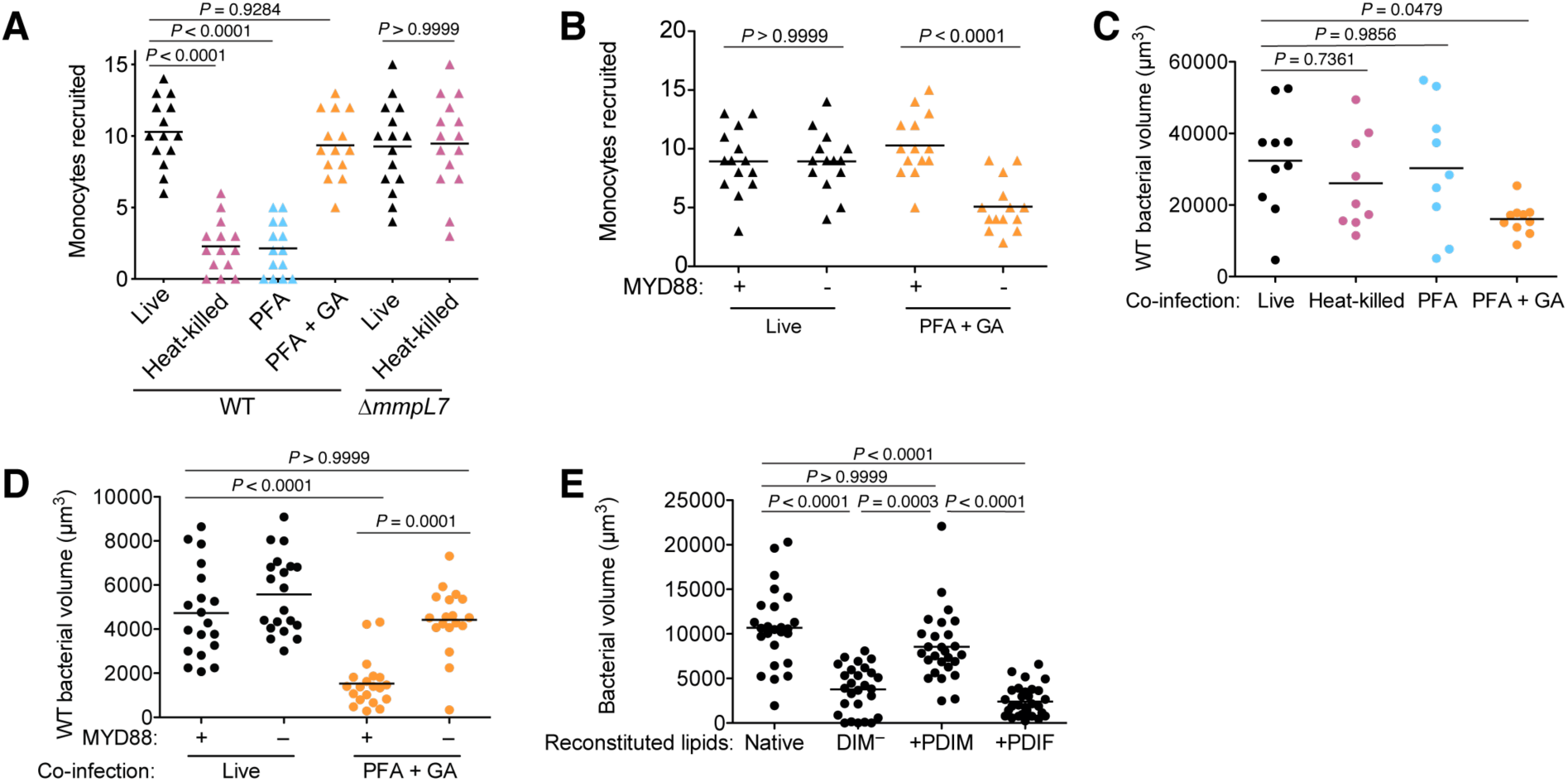
PDIM spreading into epithelial membranes is required to evade TLRs. **(A)** Mean monocyte recruitment at 3 hpi of the HBV with ∼100 live, heat-killed, PFA treated, or PFA+GA treated wildtype (WT) or∼100 live or heat-killed AmmpL7 *M. marinum*. **(B)** Mean monocyte recruitment at 3 hpi of the HBV of wildtype or MYD88-depleted fish with −100 live or PFA+GA treated wildtype *M. marinum*. Mean volume of wildtype *M. marinum* following co-infection with **(C)** wildtype *M. marinum* treated as in A or **(D)** live or PFA+GA treated wildtype M. marinum in wildtype or MYD88-depleted fish. **(E)** Mean bacterial volume 3 days following HBV infection of wildtype fish with ∼100 *M. marinum* reconstituted with Native, DIM-depleted (DIM^-^), DIM^-^ plus PDIM (+PDIM), or DIM^-^ plus PDIF (+PDIF) lipids. **(A)-(C)** ordinary one-way ANOVA with Tukey’s multiple comparisons test with selected adjusted *P* values shown. **(D)** and **(E)** Kruskal-Wallis ANOVA for unequal variances with Dunn’s multiple comparisons test with selected adjusted *P* values shown. **(A)-(E)** representative of three separate experiments.

### PDIM Spreading into Epithelial Membranes is Required to Evade TLRs

Is PDIM spreading into epithelial membranes required to inhibit TLR signaling? We first evaluated monocyte recruitment towards infecting mycobacteria to address this question. Wildtype *M. marinum* must be alive in order to stimulate resident macrophages to express CCL2 to recruit permissive monocytes as heat-killed bacteria failed to do so (Cambier et al., 2017). In contrast, PDIM-deficient *M. marinum* do not depend on resident macrophages to recruit bactericidal monocytes and do so in a manner independent of bacterial viability (Cambier et al., 2017). We now wondered if, in order for mycobacteria to gain access to resident macrophages, PDIM must first spread into epithelial membranes to prevent recruitment of bactericidal monocytes. We confirmed that heat-killed mycobacteria do not recruit monocytes (Figure 6A). Likewise, PFA-treated bacteria which can also spread PDIM did not recruit monocytes (Figure 6A). However, monocyte recruitment towards PFA+GA treated bacteria, which do not spread PDIM, phenocopied heat-killed Δ*mmpL7 M. marinum*; even though they are not viable, they recruited monocytes to a similar extent as live bacteria (Figure 6A). Moreover, this monocyte recruitment was now downstream of TLR signaling, as knockdown of MYD88 prevented monocyte recruitment towards PFA+GA treated bacteria (Figure 6B). Taken together these data demonstrate that PDIM spreading into epithelial membranes is required to prevent TLR-dependent recruitment of monocytes.

TLR signaling has been shown to recruit microbicidal monocytes that kill infecting bacteria (Cambier et al., 2014b). Not only are PDIM-deficient bacteria killed but co-infected PDIM-expressing bacteria are as well (Cambier et al., 2014b). This latter finding allowed us to directly examine the link between PDIM spreading and bacterial killing by resultant TLR-signaled monocytes. We infected larvae with live red-fluorescent wildtype *M. marinum* along with green-fluorescent wildtype *M. marinum* that were live (untreated), heat-killed, PFA treated, or PFA+GA treated and then determined the burdens of the red fluorescent wildtype *M. marinum* at 3 dpi. Only co-infection with PFA+GA treated bacteria caused attenuation of wildtype bacteria (Figure 6C). Moreover, this attenuation was dependent on MYD88 (Figure 6D). Lastly, we asked if PDIF, which is unable to spread into epithelial cells (Figure 4F), could rescue DIM-deficient bacterial growth similarly to PDIM (Figure 2F). Consistent with the requirement for PDIM to spread into epithelial cells to promote virulence, we found that reconstitution with PDIF does not rescue bacterial growth (Figure 6E). Together these findings implicate PDIM spreading into epithelial membranes in inhibiting TLR signaling and the resultant recruitment of microbicidal monocytes that can kill infecting mycobacteria.

### Host Cholesterol Promotes PDIM Spread and Mycobacterial Infectivity

We next asked if host lipids also influence PDIM spreading. Cholesterol has been reported to modulate macrophage interactions with mycobacteria (Gatfield and Pieters, 2000). However, three lines of evidence suggest a macrophage-independent relationship between mycobacteria and cholesterol: 1) 36% of the lipid extracted from *Mycobacterium bovis* harvested from necrotic mouse lung (where macrophages are sparse) was found to be host cholesterol (Kondo and Kanai, 1976); 2) mycobacterial-associated cholesterol was isolated as a mixture with PDIM (Kondo and Kanai, 1976); 3) *M. tuberculosis* grown in culture can sequester cholesterol to its outer mycomembrane, a process dependent on mycomembrane lipids (Brzostek et al., 2009). To test if host cholesterol interacted with PDIM to facilitate its spreading, we first grew wildtype and Δ*mmpL7 M. marinum* in the presence of alkyne-cholesterol (Supplemental Figure 7A) for 48 h, followed by treatment with azide-conjugated alexafluor-647. We found that only wildtype *M. marinum* could sequester cholesterol to their mycomembrane (Figure 7A–C). We next tested whether cholesterol contributes to spreading of PDIM into host membranes. We depleted cholesterol from A549 epithelial cells with methyl β–cyclodextrin (MßCD) which resulted in an eight-fold decrease in cholesterol (Supplemental Figure 7B). This was associated with decreased DIM-488 spreading (Supplemental Figure 7C). Importantly, in MßCD treated cells we could restore cholesterol to untreated levels with water-soluble cholesterol, and this restored DIM-488 spreading (Supplemental Figure 7B, C). To test if host cholesterol facilitates PDIM spreading in vivo, we treated zebrafish with statins, drugs that inhibit HMG-CoA reductase, the rate-limiting step of cholesterol biosynthesis. Specifically, we used atorvastatin which was shown to lower cholesterol in zebrafish (Maerz et al., 2019). Following 24 h of treatment, atorvastatin decreased cholesterol levels in the larvae by 25%, and co-treatment with water-soluble cholesterol restored cholesterol levels (Figure 7D). Atorvastatin treatment also decreased DIM-488 spreading onto epithelial cells (Figure 7E and F), suggesting that PDIM’s interaction with cholesterol promotes spreading into epithelial membranes in vivo.

**Figure 7.**
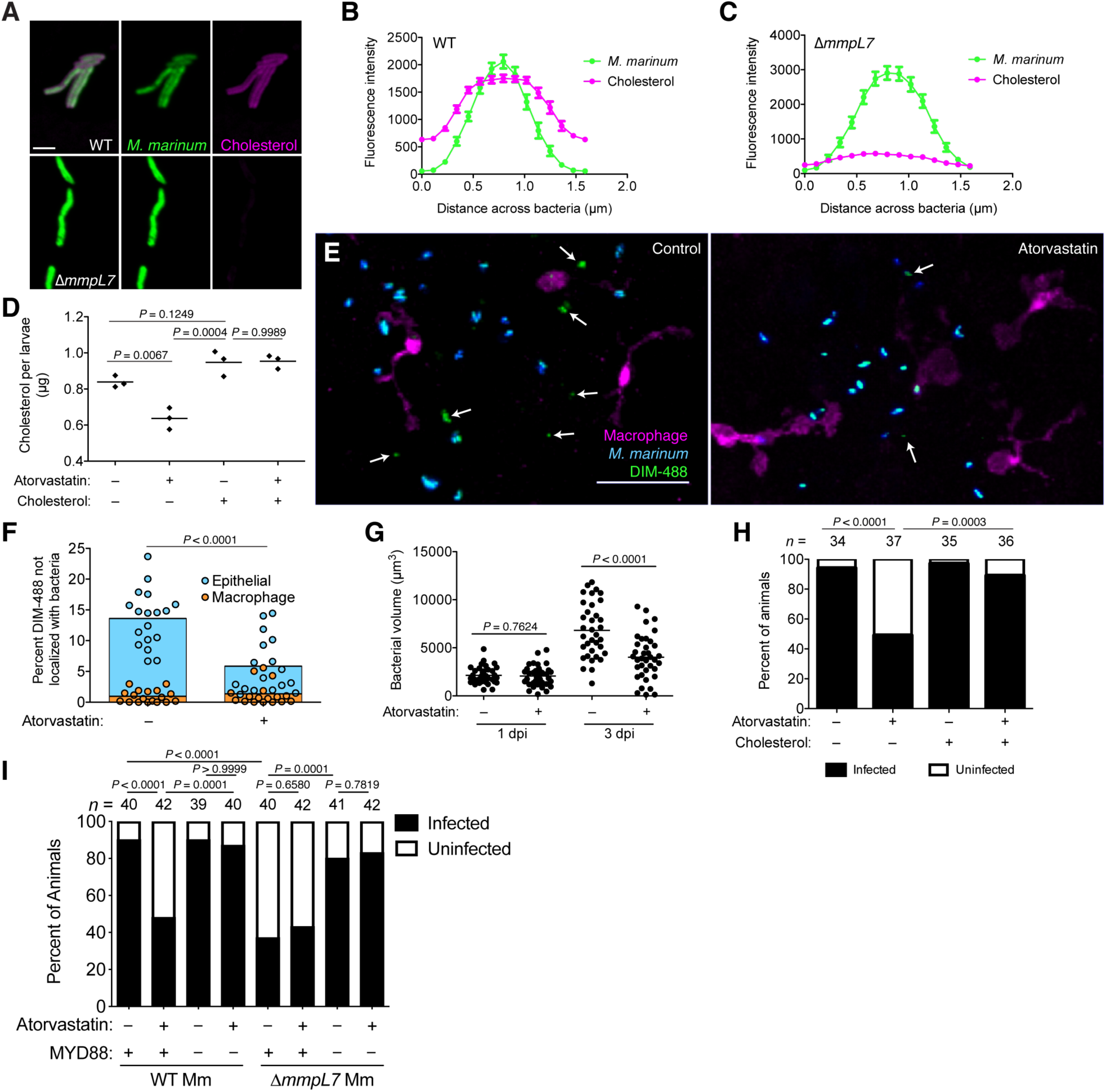
Cholesterol promotes PDIM spreading and mycobacterial infectivity **(A)** Images of vnidtype (WT) and Δ*mmpL7 M. marinum* expressing a green fluorescent protein following 48 h incubation with alkyne-choies-terof reacted with AiexaFfour647 Azide, scale bar = 3*μ*m Mean (±SEM) fluorescent intensity of line profiles drawn perpendicular to poles of **(B)** WT and **(C)** Δ*mmpL7 M. marinum* labeled as in A **(D)** Mean cholesterol content of 3 dpt zebratish following a 24 h treatment with atorvastatin, water-soluble cholesterol, or both. Ordinary one-way ANOVA with Tukey’s multiple comparisons test with selected adjusted P values shown **(E)** images of control of atorvastatin treated transgenic fish whose macrophages express tdTomato at 2 hpi with ∼*100 M. marinum* expressing a cytosolic blue fluorescent protein reconstituted with DIBO-488 labeled azido-DIM (DIM-488), scale bar = 40pm. Arrows, DIM-488 spread onto epithelial cells. **(F)** Mean percent DIM-488 in macrophage or epithelial cells not localized with bacteria at 2 h following HBV infection with ∼*100 M mannum* in control or atorvastatin treated fish. Two-tailed, unpaired t test. **(G)** Mean bacterial volume following HBV infection of control or atorvastatin treated fish with *∼*100 *M. marinum*. Two-tailed Mann Whitney test for 2 hpi and two-tailed, unpaired t test for 24 hpi **(H)** Percentage of infected or uninfected fish at 3 dpi into the HBV with 1-3 wildtype *M. mannum* with or without atorvastatin and water-soluble cholesterol. **(I)** Percentage of infected or uninfected wildtype or MYD88-depleted fish at 3 dpi into the HBV with 1-3 wildtype or Δ *mmpL7 M marinum* with or without atorvastatin **(H)** and **(I)** Fisher s exact test with Bonferroni’s correction for multiple comparisons. **(B)-(D)** and **(F)-(l)** representative of three separate experiments.

Hypercholesteremia has been shown to exacerbate mycobacterial infection in mice and humans (Martens et al., 2008; Soh et al., 2016), and studies suggest the utility of statins both in TB treatment and prevention. When used in mice, statins decreased the duration of TB therapy by one month (Dutta et al., 2016), and studies of health care databases found that statin use is associated with decreased incidence of TB (Kim et al., 2019; Lai et al., 2016b). Accordingly, we sought to evaluate the role of statins in the reduction of bacterial burdens and in preventing infection. First, we showed that statin treatment reduced bacterial burdens in the zebrafish (Figure 7G). Next, to evaluate if statins prevented infection, we infected zebrafish with 1-3 bacteria, similar to the infectious dose in humans (Bates et al., 1965; Wells et al., 1948), and evaluated their ability to establish infection. This infectivity assay previously found that PDIM-deficient *M. marinum* established infection at a reduced frequency (Cambier et al., 2017). We treated zebrafish with atorvastatin for 24 h prior to infection with 1-3 bacteria and continued daily atorvastatin treatment for the 3-day assay period. At 3 dpi, atorvastatin treatment decreased wildtype *M. marinum*’s infectivity by 50% (Figure 7H). Moreover, restoring cholesterol levels by co-treating with water-soluble cholesterol restored wildtype *M. marinum*’s infectivity (Figure 7H). This result confirmed atorvastatin reduces infectivity by lowering cholesterol rather than to its other effects on physiology and metabolism. However, this protective low-cholesterol state could be due to antimicrobial effects independent of PDIM spread. Altered cholesterol flux can have pleotropic effects on immunity (Tall and Yvan-Charvet, 2015), and mycobacteria have been shown to use cholesterol as a carbon source (Pandey and Sassetti, 2008).

To test the alternative hypothesis that atorvastatin’s protection is not through disrupting the PDIM-TLR axis we evaluated the infectivity of PDIM-deficient Δ*mmpL7 M. marinum*. Consistent with previous reports (Cambier et al., 2017), Δ*mmpL7 M. marinum* exhibited decreased infectivity compared to wildtype bacteria (Figure 7I). In agreement with our model that atorvastatin acts through modulating PDIM spread, we found that atorvastatin treatment did not further decrease Δ*mmpL7*’s infectivity (Figure 7I). However, PDIM-deficient bacteria exhibit such a severe virulence defect, any further attenuation due to other low-cholesterol-dependent antimicrobial mechanisms may not be possible. To address this concern, we took advantage of the fact that Δ*mmpL7*’s virulence defect is reversed in the absence of host TLR signaling (Figure 7I, Cambier et al., 2014b). In this condition where PDIM-deficient bacteria are fully virulent, they should become attenuated by any non-PDIM-associated antimicrobial mechanisms caused by atorvastatin. However, atorvastatin still did not decrease Δ*mmpL7*’s infectivity in MYD88-depleted hosts (Figure 7I). Finally, we showed that atorvastatin decreased the infectivity of wildtype *M. marinum* in a MYD88-dependent fashion (Figure 7I). These data strongly support the idea that atorvastatin acts to disrupt the PDIM-TLR axis. Only in conditions where PDIM is present on bacterial surfaces and when host TLR signaling is intact will lowering cholesterol provide a protective effect. Collectively, these data demonstrate that statins reduce mycobacterial infectivity by reducing host cholesterol, and thereby PDIM’s infiltration of epithelial membranes.

## DISCUSSION

Molecular Koch’s postulates are a guiding set of principles used to characterize bacterial virulence factors (Falkow, 1988; 2004; Ramakrishnan, 2020). Genetic analyses are the mainstay of assigning pathogenic functions to specific virulence factors, including those not directly encoded by the genome. Virulence lipids are often assigned their pathogenic roles from studies of bacterial mutants lacking proteins involved in the lipid’s biosynthesis or transport. Thus, these functions can be directly attributed only to the biosynthetic protein, and not the lipid per se. Here, by combining bioorthogonal chemistry with the zebrafish model of TB, we more rigorously satisfy molecular Koch’s postulates to describe PDIM’s role in virulence. PDIM’s inherent fluidity and the abundance of cholesterol in epithelial membranes influenced PDIM’s ability to infiltrate the host lipid environment. This occupation of epithelial membranes by PDIM mediated subversion of TLR signaling at the site of infection, thus enabling mycobacteria to gain access to non-activated immune cells (Figure 8).

**Figure 8.**
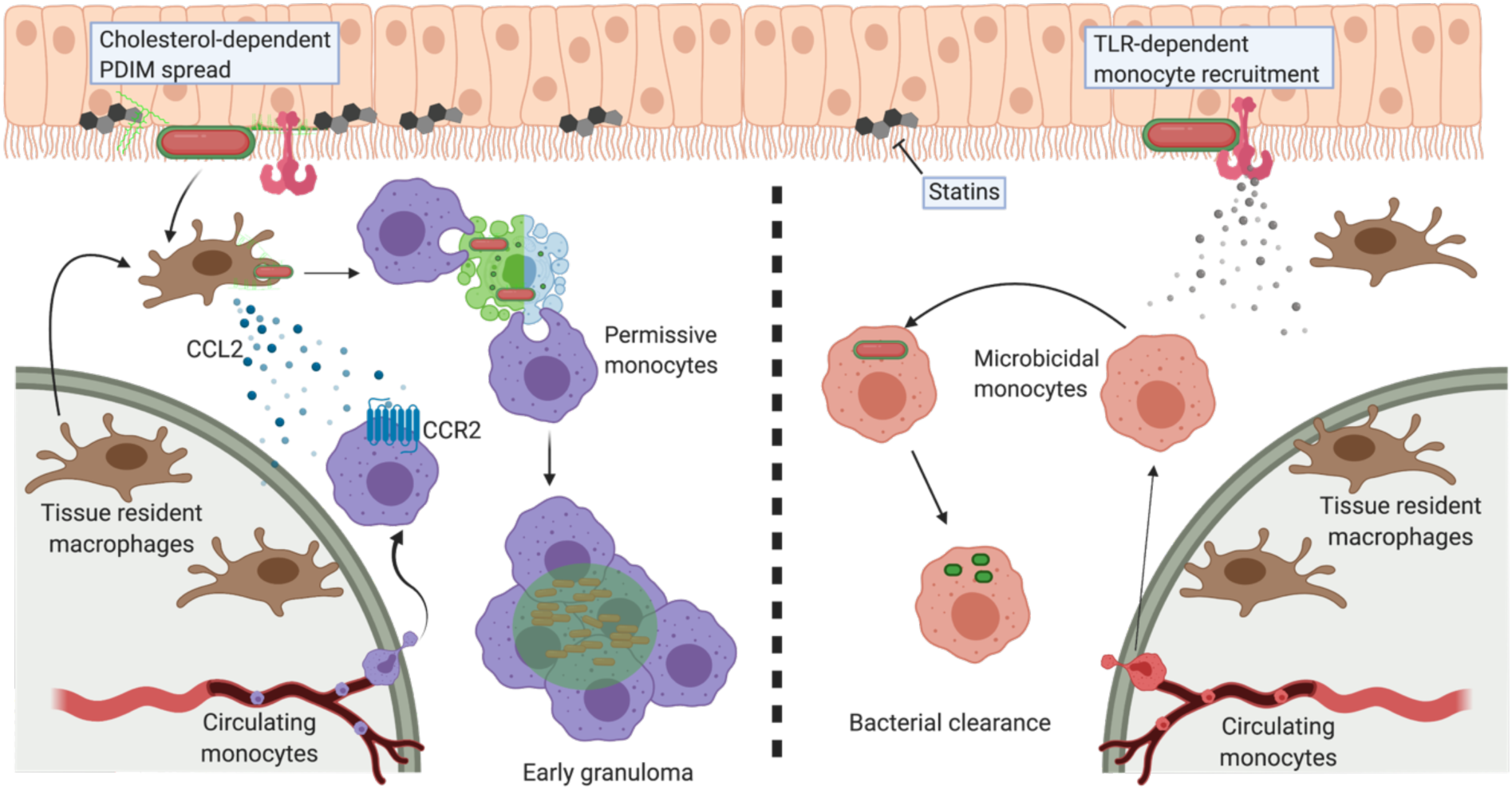
Model of PDIM spreading to promote virulence. Cholesterol-dependent PDIM spread into epithelial membranes prevents TLR detection at the site of infection. Bacteria then drive resident macrophages to produce CCL2 to recruit CCR2-positive permissive monocytes which go on to form early granulomas. PDIM continues to spread into host membranes throughout this process. In low cholesterol settings (statin treatment) PDIM does not spread as readily into epithelial membranes. TLR-dependent recruitment of microbicidal monocytes occurs, which can then clear infecting mycobacteria.

The role epithelial cells play in priming innate immunity is appreciated in several models of bacterial pathogenesis. *Pseudomonas aeruginosa* and *Staphylococcus aureus* are both recognized by TLR-2 on human epithelial cells leading to the production of inflammatory cytokines (Soong et al., 2004). In zebrafish these responses result in bacterial clearance (Cambier et al., 2014b), highlighting this pathway as a conserved response to infection. In contrast, human epithelial cells infected by mycobacteria have no significant change in global transcription, suggesting minimal immune detection (Reuschl et al., 2017). Our data now demonstrate that PDIM plays a role in silencing epithelial cell responses, promoting uptake by non-activated macrophages.

Previous models suggest PDIM facilitates a passive evasion of TLRs by masking underlying TLR ligands (Cambier et al., 2014b; 2017). This model was based on findings that PDIM-expressing mycobacteria were rendered growth-attenuated if co-infected with TLR-stimulating bacteria (Cambier et al., 2014b). While our current model suggests a more active role for PDIM, we still find that TLR-stimulating bacteria cause attenuation of wildtype bacteria. Specifically, bacteria unable to spread PDIM caused attenuation of bacteria that can spread PDIM (Figure 6B and C). These findings argue that PDIM’s influence over innate immune signaling is spatially restricted. PDIM appears to only inhibit TLR signaling at membrane sites where it has spread, emphasizing the need for PDIM localization on the bacterial surface prior to spreading into epithelial membranes. Once spread, the potential for PDIM to disrupt membrane protein signaling is intriguing. Several biophysical properties of membranes are known to mediate membrane protein function (Los and Murata, 2004), and PDIM has been implicated in altering host membrane structure (Augenstreich et al., 2019). Furthermore, TLRs have been shown to aggregate into cholesterol rich lipid domains to facilitate downstream signaling events (Ruysschaert and Lonez, 2015). How PDIM’s affinity for cholesterol promotes spread and whether this association disrupts TLR-cholesterol lipid domains is yet to be determined.

The finding that PDIM’s methyl-branched mycocerosic acids are required for virulence suggests pathogenic mycobacteria have evolved a lipid-based immunomodulatory strategy similar to that seen at host barrier tissues. While predominant across prokaryotes, methyl-branched fatty acids are only found in certain secretions of higher eukaryotes. In particular, vernix caseosa, a white wax that covers neonates during the third trimester of gestation, is implicated in promoting normal gut development and microbial colonization (Nishijima et al., 2019). The fetus swallows a significant amount of vernix shed in the amniotic fluid, in increasing amounts as term birth approaches. Premature infants are exposed to substantially less vernix and have increased rates of necrotizing enterocolitis (Ran-Ressler et al., 2011). In a rat model of necrotizing enterocolitis, feeding premature pups methyl-branched lipids led to a decrease in disease development and a shift from a pro-inflammatory to an anti-inflammatory gut environment (Ran-Ressler et al., 2011). These studies implicate lipid-based immunomodulation as playing a key role in tissue-barrier homeostasis, dampening immune responses to allow for proper microbial colonization. Mycobacteria appear to have adapted this strategy of using methyl-branched lipids as an immunomodulatory mechanism to establish infection and thereby ensure their evolutionary survival.

Systemic cholesterol metabolism in humans influences the lung environment, including the abundance of cholesterol in lung epithelial membranes (Fessler, 2017). Therefore, cholesterol-promoted PDIM spreading may also influence *M. tuberculosis* transmission. While a clinical trial is currently underway evaluating the use of statins alongside standard TB therapy (Karakousis et al., 2019), our data instead argue for the use of statins as a TB preventative therapy.

## ACKNOWLEDGEMENTS

We thank Lalita Ramakrishnan for help interpreting data and manuscript writing, Karen Dobos for help purifying PDIM, Benjamin M. Swarts for providing TreT-expressing *E. coli*, and David Tobin for zebrafish lines. This work was supported by a National Institutes of Health grant (AI51622) to C.R.B. C.J.C. was supported by a Damon Runyon Postdoctoral Fellowship. S.M.B and J.A.B were supported by NIGMS F32 Postdoctoral Fellowships.

## AUTHOR CONTRIBUTIONS

C.J.C., and C.R.B. conceived the project. C.J.C., S.M.B. and J.A.B. carried out experiments and interpreted data. C.J.C. and C.R.B. wrote the manuscript with input from all authors. C.R.B. provided supervision.

## MATERIALS AND METHODS

### RESOURCES TABLE

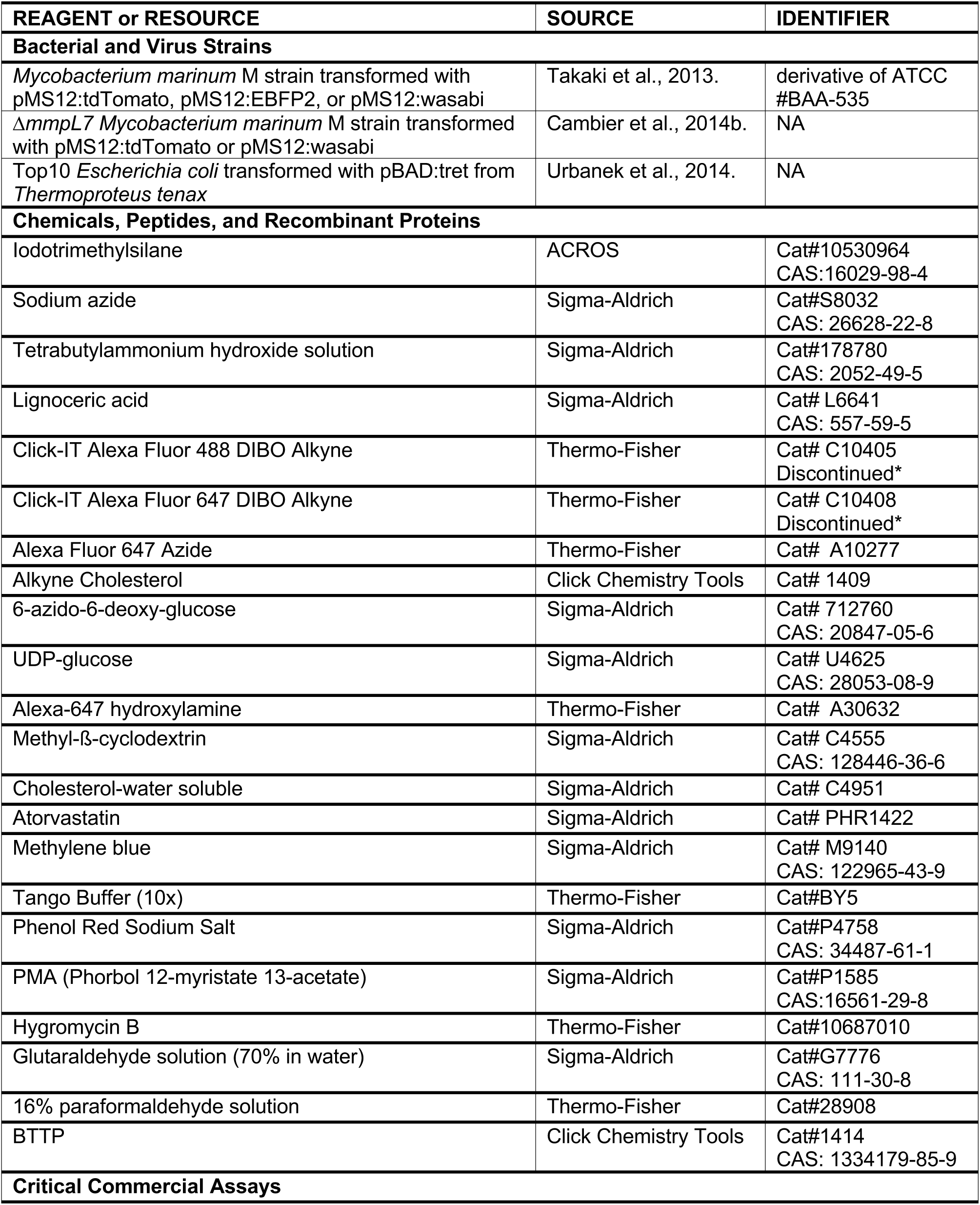
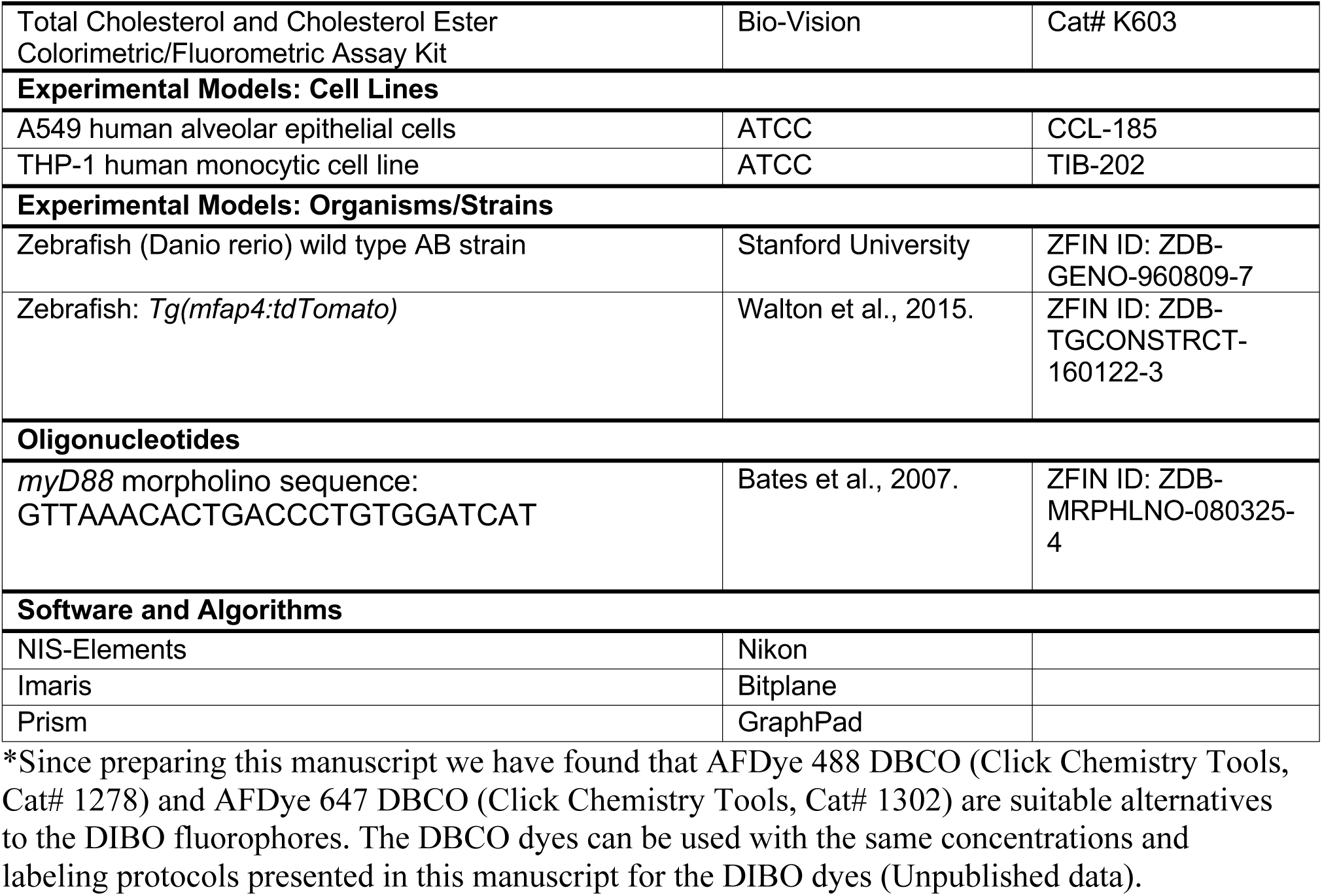

#### Procedures and materials for synthetic chemistry

All reactions were performed in dry standard glassware fitted with rubber septa under an inert atmosphere of nitrogen unless otherwise stated. Preparative thin-layer chromatography (TLC) was performed with Millipore’s 1mm and 0.2mm silica gel 60 pre-coated glass plates. Analytical TLC was used for reaction monitoring and product detection using pre-coated glass plates covered with 0.20 mm silica gel with fluorescent indicator; visualized by UV light and 10% CuSO4 in 1.3M phosphoric acid in water. Reagents were purchased in reagent grade from commercial suppliers and used as received, unless otherwise described. Anhydrous dichloromethane (DCM) was prepared by passing the solvent through an activated alumina column.

#### Chemical Analysis Instrumentation

Proton (^1^H NMR) and proton-decoupled carbon-13 (^13^C {^1^H} NMR) nuclear magnetic resonance spectra were recorded on an Inova-500 spectrometer at 25 °C, are reported in parts per million downfield from tetramethylsilane, and are referenced to the residual protium (CDCl_3_: 7.26 [CHCl_3_]) and carbon (CDCl_3_: 77.16) resonances of the NMR solvent. Data are represented as follows: chemical shift, multiplicity (br = broad, s = singlet, d = doublet, t = triplet, q = quartet, quin = quintet, sept = septet, m = multiplet), coupling constants in Hertz (Hz), integration. Mass spectra were obtained on a Bruker Microflex MALDI-TOF by mixing 0.5µl of 1mg/ml sample in chloroform with 0.5µl of 10mg/ml 2,5-dihydroxybenzoic acid before spotting onto a 96-well MALDI plate.

#### Isolation of PDIM

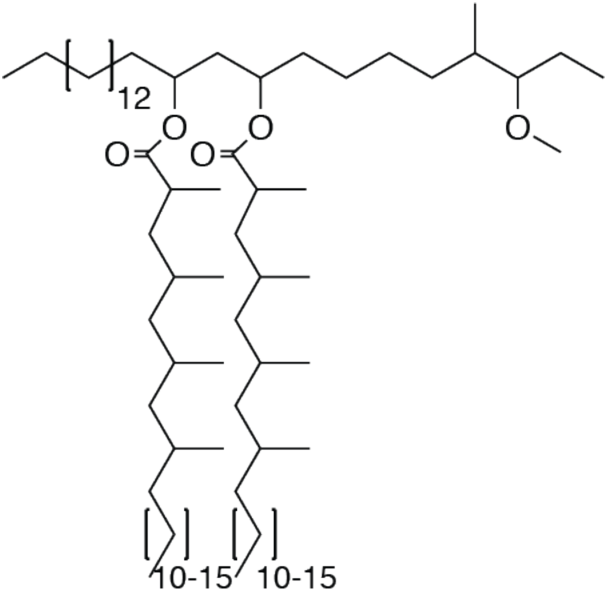

Wildtype *M. marinum* was grown in GAS medium without Tween-80 to an OD_600_ of 1.2. Bacteria were pelleted, frozen and lyophilized on a Labconoco FreezeZone 4.5 plus. Bacterial lipids were then extracted by stirring lyophilized pellet in petroleum ether for 1 h at 23 °C. Bacteria were allowed to settle, and solvent was collected. Remaining bacteria were re-extracted 3-5 times. The solvent extract was then passed through a 0.2µm PTFE filter, and crude lipids were concentrated under reduced pressure. Crude lipid extracts were separated by preparative TLC in the solvent system 98:2 petroleum ether:ethyl acetate. The band correlating to PDIM at an Rf = 0.4 was isolated. Preparative TLC was repeated twice more to further purify PDIM. Average PDIM isolated from *M. marinum* was 60mg per 10 liters of culture. _1_H NMR (500 MHz, CDCl_3_) δ 4.93-4.87 (m, 2H), 3.32 (s, 3H), 2.87-2.83 (m, 1H), 2.57-2.51 (m, 2H), 1.90-0.81 (m, 166H). ^13^C NMR (126 MHz, CDCl_3_) δ 176.69, 176.63, 86.78, 86.73, 77.41, 77.37, 77.16, 76.90, 70.72, 70.67, 57.51, 45.56, 45.51, 41.41, 41.33, 38.56, 37.85, 37.73, 37.09, 36.77, 34.91, 34.18, 34.07, 32.83, 32.08, 30.24, 29.96, 29.87, 29.82, 29.75, 29.62, 29.53, 28.34, 27.64, 27.36, 27.25, 27.16, 27.12, 25.74, 25.35, 22.85, 22.74, 22.43, 20.81, 20.57, 20.52, 20.37, 20.27, 18.67, 18.58, 18.53, 14.80, 14.75, 14.28, 14.22, 10.26, 1.16. MALDI-TOF for: C82H162O5 Calc’d [M+Na^+^]=1250.23; found 1250.41. C84H166O5 Calc’d [M+Na^+^]=1278.26, found 1278.40. C86H170O5 Calc’d [M+Na^+^]=1306.29, found 1306.55.

#### Synthesis of azido-DIM

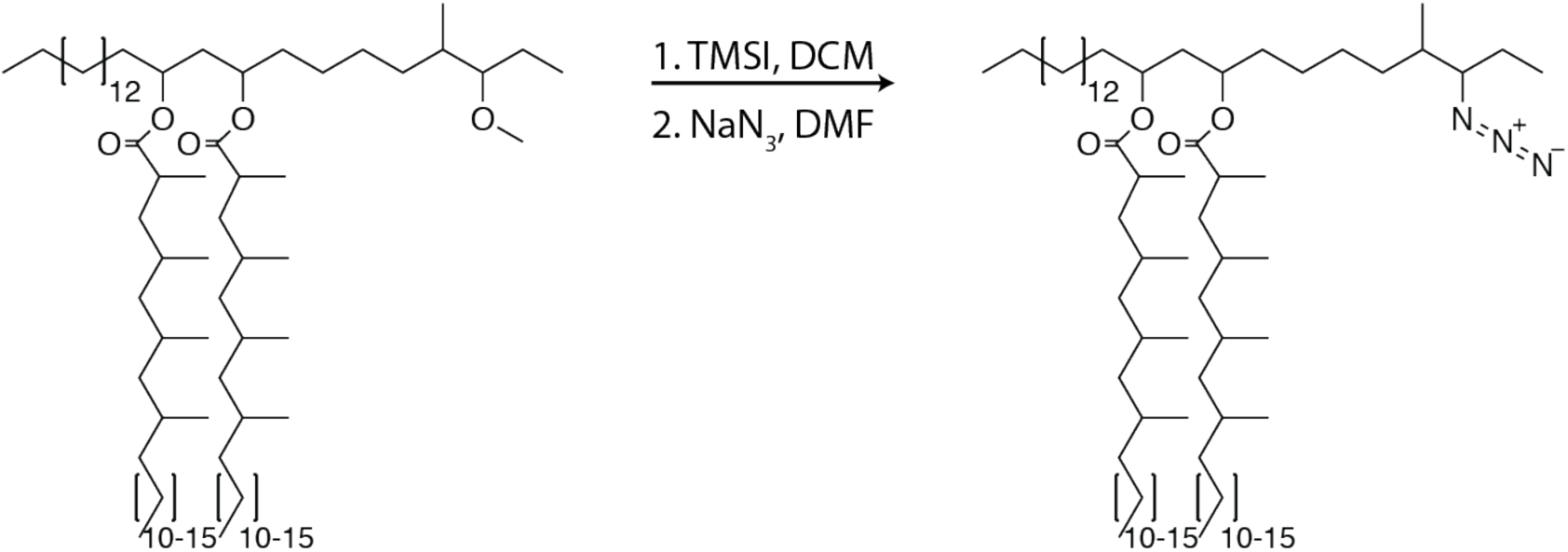

To PDIM (75.6mg, 60.2µmol, 1.0 equiv.) was added 0.3ml of DCM, the mixture was stirred and iodotrimethylsilane (TMSI, 258µl, 1.8mmol, 30.0 equiv.) was added and allowed to react for 12 h at 23 °C. The reaction was concentrated under reduced pressure. NaN3 (39.1mg, 602µmol, 10.0 equiv.) was added followed by 0.3ml of anhydrous dimethylformamide (DMF) and stirred for 12 h at 23 °C. The solvent was removed under reduced pressure and the product was purified using preparative TLC in 98:2 petroleum ether:ethyl acetate Rf=0.5 as a white wax (37.7mg, 29.8µmol, 50%). ^1^H NMR (500 MHz, CDCl_3_) δ 5.06-4.86 (m, 2H), 3.30-3.21 (m, 1H), 2.58-2.50 (m, 2H) 2.1-0.81 (m, 163H). ^13^C NMR (126 MHz, CDCl_3_) δ 176.73, 176.63, 77.41, 77.36, 77.16, 76.91, 71.08, 70.74, 59.86, 45.56, 41.96, 41.45, 41.35, 39.27, 37.84, 37.73, 37.24, 37.08, 37.01, 35.28, 34.82, 34.18, 32.08, 30.32, 30.24, 30.22, 30.04, 29.96, 29.87, 29.82, 29.75, 29.72, 29.67, 29.62, 29.52, 29.39, 28.40, 28.34, 27.72, 27.36, 27.25, 27.18, 27.16, 26.20, 25.35, 25.27, 25.23, 22.85, 21.28, 21.13, 20.35, 20.32, 20.27, 18.73, 18.66, 18.58, 18.53, 14.56, 14.29, 13.87. MALDI-TOF for: C81H159N3O4 Calc’d [M+Na^+^]=1261.22; found 1260.92. C83H163N3O4 Calc’d [M+Na^+^]=1289.25, found 1289.02. C85H167N3O4 Calc’d [M+NH4^+^]=1312.33, found 1312.37.

#### Hydrolysis of PDIM and isolation of phthiocerol

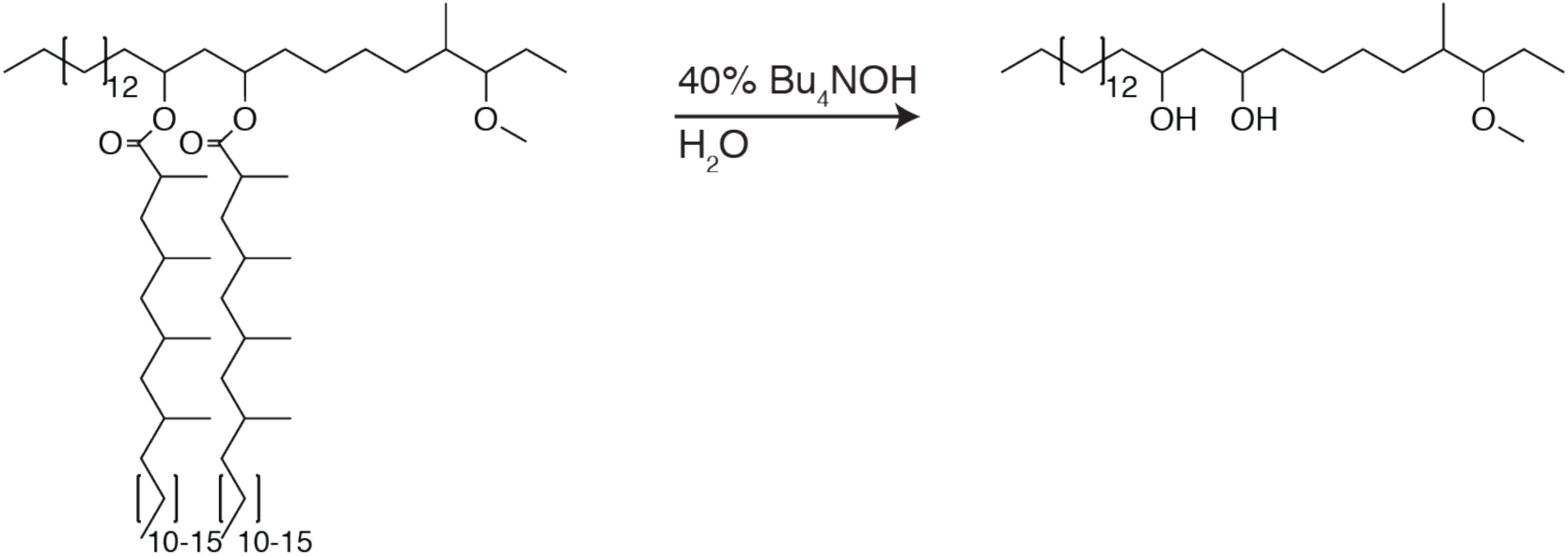

To dried PDIM (66.7mg, 53.1µmol) was added 0.5ml of 40% tetrabutylammonium hydroxide in water, the mixture was sealed, aggressively stirred, and allowed to react for 48 h at 120 °C. Reaction was cooled to 23 °C and 4M HCL was added dropwise until pH was < 3. The reaction was extracted with DCM and washed once with water. The solvent was removed under reduced pressure and phthiocerol was purified using preparative TLC in 80:20 petroleum ether:ethyl acetate Rf = 0.4. (18.8mg, 42.5µmol, 80%).^1^H NMR (500 MHz, CDCl_3_) and ^13^C NMR (126 MHz, CDCl_3_) matched previously reported spectra. ESI HRMS for: C28H58O3 Calc’d [M+H^+^]=443.4464; found 443.4449. C29H60O3 Calc’d [M+H^+^]=457.4621; found 457.4616. C30H62O3 Calc’d [M+H^+^]= 471.4777, found 471.4762.

#### Synthesis of PDIF

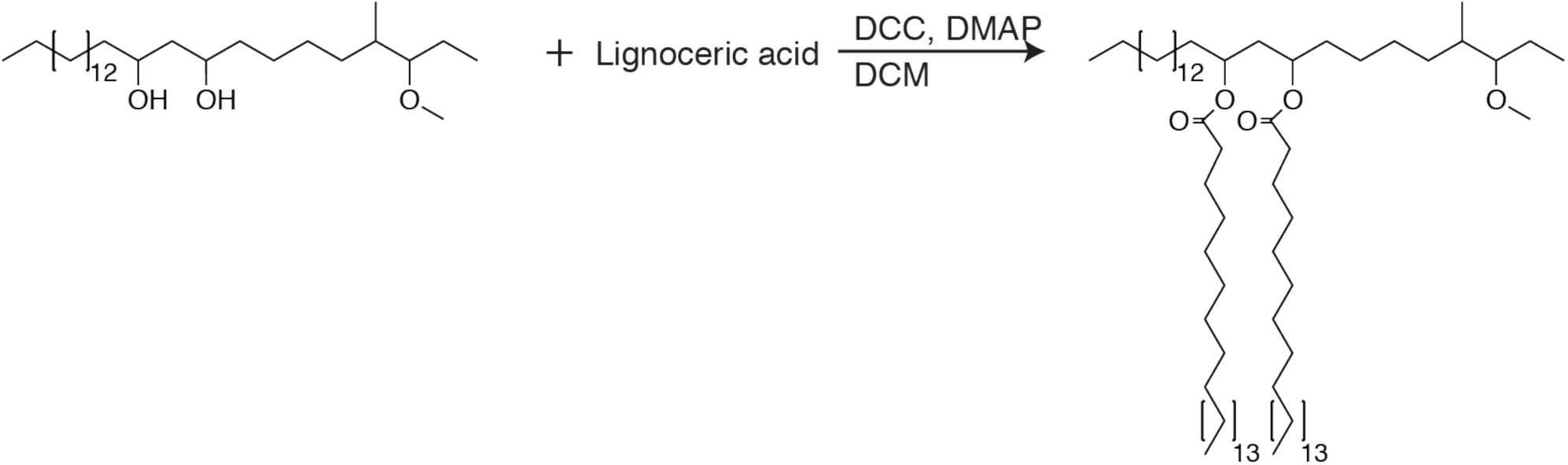

To phthiocerol (18.8mg, 42.5µmol, 1 equiv.) was added lignoceric acid (47mg, 127.5µmol, 3.0 equiv.), N,N’-dicyclohexylcarbodiimide (DCC, 35mg, 170µmol, 4.0 equiv.), 4-Dimethylaminopyridine (DMAP, 20.77mg, 170µmol, 4.0 equiv.), and 2.3ml of DCM. The reaction was stirred and allowed to react for 16 h at 23 °C. The solvent was removed under reduced pressure and the product was purified using preparative TLC in 98:2 petroleum ether:ethyl acetate Rf=0.4. (12.8mg, 11.5µmol, 27%). ^1^H NMR (500 MHz, CDCl_3_) δ 4.94-4.87 (quin, 2H), 3.33 (s, 3H), 2.88-2.83 (m, 1H), 2.30-2.25 (t, 4H), 1.75-0.79 (m, 141H). ^13^C NMR (126 MHz, CDCl_3_) δ 173.42, 86.66, 77.27, 77.02, 76.76, 70.97, 57.40, 38.45, 34.84, 34.67, 34.13, 34.09, 32.62, 31.94, 29.73, 29.68, 29.55, 29.47, 29.38, 29.34, 29.23, 27.44, 25.57, 25.17, 25.10, 22.70, 22.37, 14.76, 14.12, 10.10. MALDI-TOF for: C76H150O5 Calc’d [M+Na^+^]=1166.14, found 1166.14. C77H152O5 Calc’d [M+Na^+^]=1180.15, found 1180.24. C78H154O5 Calc’d [M+Na^+^]=1194.17; found 1194.25.

#### Synthesis of azido-DIF

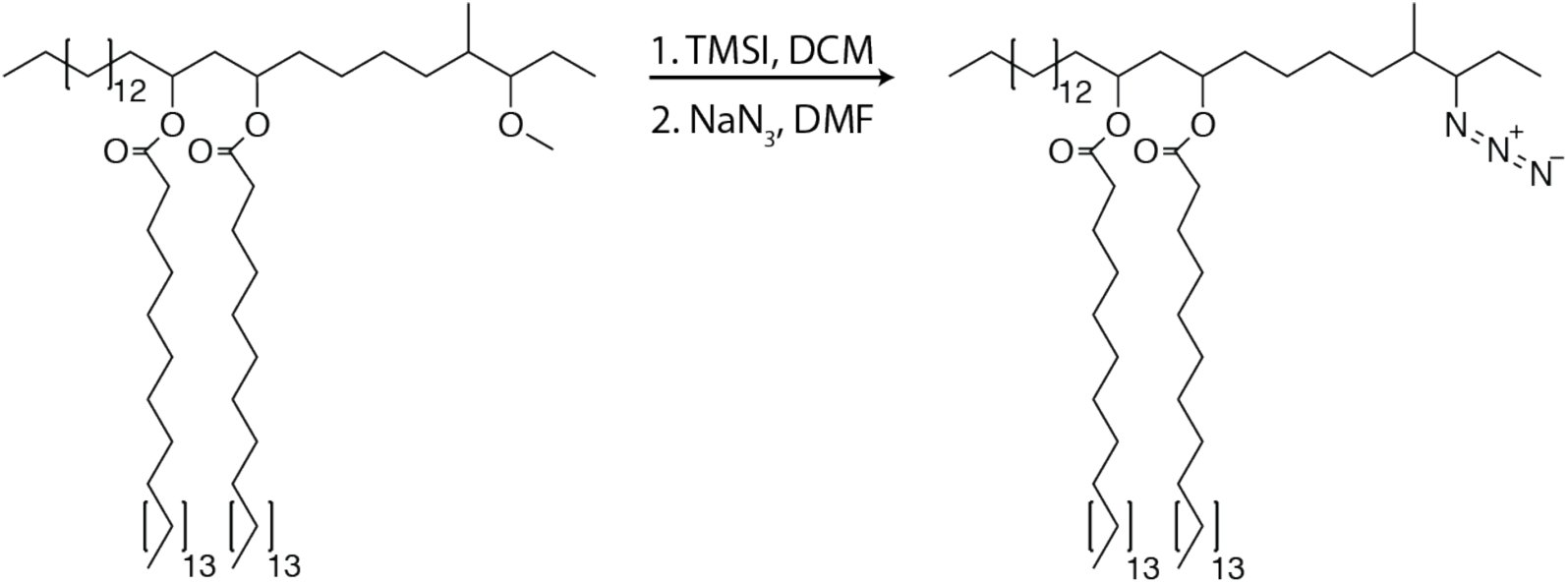

To PDIF (12.8mg, 11.5µmol, 1 equiv.) was added 0.3ml of DCM, the mixture was stirred and TMSI (49µl, 345µmol, 30.0 equiv.) was added and allowed to react for 3 h at 23 °C. The reaction was concentrated under reduced pressure. NaN_3_ (7.5mg, 115µmol, 10.0 equiv.) was added followed by 0.3ml of anhydrous dimethylformamide (DMF) and stirred for 12 h at 23 °C. The solvent was removed under reduced pressure and the product was purified using preparative TLC in 98:2 petroleum ether:ethyl acetate Rf=0.5. (6.9mg, 6µmol, 52%). ^1^H NMR (500 MHz, CDCl_3_) δ 5.08-5.00 (m, 2H), 3.32-3.20 (m, 2H), 3.12-3.04 (m, 1H), 2.33-2.25 (t, 4H), 1.80-0.73 (m, 138H). ^13^C NMR (126 MHz, CDCl_3_) δ 173.41, 77.27, 77.02, 76.76, 71.16, 70.05, 59.72, 39.09, 36.92, 35.12, 34.70, 34.64, 32.34, 31.94, 29.72, 29.67, 29.58, 29.53, 29.42, 29.38, 29.33, 29.23, 26.05, 25.15, 24.02, 22.70, 21.15, 16.10, 14.13, 11.06, 1.03. ESI C75H148NO4 calc’d [M+K, - N_2_]=1166.10, found 1166.05.

#### Zebrafish Husbandry and Infections

Wild-type AB (Zebrafish International Resource Center), and *Tg(mfap4:tdTomato)* (Walton et al., 2015) lines were maintained in buffered reverse osmotic water systems. Fish were fed twice daily a combination of dry feed and brine shrimp and were exposed to a 14 h light, 10 h dark cycle to maintain proper circadian conditions. Zebrafish embryos were maintained at 28.5 °C in embryo media which consisted of the following dissolved in Milli-Q water (%weight/volume): 0. 0875% sodium chloride, 0.00375% potassium chloride, 0.011% calcium chloride, 0.00205% monopotassium phosphate, 0.00089% disodium phosphate, and 0.0493% magnesium sulfate. Embryo media was then buffered to pH 7.2 with sodium bicarbonate. Embryos were maintained in 0.25 mg/ml methylene blue (Sigma) from collection to 1-day post-fertilization (dpf). 0.003% PTU (1-phenyl-2-thiourea, Sigma) was added from 24 h post-fertilization (hpf) on to prevent pigmentation. Larvae (of undetermined sex given the early developmental stages used) were infected at 48 hpf via the hindbrain ventricle (HBV) using single-cell mycobacterial suspensions of known titer. Number of animals to be used for each experiment was guided by pilot experiments or by past results with other bacterial mutants and/or zebrafish. On average 15 to 40 larvae per experimental condition were required to reach statistical significance and each experiment was repeated at least three times. Larvae were randomly allotted to the different experimental conditions. The zebrafish husbandry briefly described above and all experiments performed on them were in compliance with the U.S. National Institutes of Health guidelines and approved by the Stanford Institutional Animal Care and Use Committee.

#### Bacterial Strains and Methods

*M. marinum* strain M (ATCC BAA-535) and Δ*mmpL7* mutants (Cambier et al., 2014b) expressing either TdTomato, Wasabi, or EBFP2 under the control of the *msp12* promoter were grown under hygromycin (Thermo-Fisher) selection in 7H9 Middlebrook’s medium (Difco) supplemented with 10% OADC (Fisher), 0.2% glycerol, and 0.05% Tween-80 (Sigma). Where noted bacteria were also grown in glycerol-alanine-salts (GAS) medium, recipe (%weight/volume) in 18mM sodium hydroxide in Milli-Q water pH 6.6 +/- 0.05% Tween-80: 0.03% BactoCasitone (BD Science), 0.005% ferric ammonium citrate (Sigma), 0.4% potassium phosphate dibasic anhydrous (VWR), 0.2% citric acid, anhydrous (VWR), 0.1% L-alanine (Sigma), 0.12% magnesium chloride, heptahydrate (VWR), 0.06% potassium sulfate (VWR), 0.2% ammonium chloride (VWR), and 1% glycerol. To prepare heat-killed *M. marinum*, bacteria were incubated at 80°C for 20 min. To prepare fixed bacteria, bacteria were incubated in described concentrations of glutaraldehyde (Sigma) and/or paraformaldehyde (Thermo-Fisher) for 1hr at 23°C, followed by 3 washes with PBS prior to experimental use.

#### Cell Lines and infections

Cells were grown in T75 flasks (Thermo-Fisher) and maintained at 37 °C and 5% CO2. THP-1 were grown in RPMI supplemented with 10% fetal bovine serum (FBS). A549 were grown in DMEM supplemented with 10% FBS. Two days prior to infection, THP-1s were plated on 8-well Nunc Lab-Tek II chamber slides (Thermo-Fisher) at a density of 150,000 cells per well with 100nM PMA (phorbol 12-myristate 13-acetate, Sigma) in growth media and incubated at 37°C and 5% CO2 for 24 h. One day prior to infection, the media on THP-1s was replaced with fresh growth media and the cells were incubated at 33°C and 5% CO2 for 24 h, cells were then infected with *M. marinum* at a multiplicity of infection (MOI) of 2 in growth media. Infection was allowed to progress for 6 h prior to washing twice with PBS and replacing with growth media. THP-1s were then incubated for 24 h at 33°C and 5% CO2 prior to experimental end point. One day prior to infection, A549s were plated on 8-well chamber Nunc Lab-Tek II chamber slides (Thermo-Fisher) at a density of 50,000 cells per well in growth media and incubated at 37°C and 5% CO2. Day of infection cells were moved to 33°C and 5% CO2 for 3 h, and then were infected with *M. marinum* at a MOI of 5 and incubated at 33°C and 5% CO2 for 24 h. Infected THP-1s and A549s were then imaged as described below.

#### Extraction and reconstitution of *M. marinum*

##### Extraction

1 liter of *M. marinum* were grown in GAS medium plus Tween-80 to an OD_600_ of 1.2. Bacteria were pelleted in a glass 50mL conical tubes (Fisher) of known weight, frozen and lyophilized. The dry bacteria in 50mL conical tubes were then weighed and the dry bacterial weight was calculated. 25mL of petroleum ether were then added to the bacteria, and the conical tube was capped with a PTFE lined lid (Sigma) and the suspension was vortexed for 3min. An additional 25mL of petroleum ether was added and the sample was centrifuged for 3min at 1000xg at 4°C. The extract was then saved or discarded depending on the downstream experimental applications and the bacteria were extracted once more as above. For total lipid extractions (not used for reconstitution), bacteria were treated with 1:1 chloroform:methanol for 12 h at 60°C. Extracts were filtered, back extracted with water to remove water soluble contaminants, dried under reduced pressure and used for downstream experiments.

##### Reconstitution

Prior to extractions the following calculation was used to determine the amount of lipids to add back to each pellet.

(23mg lipid/gram dry bacteria)*(weight in grams of dry bacteria)/(0.75 reconstitution efficiency) Where 23mg of lipid per gram of dry bacteria is the experimentally determined average amount of lipid removed during petroleum ether extraction of bacteria grown in these conditions (Figure S1B, initial extraction) and the 0.75 reconstitution efficiency was also experimentally determined (Figure S1C). If reconstituting with DIM variants, 30% of the above weight consists of DIMs and the remainder will consist of DIM-depleted petroleum ether lipid extracts. Following two rounds of petroleum ether extraction as detailed above, bacteria were immediately mixed with pre-determined lipid mixtures (or no lipid at all for delipidated bacteria) in petroleum ether (2-3 mls) followed by extended drying under reduced pressure. Dried bacteria were then rescued into 7H9 media supplemented with 10% OADC, and 0.2% glycerol (prep media) and subjected to single cell preparation protocol.

#### *M. marinum* single cell preparation

For more thorough details, as well as rationale and explanation of the following protocol see Takaki et al., 2013. Bacteria were washed once with 15mL of prep media followed by resuspension in 500µl of prep media. Bacteria were then passed through a 27-gauge needle 10 times, followed by the addition of 1ml of prep media and centrifugation at 100xg for 3min. 1ml of supernatants were saved. This process was repeated 3-5 times. Collected supernatants were then passed through a 5.0µm acrodisc versapor membrane syringe filter (VWR). The filtrate was then pelleted at 16,000xg for 2min, pellets were resuspended in prep media to a concentration of around 1×10^8-9 bacteria per ml, aliquoted and stored at −20C for future use, or immediately subjected to copper-free click chemistry reactions.

#### Metabolic labeling of *M. marinum* with 6-TreAz

##### Expression and Purification of TreT

TreT was expressed and purified utilizing a similar method as previously reported (Urbanek et al., 2014). Top10 *E. coli* expressing the *tret* gene from *Thermoproteus tenax* (pBAD plasmid, AraC control) were streaked onto a Lysogeny broth (LB) agar plate supplemented with 100 µg/mL ampicillin and incubated at 37 °C for 24 h. A single colony was picked and used to inoculate 5 mL of LB liquid medium containing 100 µg/mL ampicillin. The starter culture was placed in a shaking incubator (175 rpm) at 37 °C for 16 h. The starter culture was then transferred to a 1 L solution of sterilized Terrific broth (TB) supplemented with 100 µg/mL ampicillin in a sterilized 2 L Fernbach culture flask. The flask was shaken (175 rpm) at 37 °C. When the absorbance at 600 nm reached between 0.6 and 0.9 (typically 4-5 h post inoculation), TreT expression was induced by adding 1 mL of 1 M arabinose solution in sterile water (1 mM final concentration). The flask was again shaken (175 rpm) at 37 °C for another 20 h. The culture was then transferred to a polypropylene bottle and pelleted for 15 minutes at 4,000 x g at 4 °C. The supernatant was discarded, and the pellet was suspended in 45 mL of lysis buffer (50 mM NaH_2_PO_4_, 500 mM NaCl, 20 mM imidazole, pH 7.4) and run through a homogenizer 5 times utilizing ice to keep the solution cool. Once homogenized, the lysate was clarified by centrifugation pelleting for 20 minutes at 21,000 x g at 4 °C and filtering through a 0.45 µm Teflon syringe filter. To the clarified lysate (generally about 50 mg/mL protein content as measured with absorbance at 280 nm) was then added 5 mL of pre-washed (lysis buffer) Ni-NTA resin slurry (Qiagen). The suspension was mixed on an orbital shaker for 60 – 90 minutes at 4 °C and then transferred to a glass column (BIO-RAD EconoColumn). Non-His-Tagged proteins were eluted with lysis buffer until the absorbance at 280 nm matched background levels utilizing 50 – 100 mL of lysis buffer. His-tagged TreT was eluted with elution buffer (50 mM NaH_2_PO_4_, 500 mM NaCl, 250 mM imidazole, pH 7.4) in multiple 2.5 mL increments, until protein elution was determined complete by absorbance at 280 nm. Buffer exchange to a storage/reaction buffer (50 mM Tris, 300 mM NaCl, pH 8.0) was performed with a desalting column (PD-10, GE Healthcare) and the protein was transferred to a conical tube and diluted to 1 mg/mL as determined by absorbance at 280 nm for long-term storage. Storage of the enzyme at 4 °C and at this concentration yields active protein that does not have significant losses in activity even after 12 months of storage under these conditions.

##### Synthesis and Purification of 6-Azido-6-deoxy-trehalose (6-TreAz)

To a 1.5 mL Eppendorf tube was added 200 µL of a 100 mM solution of 6-azido-6-deoxy-glucose (final concertation 20 mM), 100 µL of a freshly-prepared 400 mM solution of UDP-glucose (final concentration 40 mM), 100 µL of a 200 mM solution of MgCl_2_ (final concentration 20 mM), and 300 mL of reaction buffer (50 mM Tris, 300 mM NaCl, pH 7.4). Then, 300 µL of TreT solution in storage/reaction buffer was added (final concentration 300 µg/mL) and the reaction vessel was closed and mixed gently by inverting the tube. The reaction was heated to 70 °C for 60 – 90 minutes with shaking at 400 rpm and then cooled on ice before further manipulation. The reaction contents were transferred to a pre-washed Amicon centrifugal filter with a nominal molecular weight limit (NWML) of 10 kDa. The filter was washed with DI water (2 x 1 mL) to facilitate maximal recovery. The upper chamber was discarded and 1 g of prewashed mixed-bed ion-exchange resin (DOWEX) was added to the filtrate and the slurry was equilibrated for 60 – 90 minutes. The suspension was filtered, and the resin was rinsed with 3 – 5 mL of DI water. Analysis by TLC (5:3:2 *n*-butanol: ethanol: DI water) and/or LC-MS equipped with a Supelco aminopropyl column [4.6 x 250 mm, 5 µm] (isocratic 80% ACN in DI water, 0.5 mL/min flowrate) indicated full and complete conversion of 6-azido-6-deoxy-glucose to 6-azido-6-deoxy-trehalose with high purity as determined by nuclear magnetic resonance (NMR), which matched previously reported spectra.

##### Synthesis and Direct utilization of 6-TreAz for labeling Mycobacterium marinum

To facilitate easier production and labeling of *M. marinum* with 6-TreAz, the reaction was performed as above but 6-TreAz was not isolated prior to labeling. Upon reaction completion, the reaction was cooled on ice to bring the vessel back to ambient temperature. The 10 mM stock of 6-TreAz was then added to cultures of *Mycobacterium marinum* to a final concentration of 50 µM.

#### Periodate-hydroxylamine staining of mycobacterial surfaces

Surface-exposed terminal oxidizable carbohydrates were labeled with hydroxylamine following periodate oxidation. Lyophilized control or reconstituted *M. marinum* were washed twice with PBS and resuspended in 0.1 M sodium acetate (Sigma), pH 5.5 containing 1 mM sodium periodate (Sigma). Following a 20-min incubation at 4°C with gentle rotation, 0.1 mM glycerol was added to stop the reaction. Cultures were washed three times with PBS and then subjected to single cell preparation. Following single cell preparation, pellets were transferred to a 96-well plate and then incubated with PBS containing 1 mM Alexa-647 hydroxylamine (Thermo-Fisher). Following a 2 h incubation at 23 °C, the cultures were washed 5 times with PBS and twice with prep media.

#### Copper-free click chemistry of *M. marinum*

Following reconstitution or metabolic labeling, bacteria were treated for single cell preparation. Bacteria were then transferred to 96-well v-bottom dishes and were washed twice with PBS using centrifugation at 3000xg for 3min between washing. Bacteria were then stained with either 5µM DIBO-488 (Thermo-Fisher), or 30µM DIBO-647 (Thermo-Fisher) in 200µl PBS for 90min at 23 °C protected from light. Bacteria were then washed 5 times in PBS followed by 2 washes in prep media. Bacteria were then aliquoted and stored at −20C for future use. Staining efficiency was evaluated by flow cytometry on a BD-Accuri C6 Plus and analysis was performed using the FlowJo software package. Staining was also evaluated by microscopy with a 60x oil-immersion Plan Apo 1.4 NA objective on the Nikon A1R confocal microscope.

#### MYD88 morpholino and liposome injections

To generate MYD88 knockdown zebrafish larvae, the MYD88 morpholino 5’GTTAAACACTGACCCTGTGGATCAT3’ (Bates et al., 2007) was diluted to 2mM in 0.5 x tango buffer (Thermo Scientific), containing 2% phenol red sodium salt solution (Sigma). 1 nl of the morpholino mixture was injected into the 1-4 cell stage of the developing embryo. Lipo-PBS and lipo-clodronate (http://clodronateliposomes.org) were diluted 1:10 in PBS and injected into 2-dpf-old larvae in ∼10 nl via the caudal vein.

#### Confocal microscopy and image-based quantification of infection

Larvae were embedded in 1.5% low melting point agarose (Thermo-Fisher) and a series of z stack images with a 2µm step size was generated through the infected HBV. For infected THP-1 and A549 cells, a series of z stack images with a 1µm step size was generated. Images were captured using the galvo scanner (laser scanner) of the Nikon A1R confocal microscope with a 20x Plan Apo 0.75 NA objective. Higher resolution images were generated using a 40x water-immersion Apo 1.15 NA objective. Bacterial burdens were determined by using the 3D surface-rendering feature of Imaris (Bitplane Scientific Software). Spread lipid images were generated by subtracting the bacterial surface from the lipid channel. 3D surface-rendering was then done in Imaris on both the total lipid and spread lipid images to generate a percent spread value.

#### Macrophage and Monocyte hindbrain recruitment assay

For the macrophage recruitment (Figure 1G): 2 dpf zebrafish were infected in the HBV with *M. marinum* at the dose reported in the figure legends. At 3 h post infection, the number of total myeloid cells in the HBV was quantified using differential interference contrast microscopy using a 20x Plan Fluor 0.75 NA objective on Nikon’s Ti eclipse inverted microscope. For quantification of monocytes (Figure 5A and B): 2 dpf zebrafish were injected in the caudal vein with 200µg/ml of the nuclear stain Hoechst 33342 (Thermo-Fisher) 2 hours prior to HBV infection. Hoechst is unable to cross the blood-brain barrier and therefore will label circulating monocytes but will not label brain resident macrophages (Cambier et al., 2017). At 3hpi blue-fluorescent cells in the hindbrain was quantified similar to above macrophage recruitment assay.

#### Co-infection Experiments

Around 40-50 tdTomato expressing wildtype *M. mairnum* were co-infected with an equal number of wasabi expressing *M. marinum* into the HBV. At 3 dpi the bacterial volume of the wildtype tdTomato expressing *M. marinum* was quantified as described above (*Confocal microscopy and image-based quantification of infection*).

#### Infectivity Assay

3 days post fertilization larvae were infected via the hindbrain ventricle with an average of 0.8 bacteria per injection. Fish harboring 1-3 bacteria were identified at 5 h post infection by confocal microscopy. These infected fish were then evaluated at 3 dpi and were scored as infected or uninfected, based on the presence or absence of fluorescent bacteria.

#### Fluorescence recovery after photobleaching (FRAP) experiments

FRAP experiments were performed using the galvo scanner (laser scanner) of the Nikon A1R confocal microscope with a 60x oil-immersion Plan Apo 1.4 NA objective. Photobleaching was performed with the 405 nm laser for 200ms on a region of interest (ROI) encompassing ∼1*μ*m from one pole of a single bacteria. A series of images was taken every second over the course of 31 seconds, one prior to bleaching. Labeled cells were mounted on a slide and coverslip in 0.75% low melting agarose. NIS Elements software (Nikon) was used to analyze the FRAP data to extract the fluorescence recovery kinetics. Briefly, the first image before photobleaching was used to generate an ROI for the entire cell and a second ROI was generated in the photobleached area. Total fluorescence intensities in both the whole cell area and the bleached area were extracted and normalized to correct for photobleaching of the dyes due to acquisition. The normalized fluorescence intensities of the bleached area were then fitted to a non-linear regression with a one-phase association, with the plateau values from each sample plotted to represent the mobile fraction.

#### Alkyne-cholesterol labeling of *M. marinum*

Wildtype or Δ*mmpL7 M. marinum* expressing wasabi fluorescent protein were grown in 7H9 medium supplemented with 10% OADC, 0.2% glycerol with or without 0.005% alkyne-cholesterol for 48 h. Bacteria were then washed 3x with PBS prior to the copper-click reaction with AlexaFluor-647 Azide. For copper click: 400µM BTTP (Click Chemistry Tools) and 200µM copper sulfate (Sigma) were dissolved in PBS and allowed to complex for 20 minutes. 30µM AlexaFluor-647 Azide (Thermo-Fisher) and 1.2mM sodium ascorbate (Sigma) were then added to the solution. Bacterial pellets in 96-well v-bottom plates were resuspended in 50µl of the solution and were incubated at 23 °C protected from light for 45 minutes. Bacteria were then washed 5x in PBS prior to imaging on a Nikon A1R confocal microscope with a 60x oil-immersion Plan Apo 1.4 NA objective. Nikon elements software was used to determine fluorescent intensities of wasabi and cholesterol signals calculated from line profiles drawn perpendicular to bacterial membranes at least 0.5µm from either pole.

#### Cholesterol depletion and infection of A549 epithelial cells

A549 cells were seeded at 50,000 cells per 8-well Nunc Lab-Tek II chambered coverglass or at 100,000 cells per well in a 24 well plate. Cells were incubated at 37 °C for 48 h. Cells were washed 1x in PBS followed by treatment with 10mM methyl-ß cyclodextrin (Sigma), 1mM water-soluble cholesterol (Sigma), or a combination of 10mM methyl-ß cyclodextrin and 1mM water-soluble cholesterol in serum free DMEM media. Cells were treated for 1 hour at 33 °C followed by 3 washes with PBS. Cells plated on chambered coverglass were infected with azido-DIM labeled *M. marinum* at an MOI of 5 for 24 h at 33 °C followed by imaging on a Nikon A1R confocal microscope with a 20x Plan Apo 0.75 NA objective. 2µm z-stacks were generated through the infected cells. Azido-DIM spreading was calculated similar to above (Section: *Confocal microscopy and image-based quantification of infection*). Cells plated on 24-well plates were rescued in DMEM + 10% FBS for 3 h at 33 °C. Cells were then harvested for quantification of cholesterol levels.

#### Atorvastatin treatment of zebrafish larvae

At 48 h post fertilization, 0.5µM atorvastatin (Sigma), or 10µM water-soluble cholesterol (Sigma), or both were added to zebrafish water containing 1% DMSO. Control fish were incubated in water with 1% DMSO only. Zebrafish were incubated for 24 h prior to infection or to cholesterol quantification. Drugs were replenished every 24 h until experiment endpoint.

#### Quantification of cholesterol in zebrafish and A549 epithelial cells

Cholesterol was quantified using the Total Cholesterol and Cholesterol Ester Colorimetric/Fluorometric Assay Kit (BioVision). For zebrafish, 8 larvae were euthanized, transferred to a 1.5ml Eppendorf tube and excess water was removed. A solution of chloroform:isopropanol:NP-40 (7:11:0.1) was added and the sample was sonicated in a water bath for 1 hour. Samples were then centrifuged at 16,000xg for 10min and supernatants were transferred to a fresh tube and were allowed to dry in a 60 °C water in a chemical fume hood. For A549 epithelial cells, chloroform:isopropanol:NP-40 (7:11:0.1) was added directly to cells in a 24 well plate. Cells were scrapped and solution was transferred to a 1.5ml Eppendorf tube and was centrifuged at 16,000xg for 10min and supernatants were transferred to a fresh tube and were allowed to dry in a 60 °C water in a chemical fume hood. Dried lipids were then subjected to manufacturer’s protocol and total cholesterol concentrations were determined by fluorescence on a SpectraMax i3x plate reader (Molecular Devices).

#### Statistics

Statistical analyses were performed using Prism 8.4.3 (GraphPad): When appropriate D’Agostino-Pearson normality test was done to determine if all of the groups in a particular data set were of a gaussian distribution which then guided the subsequent statistical test performed (Supplemental Table 1). Where the *n* value is given and not represented graphically in the figure, *n* represents the number of zebrafish used for each experimental group (Figure 7H and I).

## Supplemental Figures, Movie, and Table

**Supplemental Figure 1.**
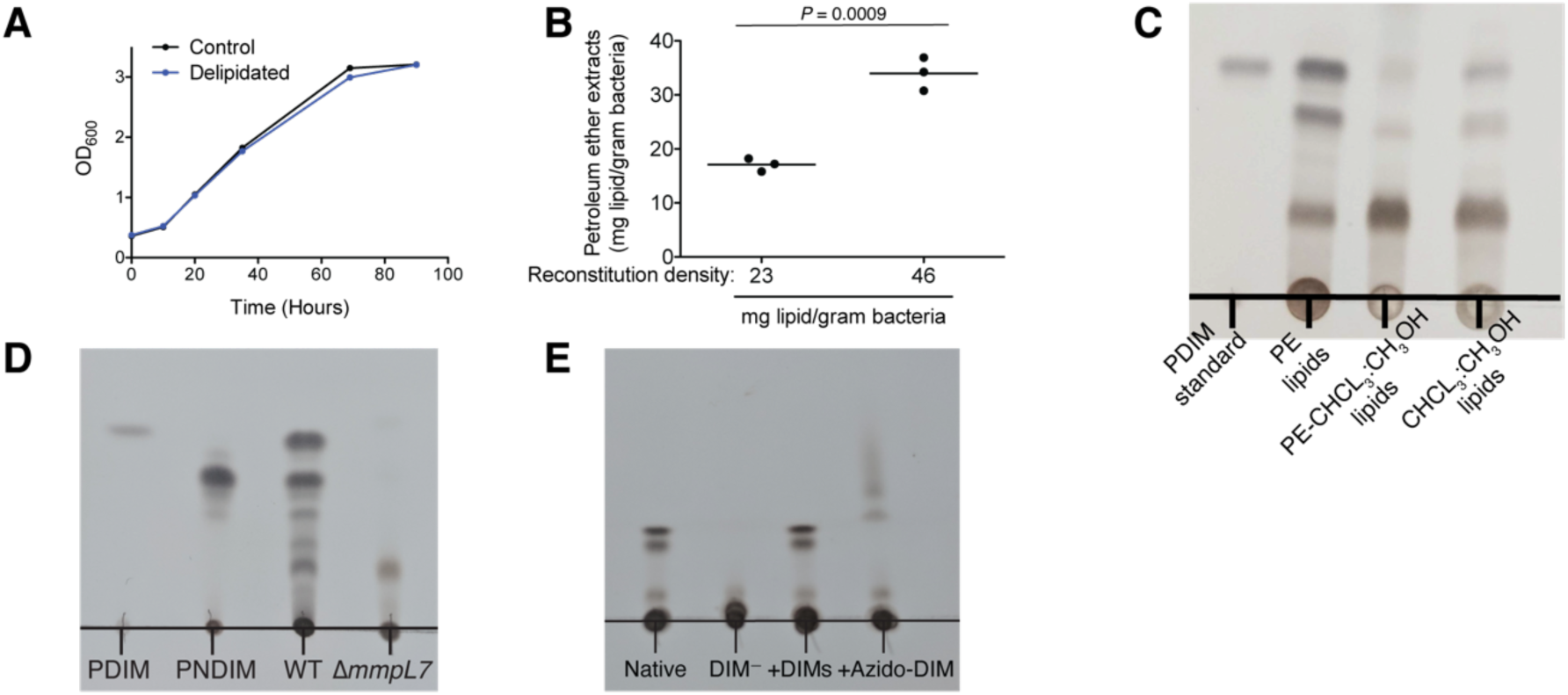
Optimization of petroleum ether extraction and reconstitution. **(A)** Mean OD_600_ values of control or delipidated *M. marinum* recovered in complete growth medium. **(B)** Mean mg of lipid extracted per gram of dry bacteria following reconstitution with designated lipid densities. Two-tailed, jnpaired t test. **(C)** Thin-layer chromatography (TLC) showing PDIM’s complete removal following petroleum ether extraction. PDIM standard, petroleum ether extract (PE lipids), chloroform:methanol extract of the eacterial pellet previously extracted with petroleum ether (PE-CHCL3:CH3OH lipids), and chloroform :metha-nol extract of a fresh bacterial pellet (CHCL3:CH3OH lipids). TLC was ran twice in 98:2 petroleum ethenethyl acetate. **(D)** TLC of PDIM standard, PNDIM standard, and petroleum ether extracts of wildtype (WT) and Δ*mmpL7 M. marinum*. TLC was ran twice in 98:2 petroleum etheriethyl acetate. **(E)** TLC of Native, DIM depleted (DIM^-^), DIM^-^ plus native DIMs (+DIMs), or DIM^-^ plus azido-DIM (+Azido-DIM) lipids prior to econstitution onto delipidated bacteria. TLC was ran once in 98:2 petroleum ether:ethyl acetate. **(A)-(E)** epresentative of three separate experiments.

**Supplemental Figure 2.**
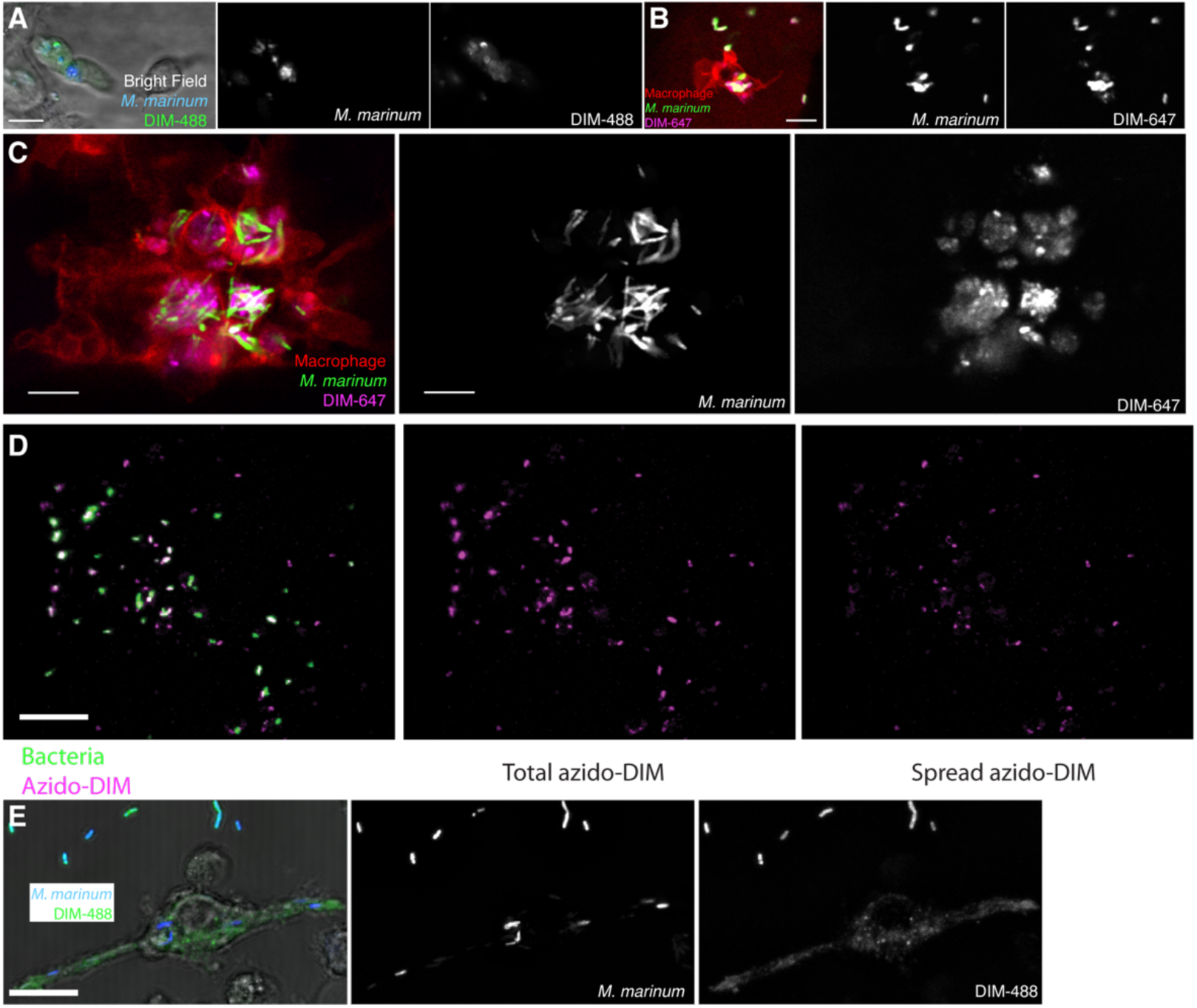
Further characterization of PDIM spread. **(A)** Image of *M. marinum* expressing a cytosolic blue fluorescent protein reconstituted with DIBO-488 (green) labeled azido-DIM at 3hpi of ∼100 *M. marinum* in the HBV of wildtype fish. Scale bar = 10pm. Images of M. marinum expresisng a cytosolic wasabi flourescent protein reconstituted with DIBO-647 labelled azido-DIM (DIM-647) at **(B)** 3 hours post infection and **(C)** 3 days post infection of ∼100 *M. marinum* in the HBV of transgenic fish whose macrophages express the red flourescent protein tdTomato. Scale bar = 10*μ*m. **(D)** Example of calculating percent spread: A surface rendering of the cytosolic expressing protein of the flourescent bacteria (green) is subtracted from the total azido-DIM signal (magenta) to give the spread azido-DIM signal. The volume of the spread azido-DIM signal is then divided by the volume of the total azido-DIM signal to calculate percent spread. Scale bar = 50*μ*m. **(E)** Image of *M. marinum* expresisng a blue fluorescent protein reconstituted with DIM-488 24 hpi of THP-1 macrophages, MOI = 5, and scale bar = 10*μ*m.

**Supplemental Figure 3.**
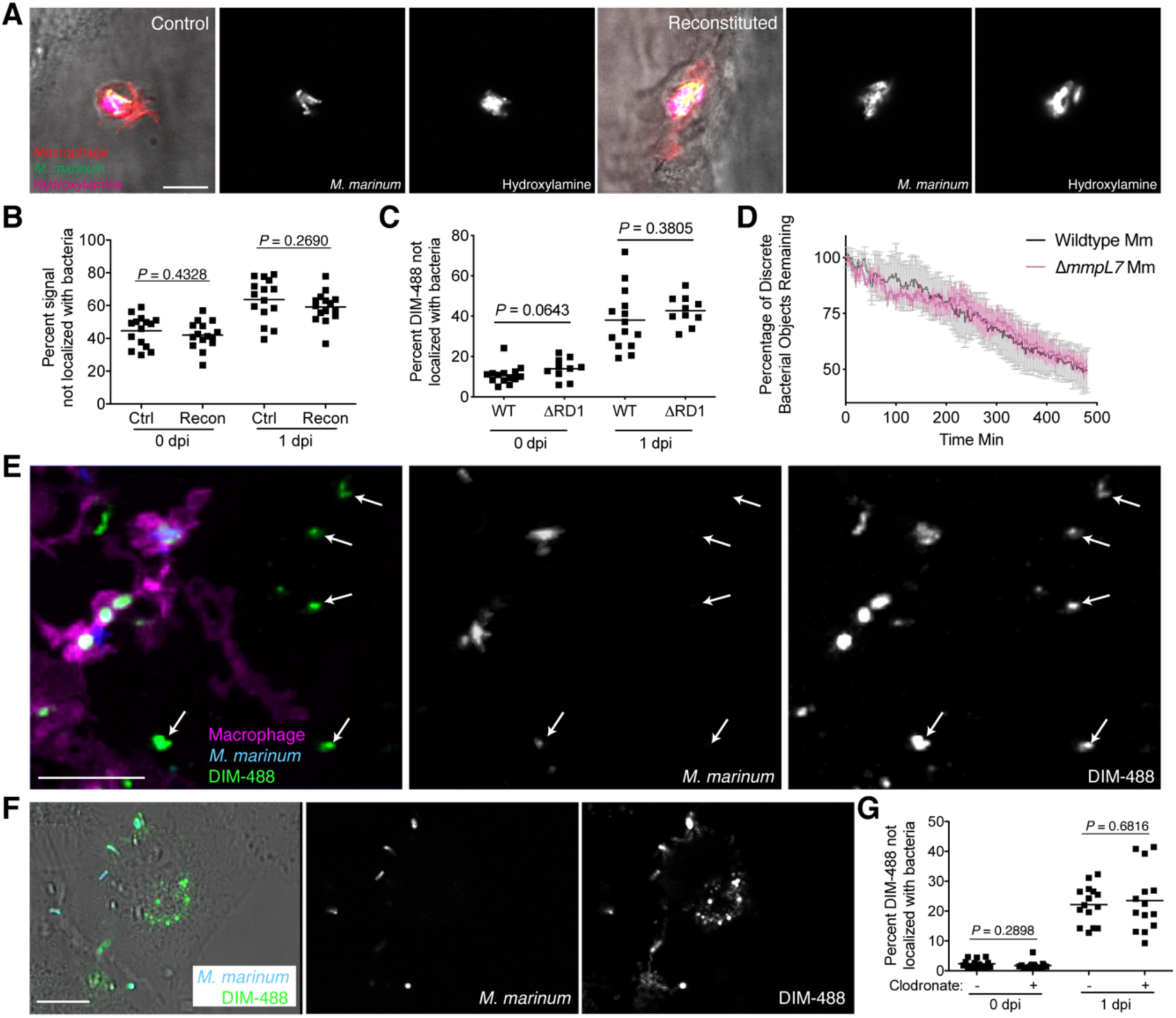
Functional evaluation of PDIM spread. **(A)** Images of control or reconstituted *M. marinum* expressing a cytosolic green fluorescent protein labeled with periodate-hydroxylamine chemistry at 3 hpi of ∼100 *M. marinum* into the HBV of transgenic fish whose macrophages express a fluorescent protein, scale bar = 10*μ*m. **(B)** Mean percent fluorescent signal not localized with bacteria following HBV infection of wildtype fish with ∼100 control or reconstituted *M. marinum* labeled with periodate-hydroxylamine chemistry. Two-tailed, unpaired t test. **(C)** Mean percent DIM-488 not localized with bacteria following HBV infection of wildtype fish with ∼100 wildtype or ARD1 *M. marinum*. Two-tailed Mann Whitney test for 0 dpi and two-tailed, unpaired t test for 1 dpi **(D)** Mean (+/- SEM) percentage of discrete bacterial objects remaining following HBV infection of wildtype fish with ∼100 wildtype or Δ*mmpL7 M. marinum*. Representative of two separate experiments. **(E)** Image of *M. marinum* expressing a cytosolic blue fluorescent protein reconstituted with DIBO-488 labeled azido-DIM (DIM-488) highlighting DIM-488 spread on epithelial cells (arrows) at 24 hpi of −100 *M. marinum* in the HBV of transgenic fish whose macrophages express a fluorescent protein, scale bar = 30*μ*m. **(F)** Image of *M. marinum* expressing a blue fluorescent protein reconstituted with DIM-488 24 hpi of A549 epithelial cells at an MOI of 5, scale bar = 10*μ*m. **(G)** Mean percent DIM-488 not localized with bacteria following HBV infection of lipo-PBS or lipo-clodronate treated fish with ∼100 *M. marinum*. Two-tailed Mann Whitney test for 0 dpi and two-tailed, unpaired t test for 1 dpi. **(B)**, **(C)**, and **(G)** representative of three separate experiments.

**Supplemental Figure 4.**
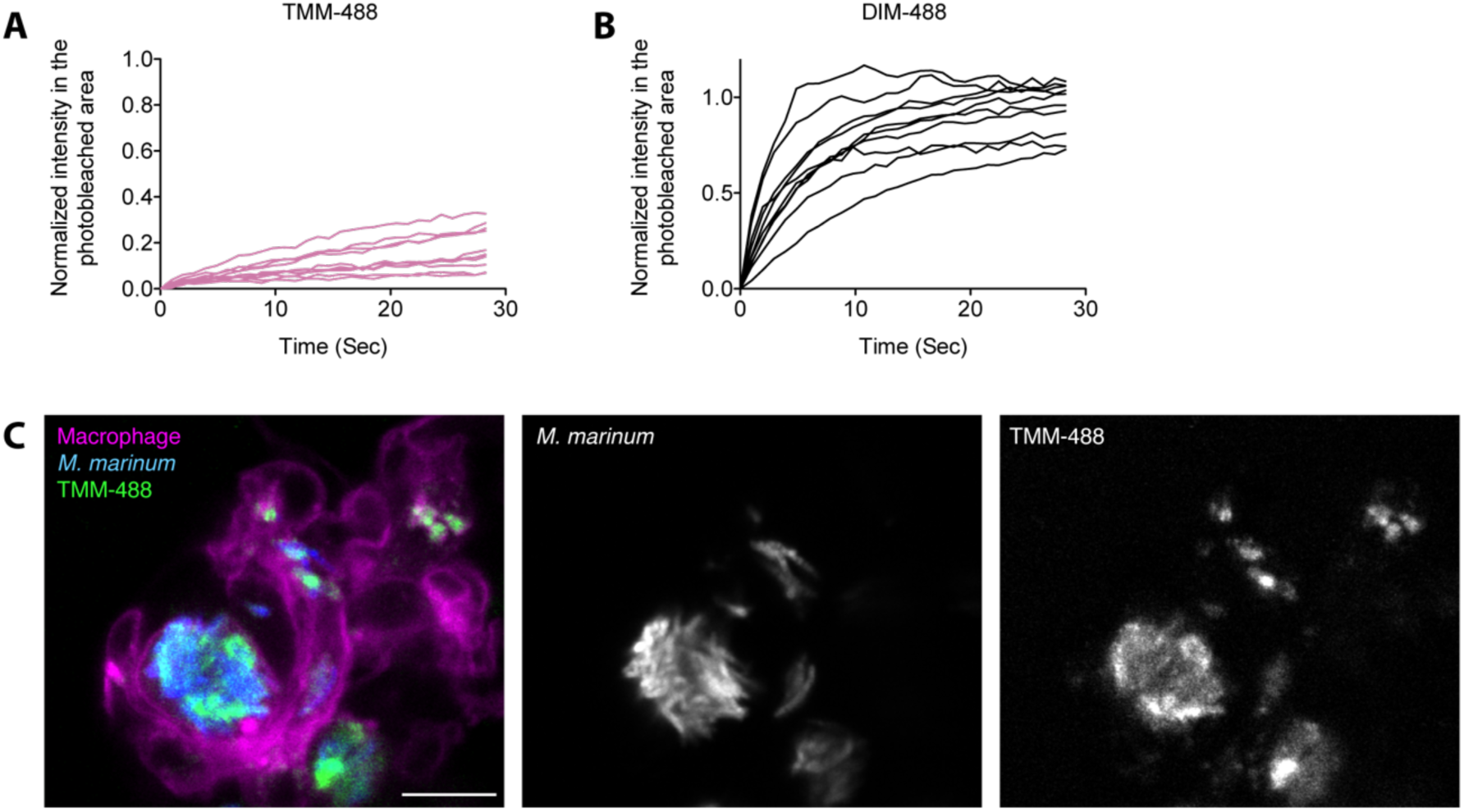
Individual FRAP curves of **(A)** TMM-488 and **(B)** DIM-488 labeled bacteria. **(C)** Image of *M. marinum* expressing a cytosolic blue fluorescent protein metabolically labeled with 6-azido trehalose followed by labeling with DIBO-488 (TMM-488) highlighting TMM-488 spread across macrophage membranes at 24 hpi of ∼100 *M. marinum* in the HBV of transgenic fish whose macrophages express a fluorescent protein.

**Supplemental Figure 5.**
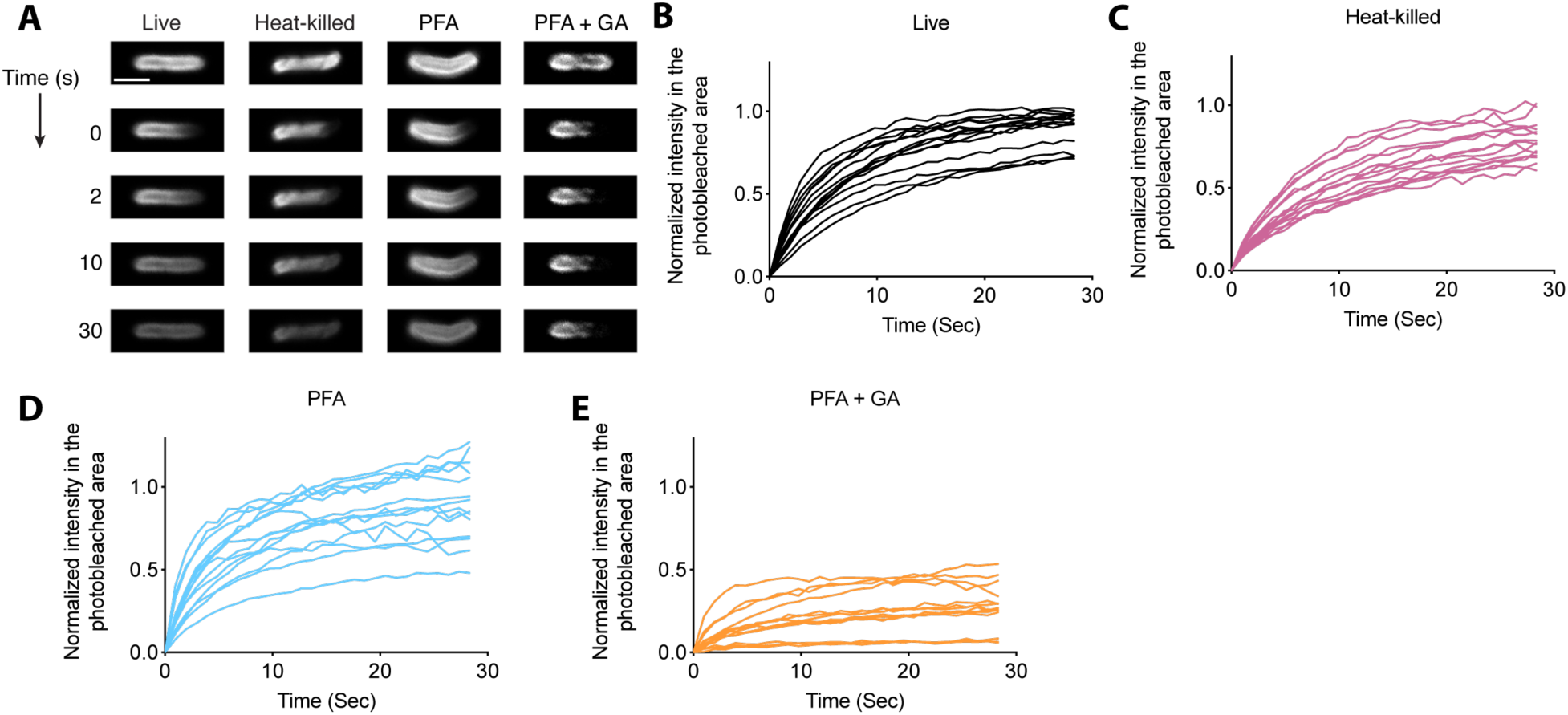
FRAP analysis of DIM-488 labeled bacteria. **(A)** Representative FRAP images of live, heat-killed. PFA, and PFA+GA treated DIM-488 labeled M. marinum, scale bar = 2*μ*m. **(B-E)** Individual fluorescent recovery curves of bacteria described in **A** Representative of three separate experiments.

**Supplementary Figure 6.**
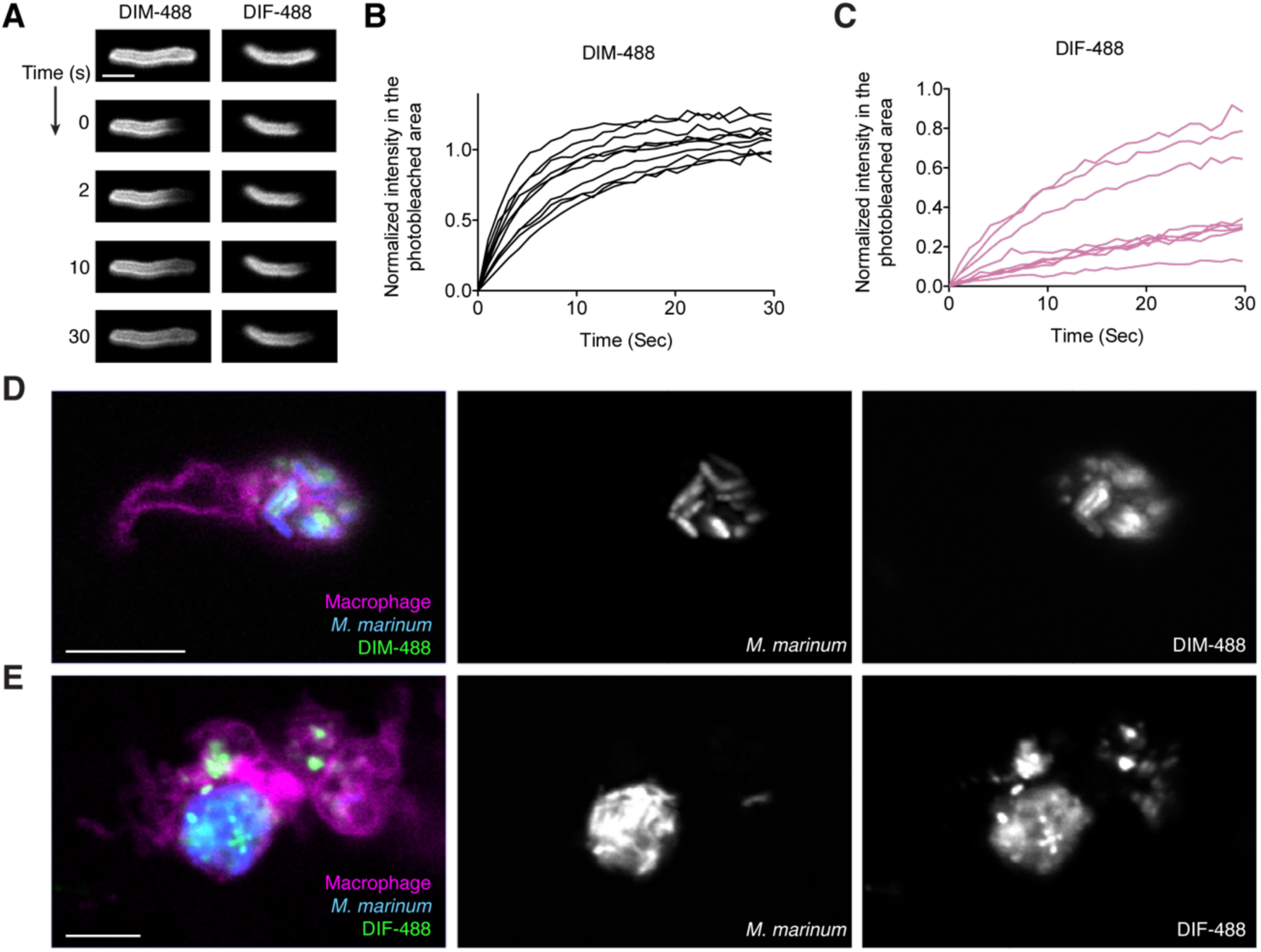
Analysis of DIM-488 and DIF-488 labeled *M. marinum*. **(A)** Representative FRAP images of DIM-488 and DIF-488 labeled *M. mannum*. scale bar = 2*μ*m Individual fluorescent recovery curves of **(B)** DIM-488 and **(C)** DIF-488 labeled *M. marinum* Images of M marinum expressing a cytosolic blue fluorescent protein reconstituted with green **(D)** DIM-488 or **(E)** DIF-488 at 24hpi of ∼100 *M. marinum* in the HBV of transgenic fish whose macrophages express a red fluorescent protein Scale bar = 10*μ*m. **(B)** and **(C)** represenatative of three separate experiments.

**Supplemental Figure 7.**
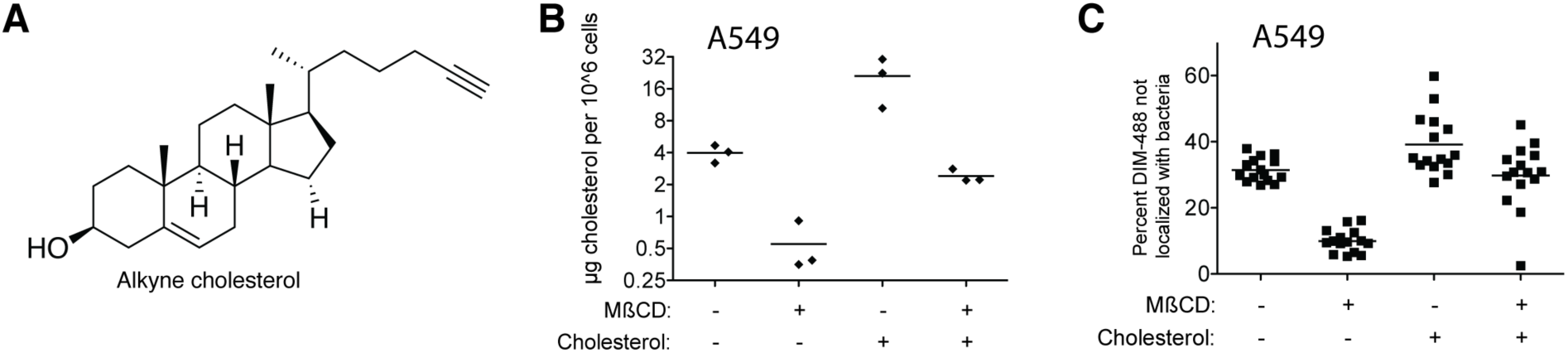
Cholesterol mediates DIM-488 spread on A549 epithelial cells. **(A)** Structure of alkyne cholesterol. **(B)** Mean cholesterol content of A549 cells treated with methyl-β cyclodextrin (MBCD), water-soluble cholesterol, or both. **(C)** Mean percent DIM-488 spread at 24 hpi of A549 epithelial cells treated with MßCD, water-soluble cholesterol, or both, MOI = 5. **(B)** and **(C)** representative of three separate experiments.

**Movie S1: PDIM dynamics**. Real-time video of *M. marinum* expressing blue fluorescent protein reconstituted with DIBO-488 labelled azido-DIM at 3 hpi of the HBV with ∼100 *M. marinum*.

**Supplementary Table 1**. Summary of *P* values and statistical tests. Gaussian distribution was determined using the D’Agostino-Pearson normality test. The result of this test guided subsequent analyses.

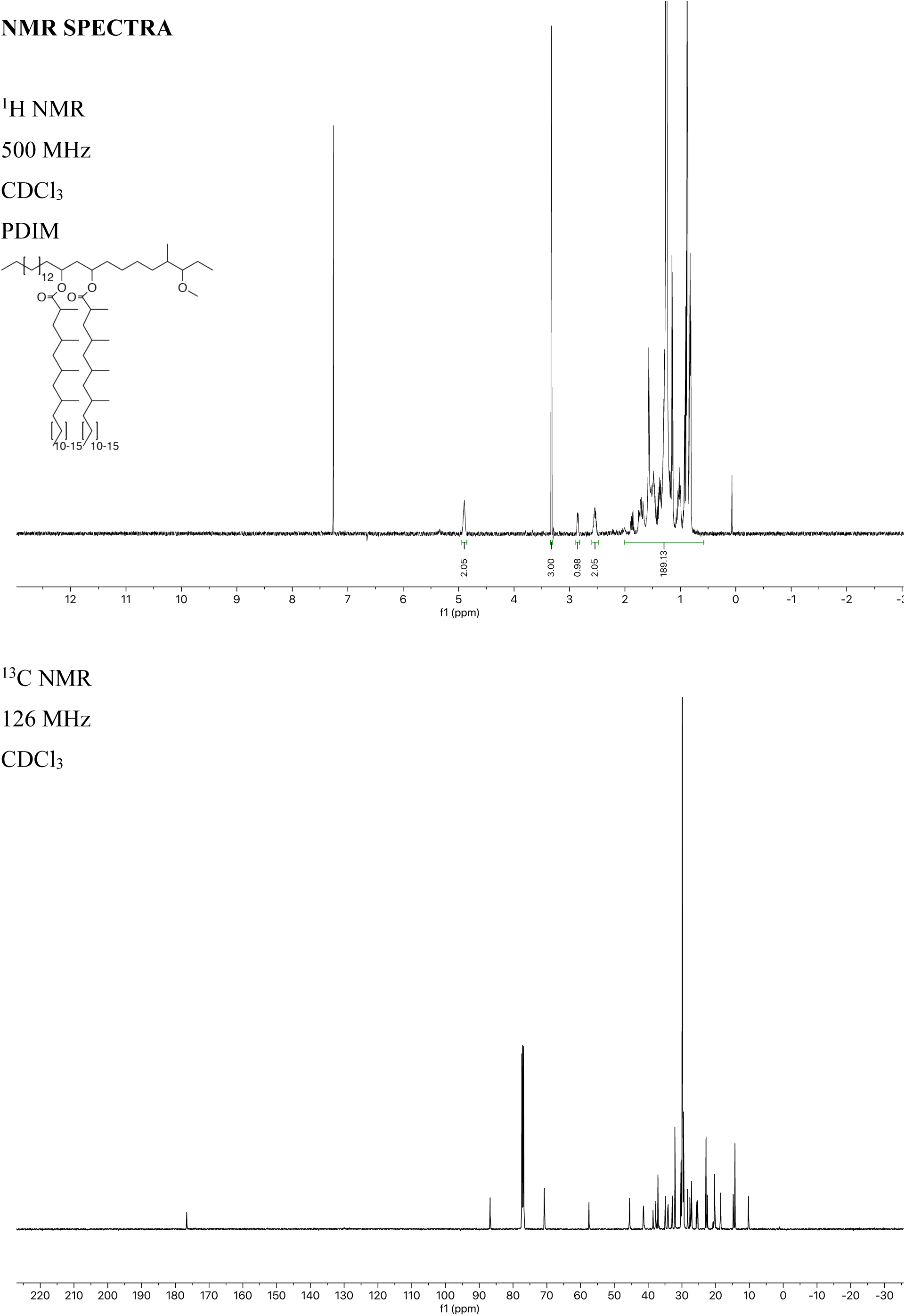

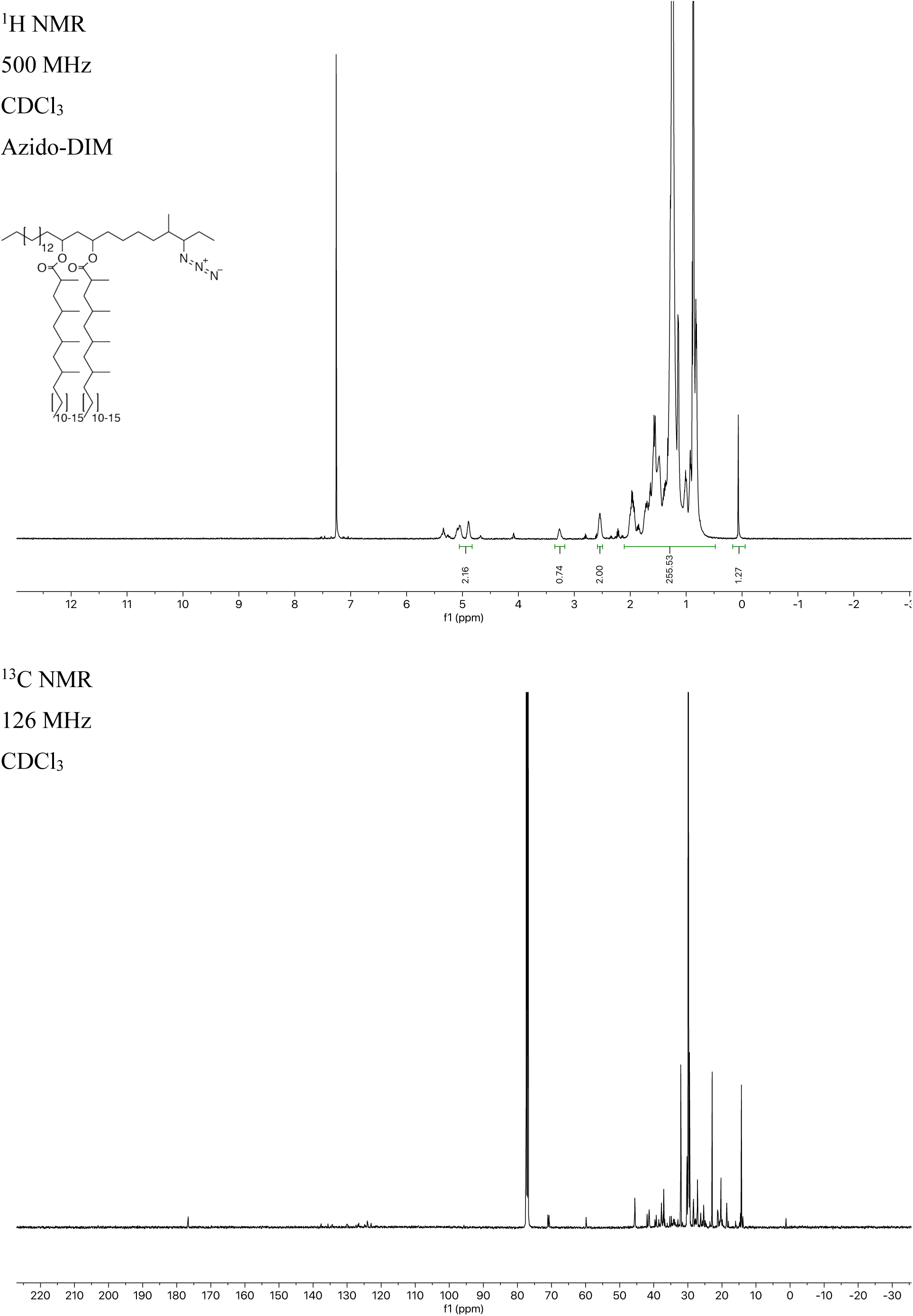

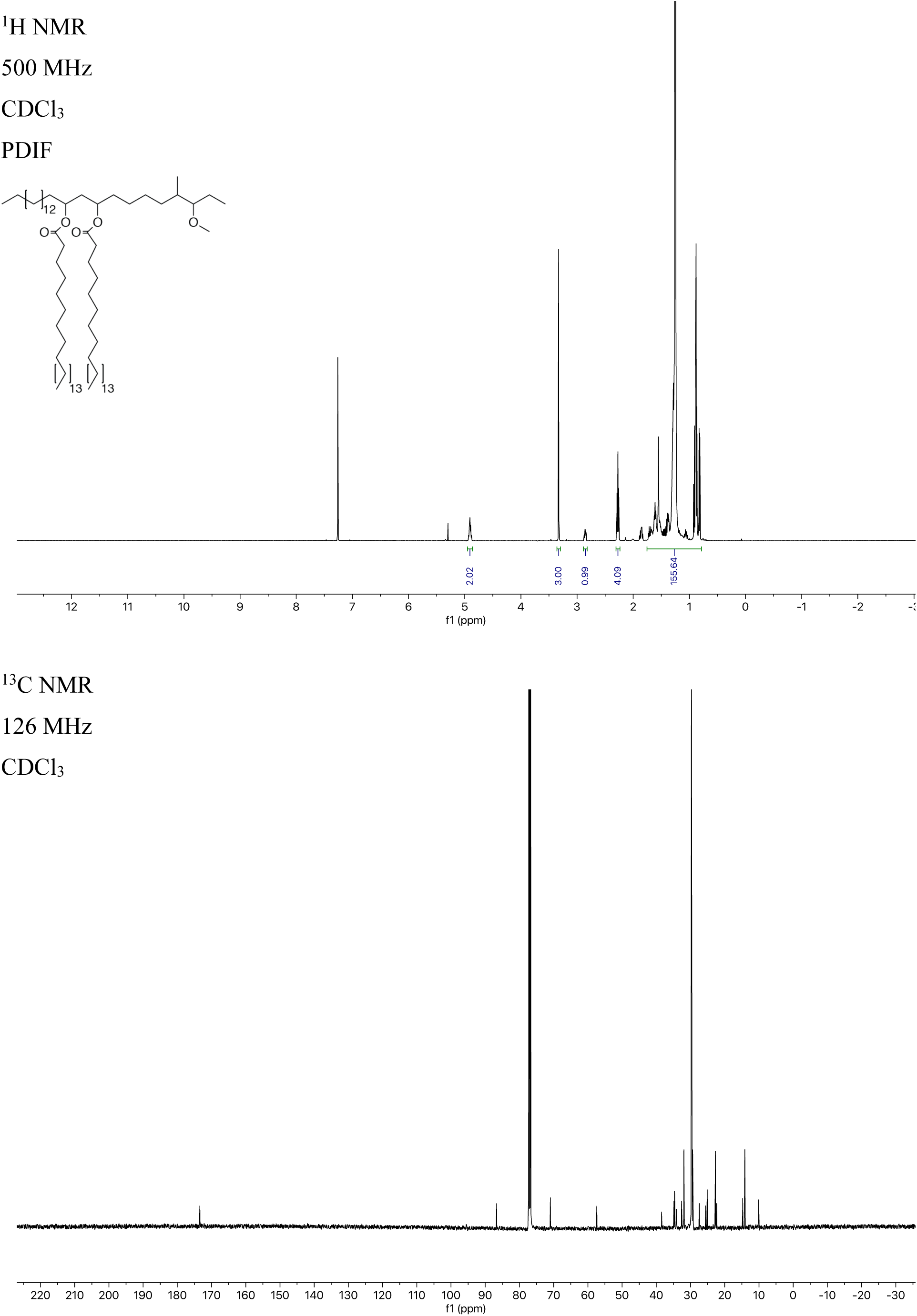

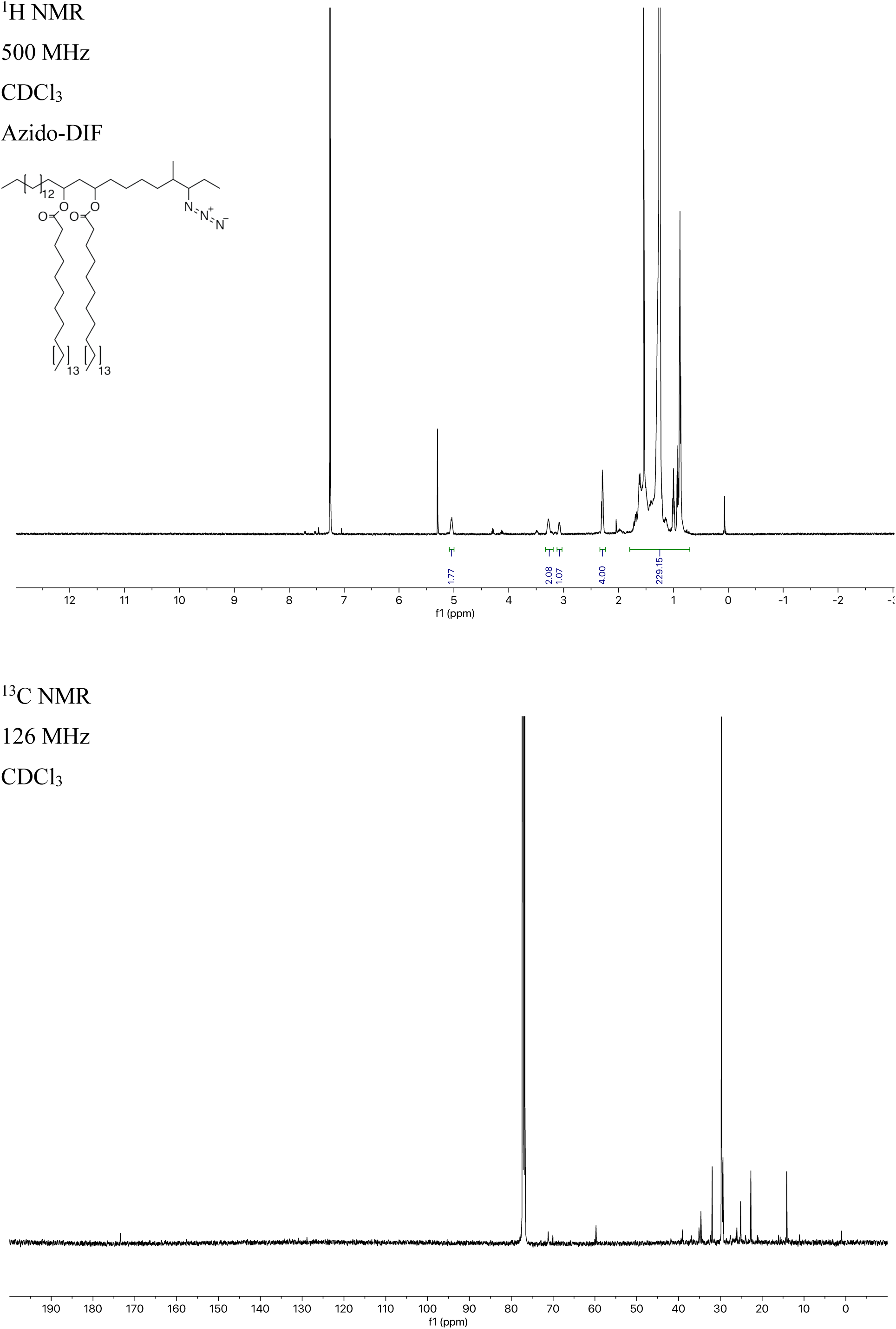

## REFERENCES

Anderson, R.J. (1929). The chemistry of the lipoids of tubercle bacilli iv. concerning the so called tubercle bacilli wax. analysis of the purified wax. J. Biol. Chem. 83, 505–522.

Astarie-Dequeker, C., Le Guyader, L., Malaga, W., Seaphanh, F.-K., Chalut, C., Lopez, A., and Guilhot, C. (2009). Phthiocerol Dimycocerosates of M. tuberculosis Participate in Macrophage Invasion by Inducing Changes in the Organization of Plasma Membrane Lipids. PLoS Pathog 5, e1000289–16.

Augenstreich, J., Arbues, A., Siméone, R., Haanappel, E., Wegener, A., Sayes, F., Le Chevalier, F., Chalut, C., Malaga, W., Guilhot, C., et al. (2017). ESX-1 and phthiocerol dimycocerosates of Mycobacterium tuberculosis act in concert to cause phagosomal rupture and host cell apoptosis. Cellular Microbiology 19.

Augenstreich, J., Haanappel, E., Ferré, G., Czaplicki, G., Jolibois, F., Destainville, N., Guilhot, C., Milon, A., Astarie-Dequeker, C., and Chavent, M. (2019). The conical shape of DIM lipids promotes Mycobacterium tuberculosis infection of macrophages. Proceedings of the National Academy of Sciences 3, 201910368.

Barczak, A.K., Avraham, R., Singh, S., Luo, S.S., Zhang, W.R., Bray, M.-A., Hinman, A.E., Thompson, M., Nietupski, R.M., Golas, A., et al. (2017). Systematic, multiparametric analysis of Mycobacterium tuberculosis intracellular infection offers insight into coordinated virulence. PLoS Pathog 13, e1006363–27.

Bates, J.H., Potts, W.E., and Lewis, M. (1965). Epidemiology of Primary Tuberculosis in an Industrial School. N. Engl. J. Med. 272, 714–717.

Bates, J.M., Akerlund, J., Mittge, E., and Guillemin, K. (2007). Intestinal alkaline phosphatase detoxifies lipopolysaccharide and prevents inflammation in zebrafish in response to the gut microbiota. Cell Host & Microbe 2, 371–382.

Beatty, W.L., Rhoades, E.R., Ullrich, H.J., Chatterjee, D., Heuser, J.E., and Russell, D.G. (2000). Trafficking and release of mycobacterial lipids from infected macrophages. Traffic 1, 235–247.

Bernut, A., Herrmann, J.-L., Kissa, K., Dubremetz, J.-F., Gaillard, J.-L., Lutfalla, G., and Kremer, L. (2014). Mycobacterium abscessus cording prevents phagocytosis and promotes abscess formation. Proceedings of the National Academy of Sciences 111, E943–E952.

Brzostek, A., Pawelczyk, J., Rumijowska-Galewicz, A., Dziadek, B., and Dziadek, J. (2009). Mycobacterium tuberculosis Is Able To Accumulate and Utilize Cholesterol. Journal of Bacteriology 191, 6584–6591.

Budin, I., de Rond, T., Chen, Y., Chan, L.J.G., Petzold, C.J., and Keasling, J.D. (2018). Viscous control of cellular respiration by membrane lipid composition. Science 362, 1186–1189.

Camacho, L.R., Constant, P., Raynaud, C., Lanéelle, M.-A., Triccas, J.A., Gicquel, B., Daffé, M., and Guilhot, C. (2001). Analysis of the Phthiocerol Dimycocerosate Locus of Mycobacterium tuberculosis. J. Biol. Chem. 276, 19845–19854.

Cambier, C.J., Falkow, S., and Ramakrishnan, L. (2014a). Host evasion and exploitation schemes of Mycobacterium tuberculosis. Cell 159, 1497–1509.

Cambier, C.J., O’Leary, S.M., O’Sullivan, M.P., Keane, J., and Ramakrishnan, L. (2017). Phenolic Glycolipid Facilitates Mycobacterial Escape from Microbicidal Tissue-Resident Macrophages. Immunity 47, 552–565.e554.

Cambier, C.J., Takaki, K.K., Larson, R.P., Hernandez, R.E., Tobin, D.M., Urdahl, K.B., Cosma, C.L., and Ramakrishnan, L. (2014b). Mycobacteria manipulate macrophage recruitment through coordinated use of membrane lipids. Nature 505, 218–222.

Cohen, S.B., Gern, B.H., Delahaye, J.L., Adams, K.N., Plumlee, C.R., Winkler, J.K., Sherman, D.R., Gerner, M.Y., and Urdahl, K.B. (2018). Alveolar Macrophages Provide an Early Mycobacterium tuberculosis Niche and Initiate Dissemination. Cell Host & Microbe 24, 439–446.e4.

Comas, I., Coscolla, M., Luo, T., Borrell, S., Holt, K.E., Kato-Maeda, M., Parkhill, J., Malla, B., Berg, S., Thwaites, G., et al. (2013). Out-of-Africa migration and Neolithic coexpansion of Mycobacterium tuberculosis with modern humans. Nat. Genet. 45, 1176–1182.

Cox, J.S., Chen, B., McNeil, M., and Jacobs, W.R. (1999). Complex lipid determines tissue-specific replication of Mycobacterium tuberculosis in mice. Nature 402, 79–83.

Davis, J.M., and Ramakrishnan, L. (2009). The role of the granuloma in expansion and dissemination of early tuberculous infection. Cell 136, 37–49.

Dutta, N.K., Bruiners, N., Pinn, M.L., Zimmerman, M.D., Prideaux, B., Dartois, V., Gennaro, M.L., and Karakousis, P.C. (2016). Statin adjunctive therapy shortens the duration of TB treatment in mice. J. Antimicrob. Chemother. 71, 1570–1577.

Falkow, S. (1988). Molecular Koch’s postulates applied to microbial pathogenicity. Rev. Infect. Dis. 10 Suppl 2, S274–S276.

Falkow, S. (2004). Molecular Koch’s postulates applied to bacterial pathogenicity--a personal recollection 15 years later. Nat. Rev. Microbiol. 2, 67–72.

Fessler, M.B. (2017). A New Frontier in Immunometabolism. Cholesterol in Lung Health and Disease. Ann Am Thorac Soc 14, S399–S405.

Gatfield, J., and Pieters, J. (2000). Essential role for cholesterol in entry of mycobacteria into macrophages. Science 288, 1647–1650.

Howell, J.I., Ahkong, Q.F., Cramp, F.C., Fisher, D., Tampion, W., and Lucy, J.A. (1972). Membrane fluidity and membrane fusion. Biochem. J. 130, 44P–44P.

Huebinger, J., Spindler, J., Holl, K.J., and Koos, B.X.R. (2018). Quantification of protein mobility and associated reshuffling of cytoplasm during chemical fixation. Scientific Reports 1– 11.

Indrigo, J. (2003). Cord factor trehalose 6,6’-dimycolate (TDM) mediates trafficking events during mycobacterial infection of murine macrophages. Microbiology 149, 2049–2059.

Jackson, M. (2014). The mycobacterial cell envelope-lipids. Cold Spring Harb Perspect Med 4.

Jung, M.E., and Lyster, M.A. (1977). Quantitative dealkylation of alkyl ethers via treatment with trimethylsilyl iodide. A new method for ether hydrolysis. J. Org. Chem. 42, 3761–3764.

Kamariza, M., Shieh, P., Ealand, C.S., Peters, J.S., Chu, B., Rodriguez-Rivera, F.P., Babu Sait, M.R., Treuren, W.V., Martinson, N., Kalscheuer, R., et al. (2018). Rapid detection of Mycobacterium tuberculosis in sputum with a solvatochromic trehalose probe. Sci Transl Med 10, eaam6310.

Karakousis, P.C., Chaisson, R.E., Martinson, N. (2019) StAT-TB (Statin Adjunctive Therapy for TB): A Phase 2b Dose-finding Study of Pravastatin in Adults with Tuberculosis. National Institute of Allergy and Infectious Diseases. https://clinicaltrials.gov/ct2/show/NCT03882177

Kim, M.-C., Yun, S.-C., Lee, S.-O., Choi, S.-H., Kim, Y.S., Woo, J.H., and Kim, S.-H. (2019). Association between Tuberculosis, Statin Use, and Diabetes: A Propensity Score–Matched Analysis. The American Journal of Tropical Medicine and Hygiene 101, 350–356.

Kondo, E., and Kanai, K. (1976). A suggested role of a host-parasite lipid complex in mycobacterial infection. Jpn. J. Med. Sci. Biol. 29, 199–210.

Lai, C.-C., Lee, M.-T.G., Lee, S.-H., Hsu, W.-T., Chang, S.-S., Chen, S.-C., and Lee, C.-C. (2016a). Statin treatment is associated with a decreased risk of active tuberculosis: an analysis of a nationally representative cohort. Thorax 71, 646–651.

Lai, C.-C., Lee, M.-T.G., Lee, S.-H., Hsu, W.-T., Chang, S.-S., Chen, S.-C., and Lee, C.-C. (2016b). Statin treatment is associated with a decreased risk of active tuberculosis: an analysis of a nationally representative cohort. Thorax 71, 646–651.

Lerner, T.R., Queval, C.J., Fearns, A., Repnik, U., Griffiths, G., and Gutierrez, M.G. (2017). Phthiocerol dimycocerosates promote access to the cytosol and intracellular burden of Mycobacterium tuberculosis in lymphatic endothelial cells. 1–13.

Los, D.A., and Murata, N. (2004). Membrane fluidity and its roles in the perception of environmental signals. Biochimica Et Biophysica Acta (BBA) - Biomembranes 1666, 142–157.

Maerz, L.D., Burkhalter, M.D., Schilpp, C., Wittekindt, O.H., Frick, M., and Philipp, M. (2019). Pharmacological cholesterol depletion disturbs ciliogenesis and ciliary function in developing zebrafish. Communications Biology 1–13.

Martens, G.W., Arikan, M.C., Lee, J., Ren, F., Vallerskog, T., and Kornfeld, H. (2008). Hypercholesterolemia Impairs Immunity to Tuberculosis. Infection and Immunity 76, 3464–3472.

Medzhitov, R. (2007). Recognition of microorganisms and activation of the immune response. Nature 449, 819–826.

Moliva, J.I., Hossfeld, A.P., Sidiki, S., Canan, C.H., Dwivedi, V., Beamer, G., Turner, J., and Torrelles, J.B. (2019). Selective delipidation of Mycobacterium bovis BCG enables direct pulmonary vaccination and enhances protection against Mycobacterium tuberculosis. Mucosal Immunology 12, 805–815.

Murry, J.P., Pandey, A.K., Sassetti, C.M., and Rubin, E.J. (2009). Phthiocerol dimycocerosate transport is required for resisting interferon-gamma-independent immunity. J Infect Dis 200, 774–782.

Nishijima, K., Yoneda, M., Hirai, T., Takakuwa, K., and Enomoto, T. (2019). Biology of the vernix caseosa: A review. J. Obstet. Gynaecol. Res. 45, 2145–2149.

Onwueme, K.C., Vos, C.J., Zurita, J., Ferreras, J.A., and Quadri, L.E.N. (2005). The dimycocerosate ester polyketide virulence factors of mycobacteria. Progress in Lipid Research 44, 259–302.

Osman, M.M., Pagán, A.J., Shanahan, J.K., and Ramakrishnan, L. (2020). Mycobacterium marinumphthiocerol dimycocerosates enhance macrophage phagosomal permeabilization and membrane damage. bioRxiv 1–20.

Pandey, A.K., and Sassetti, C.M. (2008). Mycobacterial persistence requires the utilization of host cholesterol. Proceedings of the National Academy of Sciences 105, 4376–4380.

Poger, D., Caron, B., and Mark, A.E. (2014). Effect of Methyl-Branched Fatty Acids on the Structure of Lipid Bilayers. J. Phys. Chem. B 118, 13838–13848.

Quigley, J., Hughitt, V.K., Velikovsky, C.A., Mariuzza, R.A., El-Sayed, N.M., and Briken, V. (2017). The Cell Wall Lipid PDIM Contributes to Phagosomal Escape and Host Cell Exit of Mycobacterium tuberculosis. mBio 8, e00148.#x2013;17–12.

Ramakrishnan, L. (2012). Revisiting the role of the granuloma in tuberculosis. Nat Rev Immunol 12, 352–366.

Ramakrishnan, L. (2020). Mycobacterium tuberculosis pathogenicity viewed through the lens of molecular Koch’s postulates. Curr. Opin. Microbiol. 54, 103–110.

Ran-Ressler, R.R., Khailova, L., Arganbright, K.M., Adkins-Rieck, C.K., Jouni, Z.E., Koren, O., Ley, R.E., Brenna, J.T., and Dvorak, B. (2011). Branched Chain Fatty Acids Reduce the Incidence of Necrotizing Enterocolitis and Alter Gastrointestinal Microbial Ecology in a Neonatal Rat Model. PLoS ONE 6, e29032–10.

Reuschl, A.-K., Edwards, M.R., Parker, R., Connell, D.W., Hoang, L., Halliday, A., Jarvis, H., Siddiqui, N., Wright, C., Bremang, S., et al. (2017). Innate activation of human primary epithelial cells broadens the host response to Mycobacterium tuberculosis in the airways. PLoS Pathog 13, e1006577–26.

Rodriguez-Rivera, F.P., Zhou, X., Theriot, J.A., and Bertozzi, C.R. (2017). Visualization of mycobacterial membrane dynamics in live cells. J. Am. Chem. Soc. 139, 3488–3495.

Rousseau, C., Winter, N., Pivert, E., Bordat, Y., Neyrolles, O., Avé, P., Huerre, M., Gicquel, B., and Jackson, M. (2004). Production of phthiocerol dimycocerosates protects Mycobacterium tuberculosis from the cidal activity of reactive nitrogen intermediates produced by macrophages and modulates the early immune response to infection. Cellular Microbiology 6, 277–287.

Ruysschaert, J.-M., and Lonez, C. (2015). Role of lipid microdomains in TLR-mediated signalling. BBA - Biomembranes 1848, 1860–1867.

Siegrist, M.S., Swarts, B.M., Fox, D.M., Lim, S.A., and Bertozzi, C.R. (2015). Illumination of growth, division and secretion by metabolic labeling of the bacterial cell surface. FEMS Microbiology Reviews 39, 184–202.

Silva, C.L., Ekizlerian, S.M., and Fazioli, R.A. (1985). Role of cord factor in the modulation of infection caused by mycobacteria. Am. J. Pathol. 118, 238–247.

Siméone, R., Constant, P., Malaga, W., Guilhot, C., Daffé, M., and Chalut, C. (2007). Molecular dissection of the biosynthetic relationship between phthiocerol and phthiodiolone dimycocerosates and their critical role in the virulence and permeability of Mycobacterium tuberculosis. FEBS Journal 274, 1957–1969.

Sletten, E.M., and Bertozzi, C.R. (2009). Bioorthogonal chemistry: fishing for selectivity in a sea of functionality. Angew. Chem. Int. Ed. Engl. 48, 6974–6998.

Soh, A.Z., Chee, C.B., Wang, Y.-T., Yuan, J.-M., and Koh, W.-P. (2016). Dietary Cholesterol Increases the Risk whereas PUFAs Reduce the Risk of Active Tuberculosis in Singapore Chinese. The Journal of Nutrition 146, 1093–1100.

Soong, G., Reddy, B., Sokol, S., Adamo, R., and Prince, A. (2004). TLR2 is mobilized into an apical lipid raft receptor complex to signal infection in airway epithelial cells. J. Clin. Invest. 113, 1482–1489.

Srivastava, S., Ernst, J.D., and Desvignes, L. (2014). Beyond macrophages: the diversity of mononuclear cells in tuberculosis. Immunol. Rev. 262, 179–192.

Swarts, B.M., Holsclaw, C.M., Jewett, J.C., Alber, M., Fox, D.M., Siegrist, M.S., Leary, J.A., Kalscheuer, R., and Bertozzi, C.R. (2012). Probing the mycobacterial trehalome with bioorthogonal chemistry. J. Am. Chem. Soc. 134, 16123–16126.

Takaki, K., Davis, J.M., Winglee, K., and Ramakrishnan, L. (2013). Evaluation of the pathogenesis and treatment of Mycobacterium marinum infection in zebrafish. Nat Protoc 8, 1114–1124.

Tall, A.R., and Yvan-Charvet, L. (2015). Cholesterol, inflammation and innate immunity. Nature Publishing Group 1–13.

Urbanek, B.L., Wing, D.C., Haislop, K.S., Hamel, C.J., Kalscheuer, R., Woodruff, P.J., and Swarts, B.M. (2014). Chemoenzymatic Synthesis of Trehalose Analogues: Rapid Access to Chemical Probes for Investigating Mycobacteria. Chembiochem 15, 2066–2070.

Urdahl, K.B. (2014). Understanding and overcoming the barriers to T cell-mediated immunity against tuberculosis. Seminars in Immunology 26, 578–587.

van der Wel, N., Hava, D., Houben, D., Fluitsma, D., van Zon, M., Pierson, J., Brenner, M., and Peters, P.J. (2007). M. tuberculosis and M. leprae translocate from the phagolysosome to the cytosol in myeloid cells. Cell 129, 1287–1298.

Volkman, H.E., Pozos, T.C., Zheng, J., Davis, J.M., Rawls, J.F., and Ramakrishnan, L. (2010). Tuberculous Granuloma Induction via Interaction of a Bacterial Secreted Protein with Host Epithelium. Science 327, 466–469.

Walton, E.M., Cronan, M.R., Beerman, R.W., and Tobin, D.M. (2015). The Macrophage-Specific Promoter mfap4 Allows Live, Long-Term Analysis of Macrophage Behavior during Mycobacterial Infection in Zebrafish. PLoS ONE 10, e0138949.

Wells, W.F., Ratcliffe, H.L., and Grumb, C. (1948). On the mechanics of droplet nuclei infection; quantitative experimental air-borne tuberculosis in rabbits. Am J Hyg 47, 11–28.

