## Supplemental Table 1 for "Spreading of a mycobacterial cell surface lipid into host epithelial membranes promotes infectivity"

### Supplementary Table 1

| | | | | | |
|---|---|---|---|---|---|
| Figure 1D | | | | | |
| Passed D'Agostino-Pearson normality test? Yes | | | | | |
| Ordinary One-Way ANOVA | | | | | |
| Sidak's multiple comparisons test | Mean Diff. | 95.00% CI of diff. | Significant? | Summary | Adjusted P Value |
| Control vs. Delipidated 1dpi | 2105 | -4602 to 8812 | No | ns | 0.8293 |
| Control vs. Reconstituted 1dpi | 3392 | -2986 to 9770 | No | ns | 0.4833 |
| Delipidated vs. Reconstituted 1dpi | 1287 | -4936 to 7510 | No | ns | 0.9427 |
| Sidak's multiple comparisons test | Mean Diff. | 95.00% CI of diff. | Significant? | Summary | Adjusted P Value |
| Control vs. Delipidated 3dpi | 13060 | 6353 to 19767 | Yes | **** | <0.0001 |
| Control vs. Reconstituted 3dpi | 2399 | -3979 to 8778 | No | ns | 0.7386 |
| Delipidated vs. Reconstituted 3dpi | -10660 | -16883 to -4437 | Yes | *** | 0.0002 |

| | | | | | |
|---|---|---|---|---|---|
| Figure 1F | | | | | |
| Passed D'Agostino-Pearson normality test? Yes | | | | | |
| Ordinary One-Way ANOVA | | | | | |
| Tukey's multiple comparisons test | Mean Diff. | 95.00% CI of diff. | Significant? | Summary | Adjusted P Value |
| wt ctrl vs wt + wt lipids | 14.55 | -4279 to 4308 | No | ns | >0.9999 |
| wt ctrl vs wt + ΔmmpL7 lipids | 8126 | 3469 to 12782 | Yes | **** | <0.0001 |
| wt ctrl vs ΔmmpL7 ctrl | 9328 | 4879 to 13776 | Yes | **** | <0.0001 |
| wt ctrl vs ΔmmpL7 + ΔmmpL7 lipids | 8607 | 3817 to 13397 | Yes | **** | <0.0001 |
| wt ctrl vs ΔmmpL7 + wt lipids | 3019 | -1347 to 7385 | No | ns | 0.3361 |
| wt + wt lipids vs wt + ΔmmpL7 lipids | 8111 | 3699 to 12523 | Yes | **** | <0.0001 |
| wt + wt lipids vs ΔmmpL7 ctrl | 9313 | 5121 to 13505 | Yes | **** | <0.0001 |
| wt + wt lipids vs ΔmmpL7 + ΔmmpL7 lipids | 8593 | 4040 to 13146 | Yes | **** | <0.0001 |
| wt + wt lipids vs ΔmmpL7 + wt lipids | 3004 | -1100 to 7109 | No | ns | 0.2752 |
| wt + ΔmmpL7 lipids vs ΔmmpL7 ctrl | 1202 | -3361 to 5765 | No | ns | 0.9709 |
| wt + ΔmmpL7 lipids vs ΔmmpL7 + ΔmmpL7 lipids | 481.5 | -4415 to 5378 | No | ns | 0.9997 |
| wt + ΔmmpL7 lipids vs ΔmmpL7 + wt lipids | -5107 | -9589 to -624.4 | Yes | * | 0.0165 |
| ΔmmpL7 ctrl vs ΔmmpL7 + ΔmmpL7 lipids | -720.4 | -5420 to 3979 | No | ns | 0.9976 |
| ΔmmpL7 ctrl vs ΔmmpL7 + wt lipids | -6309 | -10575 to -2043 | Yes | *** | 0.0007 |
| ΔmmpL7 + ΔmmpL7 lipids vs ΔmmpL7 + wt lipids | -5588 | -10209 to -967.2 | Yes | ** | 0.0091 |

| | | | | |
|---|---|---|---|---|
| Figure 1G | | | | |
| Passed D'Agostino-Pearson normality test? No | | | | |
| Kruskal-Wallis test | | | | |
| Dunn's multiple comparisons test | Mean rank diff. | Significant? | Summary | Adjusted P Value |
| MYD88(+) ctrl vs MYD88(-) ctrl | 50.65 | Yes | **** | <0.0001 |
| MYD88(+) ctrl vs MYD88(+) +ΔmmpL7 lipids | 5.7 | No | ns | >0.9999 |
| MYD88(+) ctrl vs MYD88(-) +ΔmmpL7 lipids | 58.78 | Yes | **** | <0.0001 |
| MYD88(+) ctrl vs MYD88(+) +wt lipids | 5.998 | No | ns | >0.9999 |
| MYD88(+) ctrl vs MYD88(-) +wt lipids | 12.94 | No | ns | >0.9999 |
| MYD88(-) ctrl vs MYD88(+) +ΔmmpL7 lipids | -44.95 | Yes | *** | 0.0002 |
| MYD88(-) ctrl vs MYD88(-) +ΔmmpL7 lipids | 8.131 | No | ns | >0.9999 |
| MYD88(-) ctrl vs MYD88(+) +wt lipids | -44.65 | Yes | *** | 0.0003 |
| MYD88(-) ctrl vs MYD88(-) +wt lipids | -37.71 | Yes | *** | 0.0007 |
| MYD88(+) +ΔmmpL7 lipids vs MYD88(-) +ΔmmpL7 lipids | 53.08 | Yes | **** | <0.0001 |
| MYD88(+) +ΔmmpL7 lipids vs MYD88(+) +wt lipids | 0.2976 | No | ns | >0.9999 |
| MYD88(+) +ΔmmpL7 lipids vs MYD88(-) +wt lipids | 7.242 | No | ns | >0.9999 |
| MYD88(-) +ΔmmpL7 lipids vs MYD88(+) +wt lipids | -52.79 | Yes | **** | <0.0001 |
| MYD88(-) +ΔmmpL7 lipids vs MYD88(-) +wt lipids | -45.84 | Yes | **** | <0.0001 |
| MYD88(+) +wt lipids vs MYD88(-) +wt lipids | 6.945 | No | ns | >0.9999 |

| | | | | |
|---|---|---|---|---|
| Figure 2F | | | | |
| Passed D'Agostino-Pearson normality test? No | | | | |
| Kruskal-Wallis test | | | | |
| Dunn's multiple comparisons test | Mean rank diff. | Significant? | Summary | Adjusted P Value |
| native vs. dim- | 18.75 | Yes | ** | 0.0078 |
| native vs. +dims | 3.167 | No | ns | >0.9999 |
| native vs. +AzDim | -7.038 | No | ns | >0.9999 |
| dim- vs. +dims | -15.58 | Yes | * | 0.0453 |
| dim- vs. +AzDim | -25.79 | Yes | **** | <0.0001 |
| +dims vs. +AzDim | -10.21 | No | ns | 0.4464 |

| | | | | |
|---|---|---|---|---|
| Figure 3C | | | | |
| Passed D'Agostino-Pearson normality test? No | | | | |
| Kruskal-Wallis test | | | | |
| Dunn's multiple comparisons test | Mean rank diff. | Significant? | Summary | Adjusted P Value |
| 0dpi vs. 1dpi | -19.47 | Yes | *** | 0.0009 |
| 0dpi vs. 3dpi | -36.42 | Yes | **** | <0.0001 |
| 1dpi vs. 3dpi | -16.95 | Yes | ** | 0.0049 |

| | |
|---|---|
| Figue 4C | |
| Passed D'Agostino-Pearson normality test? Yes | |
| Unpaired t test, DMM-488 vs TMM-488 | |
| P value | <0.0001 |
| P value summary | **** |
| Significantly different (P < 0.05)? | Yes |
| One- or two-tailed P value? | Two-tailed |
| t, df | t=7.711, df=17 |

| | |
|---|---|
| Figure 4D | |
| Passed D'Agostino-Pearson normality test? No | |
| Mann Whitney test, DIM-488 vs TMM-488 2hpi | |
| P value | <0.0001 |
| Exact or approximate P value? | Exact |
| P value summary | **** |
| Significantly different (P < 0.05)? | Yes |
| One- or two-tailed P value? | Two-tailed |
| Sum of ranks in column A,B | 484 , 146 |
| Mann-Whitney U | 10 |
| Mann Whitney test, DIM-488 vs TMM-488 24hpi | |
| P value | 0.9851 |
| Exact or approximate P value? | Exact |

| | |
|---|---|
| P value summary | ns |
| Significantly different (P < 0.05)? | No |
| One- or two-tailed P value? | Two-tailed |
| Sum of ranks in column C,D | 296 , 232 |
| Mann-Whitney U | 125 |

| | | | | | |
|---|---|---|---|---|---|
| Figure 4E | | | | | |
| Passed D'Agostino-Pearson normality test? Yes | | | | | |
| Ordinary One-Way ANOVA | | | | | |
| Tukey's multiple comparisons test | Mean Diff. | 95.00% CI of diff. | Significant? | Summary | Adjusted P Value |
| lipoPBS_kt27_azDIM vs. lipoCLOD_kt27_azDIM | 3.39 | -3.326 to 10.11 | No | ns | 0.5341 |
| lipoPBS_kt27_azDIM vs. lipoPBS_kt27_azTRE | 4.99 | -1.572 to 11.55 | No | ns | 0.1906 |
| lipoPBS_kt27_azDIM vs. lipoCLOD_kt27_azTRE | 21.94 | 15.38 to 28.50 | Yes | **** | <0.0001 |
| lipoCLOD_kt27_azDIM vs. lipoPBS_kt27_azTRE | 1.6 | -4.962 to 8.162 | No | ns | 0.9131 |
| lipoCLOD_kt27_azDIM vs. lipoCLOD_kt27_azTRE | 18.55 | 11.99 to 25.11 | Yes | **** | <0.0001 |
| lipoPBS_kt27_azTRE vs. lipoCLOD_kt27_azTRE | 16.95 | 10.55 to 23.35 | Yes | **** | <0.0001 |

| | | | | | |
|---|---|---|---|---|---|
| Figure 4G | | | | | |
| Passed D'Agostino-Pearson normality test? Yes | | | | | |
| Ordinary One-Way ANOVA | | | | | |
| Tukey's multiple comparisons test | Mean Diff. | 95.00% CI of diff. | Significant? | Summary | Adjusted P Value |
| Live vs. Heat-killed | 0.1015 | -0.03864 to 0.2417 | No | ns | 0.2319 |
| Live vs. PFA | 0.0258 | -0.1168 to 0.1684 | No | ns | 0.9633 |
| Live vs. PFA + GA | 0.6272 | 0.4845 to 0.7698 | Yes | **** | <0.0001 |
| Heat-killed vs. PFA | -0.07571 | -0.2183 to 0.06692 | No | ns | 0.5005 |
| Heat-killed vs. PFA + GA | 0.5257 | 0.3830 to 0.6683 | Yes | **** | <0.0001 |
| PFA vs. PFA + GA | 0.6014 | 0.4563 to 0.7464 | Yes | **** | <0.0001 |

| | | | | | |
|---|---|---|---|---|---|
| Figure 4H | | | | | |
| Passed D'Agostino-Pearson normality test? Yes | | | | | |
| Ordinary One-Way ANOVA | | | | | |
| Tukey's multiple comparisons test | Mean Diff. | 95.00% CI of diff. | Significant? | Summary | Adjusted P Value |
| live vs. pfa+ga | 16.97 | 8.863 to 25.07 | Yes | **** | <0.0001 |
| live vs. pfa | -0.03518 | -8.319 to 8.249 | No | ns | >0.9999 |
| live vs. heat-killed | 5.245 | -2.858 to 13.35 | No | ns | 0.3211 |
| pfa+ga vs. pfa | -17 | -25.28 to -8.716 | Yes | **** | <0.0001 |
| pfa+ga vs. heat-killed | -11.72 | -19.82 to -3.619 | Yes | ** | 0.002 |
| pfa vs. heat-killed | 5.28 | -3.004 to 13.56 | No | ns | 0.3346 |

| | |
|---|---|
| Figure 4J | |
| Passed D'Agostino-Pearson normality test? Yes | |
| Unpaired t test | |
| P value | <0.0001 |
| P value summary | **** |
| Significantly different (P < 0.05)? | Yes |
| One- or two-tailed P value? | Two-tailed |
| t, df | t=8.702, df=19 |

| | |
|---|---|
| Figure 4K | |
| Passed D'Agostino-Pearson normality test? Yes | |
| Unpaired t test | |
| P value | <0.0001 |
| P value summary | **** |
| Significantly different (P < 0.05)? | Yes |
| One- or two-tailed P value? | Two-tailed |
| t, df | t=11.42, df=28 |

| | |
|---|---|
| Figure 5C | |
| Passed D'Agostino-Pearson normality test? Yes | |
| Unpaired t test | |
| P value | 0.0004 |
| P value summary | *** |
| Significantly different (P < 0.05)? | Yes |
| One- or two-tailed P value? | Two-tailed |
| t, df | t=4.448, df=17 |

| | |
|---|---|
| Figure 5F | |
| Passed D'Agostino-Pearson normality test? No | |
| Mann Whitney test, DIM-488 vs DIF-488 2hpi | |
| P value | <0.0001 |
| Exact or approximate P value? | Exact |
| P value summary | **** |
| Significantly different (P < 0.05)? | Yes |
| One- or two-tailed P value? | Two-tailed |
| Sum of ranks in column A,B | 323 , 142 |
| Mann-Whitney U | 22 |
| Unpaired t test, DIM-488 vs DIF-488 24hpi | |
| P value | 0.0605 |
| P value summary | ns |
| Significantly different (P < 0.05)? | No |
| One- or two-tailed P value? | Two-tailed |
| t, df | t=1.962, df=26 |

| | | | | | |
|---|---|---|---|---|---|
| Figure 6A | | | | | |
| Passed D'Agostino-Pearson normality test? Yes | | | | | |
| Ordinary One-Way ANOVA | | | | | |
| Tukey's multiple comparisons test | Mean Diff. | 95.00% CI of diff. | Significant? | Summary | Adjusted P Value |
| WT Live vs. WT Heat-killed | 8 | 5.183 to 10.82 | Yes | **** | <0.0001 |
| WT Live vs. WT PFA | 8.143 | 5.326 to 10.96 | Yes | **** | <0.0001 |
| WT Live vs. WT PFA+GA | 0.9286 | -1.889 to 3.746 | No | ns | 0.9284 |
| WT Live vs. ∆mmpL7 Live | 1.019 | -1.751 to 3.789 | No | ns | 0.8902 |
| WT Live vs. ∆mmpL7 Heat-killed | 0.819 | -1.951 to 3.589 | No | ns | 0.954 |
| WT Heat-killed vs. WT PFA | 0.1429 | -2.674 to 2.960 | No | ns | >0.9999 |

| | | | | | |
|---|---|---|---|---|---|
| WT Heat-killed vs. WT PFA + GA | -7.071 | -9.889 to -4.254 | Yes | **** | <0.0001 |
| WT Heat-killed vs. ∆mmpL7 Live | -6.981 | -9.751 to -4.211 | Yes | **** | <0.0001 |
| WT Heat-killed vs. ∆mmpL7 Heat-killed | -7.181 | -9.951 to -4.411 | Yes | **** | <0.0001 |
| WT PFA vs. WT PFA + GA | -7.214 | -10.03 to -4.397 | Yes | **** | <0.0001 |
| WT PFA vs. ∆mmpL7 Live | -7.124 | -9.894 to -4.354 | Yes | **** | <0.0001 |
| WT PFA vs. ∆mmpL7 Heat-killed | -7.324 | -10.09 to -4.554 | Yes | **** | <0.0001 |
| WT PFA + GA vs. ∆mmpL7 Live | 0.09048 | -2.679 to 2.860 | No | ns | >0.9999 |
| WT PFA + GA vs. ∆mmpL7 Heat-killed | -0.1095 | -2.879 to 2.660 | No | ns | >0.9999 |
| ∆mmpL7 Live vs. ∆mmpL7 Heat-killed | -0.2 | -2.922 to 2.522 | No | ns | >0.9999 |

| | | | | | |
|---|---|---|---|---|---|
| Figure 6B | | | | | |
| Passed D'Agostino-Pearson normality test? Yes | | | | | |
| Ordinary One-Way ANOVA | | | | | |
| Tukey's multiple comparisons test | Mean Diff. | 95.00% CI of diff. | Significant? | Summary | Adjusted P Value |
| MYD88(+) live vs. MYD88(-) live | 0 | -2.579 to 2.579 | No | ns | >0.9999 |
| MYD88(+) live vs. MYD88(+) PFA+GA | -1.357 | -3.936 to 1.222 | No | ns | 0.5071 |
| MYD88(+) live vs. MYD88(-) PFA+GA | 3.857 | 1.278 to 6.436 | Yes | ** | 0.0012 |
| MYD88(-) live vs. MYD88(+) PFA+GA | -1.357 | -3.936 to 1.222 | No | ns | 0.5071 |
| MYD88(-) live vs. MYD88(-) PFA+GA | 3.857 | 1.278 to 6.436 | Yes | ** | 0.0012 |
| MYD88(+) PFA+GA vs. MYD88(-) PFA+GA | 5.214 | 2.635 to 7.793 | Yes | **** | <0.0001 |

| | | | | | |
|---|---|---|---|---|---|
| Figure 6C | | | | | |
| Passed D'Agostino-Pearson normality test? Yes | | | | | |
| Ordinary One-Way ANOVA | | | | | |
| Tukey's multiple comparisons test | Mean Diff. | 95.00% CI of diff. | Significant? | Summary | Adjusted P Value |
| Live vs. Heat-killed | 6320 | -10333 to 22973 | No | ns | 0.7361 |
| Live vs. PFA+GA | 16320 | 110.8 to 32529 | Yes | * | 0.0479 |
| Live vs. PFA | 2129 | -14524 to 18782 | No | ns | 0.9856 |
| Heat-killed vs. PFA+GA | 10000 | -6654 to 26653 | No | ns | 0.3803 |
| Heat-killed vs. PFA | -4191 | -21277 to 12894 | No | ns | 0.9104 |
| PFA+GA vs. PFA | -14191 | -30844 to 2462 | No | ns | 0.1177 |

| | | | | |
|---|---|---|---|---|
| Figure 6D | | | | |
| Passed D'Agostino-Pearson normality test? No | | | | |
| Kruskal-Wallis test | | | | |
| Dunn's multiple comparisons test | Mean rank diff. | Significant? | Summary | Adjusted P Value |
| MYD88 (+) live vs. MYD88(+) PFA+GA | 31.15 | Yes | **** | <0.0001 |
| MYD88(+) live vs. MYD88(-) live | -8.453 | No | ns | >0.9999 |
| MYD88(+) live vs. MYD88(-) PFA+GA | 0.2251 | No | ns | >0.9999 |
| MYD88(+) PFA+GA vs. MYD88(-) live | -39.6 | Yes | **** | <0.0001 |
| MYD88(+) PFA+GA vs. MYD88(-) PFA+GA | -30.92 | Yes | *** | 0.0001 |
| MYD88(-) live vs. MYD88(-) PFA+GA | 8.678 | No | ns | >0.9999 |

| | | | | |
|---|---|---|---|---|
| Figure 6E | | | | |
| Passed D'Agostino-Pearson normality test? No | | | | |
| Kruskal-Wallis test | | | | |
| Dunn's multiple comparisons test | Mean rank diff. | Significant? | Summary | Adjusted P Value |
| Native vs. DIM- | 45.31 | Yes | **** | <0.0001 |
| Native vs. +PDIM | 10.2 | No | ns | >0.9999 |
| Native vs. +PDIF | 57.88 | Yes | **** | <0.0001 |
| DIM- vs. +PDIM | -35.11 | Yes | *** | 0.0003 |
| DIM- vs. +PDIF | 12.57 | No | ns | 0.803 |
| +PDIM vs. +PDIF | 47.68 | Yes | **** | <0.0001 |

| | | | | | |
|---|---|---|---|---|---|
| Figure 7D | | | | | |
| Passed D'Agostino-Pearson normality test? n too small | | | | | |
| Tukey's multiple comparisons test | Mean Diff. | 95.00% CI of diff. | Significant? | Summary | Adjusted P Value |
| DMSO vs. Atorvastatin | 0.2015 | 0.06432 to 0.3387 | Yes | ** | 0.0067 |
| DMSO vs. Cholesterol | -0.1094 | -0.2466 to 0.02785 | No | ns | 0.1249 |
| DMSO vs. atorvastatin + cholesterol | -0.1154 | -0.2526 to 0.02182 | No | ns | 0.1026 |
| atorvastatin vs. cholesterol | -0.3109 | -0.4481 to -0.1737 | Yes | *** | 0.0004 |
| atorvastatin vs. atorvastatin + cholesterol | -0.3169 | -0.4541 to -0.1797 | Yes | *** | 0.0004 |
| cholesterol vs. atorvastatin + cholesterol | -0.006022 | -0.1432 to 0.1312 | No | ns | 0.9989 |

| | |
|---|---|
| Figure 7F | |
| Passed D'Agostino-Pearson normality test? Yes | |
| Unpaired t test | |
| P value | <0.0001 |
| P value summary | **** |
| Significantly different (P < 0.05)? | Yes |
| One- or two-tailed P value? | Two-tailed |
| t, df | t=5.052, df=33 |

| | |
|---|---|
| Figure 7G | |
| Passed D'Agostino-Pearson normality test? No | |
| Mann Whitney test, DMSO vs Atorvastatin 1dpi | |
| P value | 0.7624 |
| Exact or approximate P value? | Exact |
| P value summary | ns |
| Significantly different (P < 0.05)? | No |
| One- or two-tailed P value? | Two-tailed |
| Sum of ranks in column D,E | 1305 , 1323 |
| Mann-Whitney U | 620 |
| Passed D'Agostino-Pearson normality test? Yes | |
| Unpaired t test, DMSO vs Atorvastatin 3dpi | |
| P value | <0.0001 |
| P value summary | **** |
| Significantly different (P < 0.05)? | Yes |
| One- or two-tailed P value? | Two-tailed |
| t, df | t=4.563, df=70 |

| |
|---|
| Figure 7H |
| Fisher's exact test with Bonferroni's correction for multiple comparisons |

| | | | |
|---|---|---|---|
| Significance P < 0.05/(number of comparisons = 6) = 0.008 | | | |
| Comparison | P value | One- or two-sided | Statistically significant (P < 0.008)? |
| DMSO vs Atorvastatin | <0.0001 | Two-sided | Yes |
| DMSO vs Cholesterol | 0.6139 | Two-sided | No |
| DMSO vs Atrovastatin + Cholesterol | 0.6741 | Two-sided | No |
| Atorvastatin vs Cholesterol | <0.0001 | Two-sided | Yes |
| Atorvastatin vs Atorvastatin + Cholesterol | 0.0003 | Two-sided | Yes |
| Cholesterol vs Atorvastatin + Cholesterol | 0.357 | Two-sided | No |

| | | | |
|---|---|---|---|
| Figure 7I | | | |
| Fisher's exact test with Bonferroni's correction for multiple comparisons | | | |
| Significance P < 0.05/(number of comparisons = 28) = 0.0018 | | | |
| Comparison | P value | One- or two-sided | Statistically significant (P < 0.0018)? |
| MYD88(+)_DMSO WT vs MYD88(+)_Atorvastatin WT | <0.0001 | Two-sided | Yes |
| MYD88(+)_DMSO WT vs MYD88(-)_DMSO WT | >0.9999 | Two-sided | No |
| MYD88(+)_DMSO WT vs MYD88(-)_Atorvastatin WT | >0.9999 | Two-sided | No |
| MYD88(+)_DMSO WT vs MYD88(+)_DMSO ΔmmpL7 | <0.0001 | Two-sided | Yes |
| MYD88(+)_DMSO WT vs MYD88(+)_Atorvastatin ΔmmpL7 | <0.0001 | Two-sided | Yes |
| MYD88(+)_DMSO WT vs MYD88(-)_DMSO ΔmmpL7 | 0.3493 | Two-sided | No |
| MYD88(+)_DMSO WT vs MYD88(-)_Atorvastatin ΔmmpL7 | 0.5203 | Two-sided | No |
| MYD88(+)_Atorvastatin WT vs MYD88(-)_DMSO WT | <0.0001 | Two-sided | Yes |
| MYD88(+)_Atorvastatin WT vs MYD88(-)_Atorvastatin WT | 0.0001 | Two-sided | Yes |
| MYD88(+)_Atorvastatin WT vs MYD88(+)_DMSO ΔmmpL7 | 0.3805 | Two-sided | No |
| MYD88(+)_Atorvastatin WT vs MYD88(+)_Atorvastatin ΔmmpL7 | 0.8267 | Two-sided | No |
| MYD88(+)_Atorvastatin WT vs MYD88(-)_DMSO ΔmmpL7 | 0.0028 | Two-sided | No |
| MYD88(+)_Atorvastatin WT vs MYD88(-)_DMSO ΔmmpL7 | 0.0011 | Two-sided | Yes |
| MYD88(-)_DMSO WT vs MYD88(-)_Atorvastatin WT | >0.9999 | Two-sided | No |
| MYD88(-)_DMSO WT vs MYD88(+)_DMSO ΔmmpL7 | <0.0001 | Two-sided | Yes |
| MYD88(-)_DMSO WT vs MYD88(+)_Atorvastatin ΔmmpL7 | <0.0001 | Two-sided | Yes |
| MYD88(-)_DMSO WT vs MYD88(-)_DMSO ΔmmpL7 | 0.3503 | Two-sided | No |
| MYD88(-)_DMSO WT vs MYD88(-)_Atorvastatin ΔmmpL7 | 0.522 | Two-sided | No |
| MYD88(-)_Atorvastatin WT vs MYD88(+)_DMSO ΔmmpL7 | <0.0001 | Two-sided | Yes |
| MYD88(-)_Atorvastatin WT vs MYD88(+)_Atorvastatin ΔmmpL7 | <0.0001 | Two-sided | Yes |
| MYD88(-)_Atorvastatin WT vs MYD88(-)_DMSO ΔmmpL7 | 0.5468 | Two-sided | No |
| MYD88(-)_Atorvastatin WT vs MYD88(-)_Atorvastatin ΔmmpL7 | 0.757 | Two-sided | No |
| MYD88(+)_DMSO ΔmmpL7 vs MYD88(+)_Atorvastatin ΔmmpL7 | 0.658 | Two-sided | No |
| MYD88(+)_DMSO ΔmmpL7 vs MYD88(-)_DMSO ΔmmpL7 | 0.0001 | Two-sided | Yes |
| MYD88(+)_DMSO ΔmmpL7 vs MYD88(-)_Atorvastatin ΔmmpL7 | <0.0001 | Two-sided | Yes |
| MYD88(+)_Atorvastatin ΔmmpL7 vs MYD88(-)_DMSO ΔmmpL7 | 0.0006 | Two-sided | Yes |
| MYD88(+)_Atorvastatin ΔmmpL7 vs MYD88(-)_Atorvastatin ΔmmpL7 | 0.0002 | Two-sided | Yes |
| MYD88(-)_DMSO ΔmmpL7 vs MYD88(-)_Atorvastatin ΔmmpL7 | 0.7819 | Two-sided | No |

| | |
|---|---|
| Supplemental Figure 1B | |
| Passed D'Agostino-Pearson normality test? n too small | |
| Unpaired t test | |
| P value | 0.0009 |
| P value summary | *** |
| Significantly different (P < 0.05)? | Yes |
| One- or two-tailed P value? | Two-tailed |
| t, df | t=8.912, df=4 |

| | |
|---|---|
| Supplemental Figure 3B | |
| Passed D'Agostino-Pearson normality test? Yes | |
| Unpaired t test, Ctrl vs Recon 0dpi | |
| P value | 0.4328 |
| P value summary | ns |
| Significantly different (P < 0.05)? | No |
| One- or two-tailed P value? | Two-tailed |
| t, df | t=0.7959, df=28 |
| Unpaired t test, ctrl vs Recon 1dpi | |
| P value | 0.269 |
| P value summary | ns |
| Significantly different (P < 0.05)? | No |
| One- or two-tailed P value? | Two-tailed |
| t, df | t=1.128, df=28 |

| | |
|---|---|
| Supplemental Figure 3C | |
| Passed D'Agostino-Pearson normality test? No | |
| Mann Whitney test, WT vs ΔRD1 0dpi | |
| P value | 0.0643 |
| Exact or approximate P value? | Exact |
| P value summary | ns |
| Significantly different (P < 0.05)? | No |
| One- or two-tailed P value? | Two-tailed |
| Sum of ranks in column A,B | 143 , 157 |
| Mann-Whitney U | 38 |
| Passed D'Agostino-Pearson normality test? Yes | |
| Unpaired t test, WT vs ΔRD1 1dpi | |
| P value | 0.3805 |
| P value summary | ns |
| Significantly different (P < 0.05)? | No |
| One- or two-tailed P value? | Two-tailed |
| t, df | t=0.8950, df=22 |

| | |
|---|---|
| Supplemental FIgure 3G | |
| Passed D'Agostino-Pearson normality test? No | |
| Mann Whitney test, CLOD(-) vs CLOD(+) 0dpi | |
| P value | 0.2898 |
| Exact or approximate P value? | Exact |
| P value summary | ns |
| Significantly different (P < 0.05)? | No |
| One- or two-tailed P value? | Two-tailed |
| Sum of ranks in column A,B | 235 , 200 |

| | |
|---|---|
| Mann-Whitney U | 80 |
| Passed D'Agostino-Pearson normality test? Yes | |
| Unpaired t test, CLOD(-) vs CLOD(+) 1dpi | |
| P value | 0.6816 |
| P value summary | ns |
| Significantly different (P < 0.05)? | No |
| One- or two-tailed P value? | Two-tailed |
| t, df | t=0.4147, df=27 |

Supplementary Table 1: Summary of P values and statistical tests. Gaussian distribution was determined using the D'Agostino-Pearson normality test. The result of this test guided subsequent analyses.
